# Estimating abundance with interruptions in data collection using open population spatial capture-recapture models

**DOI:** 10.1101/671461

**Authors:** Cyril Milleret, Pierre Dupont, Joseph Chipperfield, Daniel Turek, Henrik Brøseth, Olivier Gimenez, Perry de Valpine, Richard Bischof

## Abstract

1. The estimation of population size remains one of the primary goals and challenges in ecology and provides a basis for debate and policy in wildlife management. Despite the development of efficient non-invasive sampling methods and robust statistical tools to estimate abundance, maintenance of field sampling is still subject to economic and logistic constraints. These can result in intentional or unintentional interruptions in sampling and cause gaps in data time series, posing a challenge to abundance estimation, and ultimately conservation and management decisions.
2. We applied an open population spatial capture-recapture (OPSCR) model to simulations and a real case study to test the reliability of abundance inferences models to interruption in data collection. Using individual detections occurring over consecutive sampling occasions, OPSCR models allow the estimation of abundance from individual detection data while accounting for lack of demographic and geographic closure between occasions. First, we simulated sampling data with interruptions in field sampling of different lengths and timing. We checked the performance of an OPSCR model in deriving abundance for species with slow and intermediate life history strategies. Finally, we introduced artificial sampling interruptions of various magnitudes and timing to a five-year non-invasive monitoring data set of wolverines (*Gulo gulo*) in Norway and quantified the consequences for OPSCR model predictions.
3. Inferences from OPSCR models were reliable even with temporal interruptions in monitoring. Interruption did not cause any systematic bias, but increased uncertainty. Interruptions occurring at occasions towards the beginning and the end of the sampling caused higher uncertainty. The loss in precision was more severe for species with a faster life history strategy.
4. We provide a reliable framework to estimate abundance even in the presence of sampling interruptions. OPSCR allows monitoring studies to provide contiguous abundance estimates to managers, stakeholders, and policy makers even when data are non-contiguous. OPSCR models do not only help cope with unintentional interruptions during sampling but also offer opportunities for using intentional sampling interruptions during the design of cost-effective population surveys.

## 1. Introduction

Estimating population size remains one of the most fundamental goals and challenges in wildlife ecology. Statistical tools that can account for imperfect detection, such as capture-recapture (CR) methods, are instrumental for estimating abundance of free-ranging populations (Seber 1982). Spatial-capture recapture (SCR) models, a recent extension of CR models, enable investigators to obtain spatially-explicit estimates of abundance (Efford 2004, Borchers and Efford 2008, Royle and Young 2008). SCR models estimate the location of individual activity centers (AC) using an observation model that describes the relationship between the spatial pattern of individual encounters and distance from the AC (i.e. detection probability). This allows SCR models to specify the spatial extent over which individuals occur and therefore generate spatially explicit estimates of abundance.

The SCR framework is suitable for analyzing observation data derived using not only physical capture and marking, but also non-invasive approaches, such as camera trapping (Efford et al. 2009, Royle et al. 2009), genetic sampling (Bischof et al. 2016a, Milleret et al. 2018), and acoustic sampling (Dawson and Efford 2009). Technical development in non-invasive methods have greatly expanded the spatial scope of monitoring and long-term studies. Many monitoring programs now collect individual detections with the aim of fitting SCR models. SCR models have, for example, been used to estimate density of brown bears (*Ursus arctos*) in Norway (Bischof et al. 2016a), of wolverines (*Gulo gulo*) in Alaska (Royle et al. 2011), and wolves (*Canis lupus*) in Spain (López-Bao et al. 2018). However, the maintenance of long-term data series, which is essential for establishing sound conservation and management plans (Lindenmayer and Likens 2009), can be subject to economic, logistic and other constraints. These can ultimately lead to intentional and unintentional interruption in sampling and thereby modify the temporal frequency of sampling (i.e. causing gaps in data time series).

When individual encounter data are collected over long periods relative to the lifespan of the study species, open population CR models can be used to account for the lack of demographic closure (i.e. death and emigration/ recruitment and immigration) between sampling occasions (i.e. generally referred to as primary occasions). Many monitoring projects are exposed to interruption in the sampling and result in gaps in CR time series.(e.g. Plummer 2003, Schmidt et al. 2007, Bears et al. 2009, Zabala et al. 2011, Zuberogoitia et al. 2016, Sanz–Aguilar et al. 2019). A gap causes unequal time intervals between primary sampling occasions. Unequal time intervals, are not a major problem in traditional CR models (Schmidt et al. 2007, Bears et al. 2009, Sanz–Aguilar et al. 2019), as it is possible to specify interval lengths when estimating demographic parameters such as survival or recruitment (Schmidt et al. 2007, Bears et al. 2009). However, when abundance estimates are the goal of the study, unequal time intervals in CR do not allow estimation of abundance during the primary period without data.. For example, the monitoring strategy for brown bears in Sweden is to conduct periodic sampling of different areas over multiple years (Kindberg et al. 2011, Swenson et al. 2017), which results in regions without detections available for inferences. Since information about annual population size is required by stakeholders, the current estimates are derived by combining periodic regional abundance estimates obtained with CR methods and an observation index collected on a yearly basis (Kindberg et al. 2011). Clearly, there is a need for methodology to cope with gaps in data time series.

Although no individual detections are available during the gap in the time-series, the Markovian structure of individual survival should help estimate the hidden state of the individual (i.e. dead or alive). Indeed, by modelling demographic processes (e.g. survival and recruitment) between primary occasions (e.g. years), the individual-based information is propagated across occasions. This means that the state of individuals at each sampling occasion (e.g. alive) can be reconstructed from the time series of detections (Figure 1). Therefore, open population CR models make effective use of the information obtained from multiple primary occasions compared to a series of independent CR models. Open population SCR models (OPSCR), which are a spatial extension of open population CR models, could offer practical solutions to deal with interruptions in sampling. OPSCR models do not only use information from individual detection collected during several occasions (such as CR models), but also use the spatial information contained in the detections and model movement of individuals between occasions (Ergon and Gardner 2014, Royle et al. 2014, Bischof et al. 2016a). In OPSCR, modelling individual movement between occasions allows estimating the probability of the individual being alive but outside of the study area, which should facilitate abundance estimates (Ergon and Gardner 2014, Gardner et al. 2018), especially during the gap years. The use of data collected over multiple occasions and propagation of individual information on spatial location and demographic status across time steps should help OPSCR bridge gaps in data collection, allowing inferences about abundance during occasions with sampling interruption. Although OPSCR models have already been used to infer abundance at occasions without individual detections (Chandler and Clark 2014, Augustine et al. 2019), there is a lack of knowledge about the quantitative consequences of sampling interruptions under different conditions (i.e. multiple interruptions, different life history characteristics).

**Figure 1.**
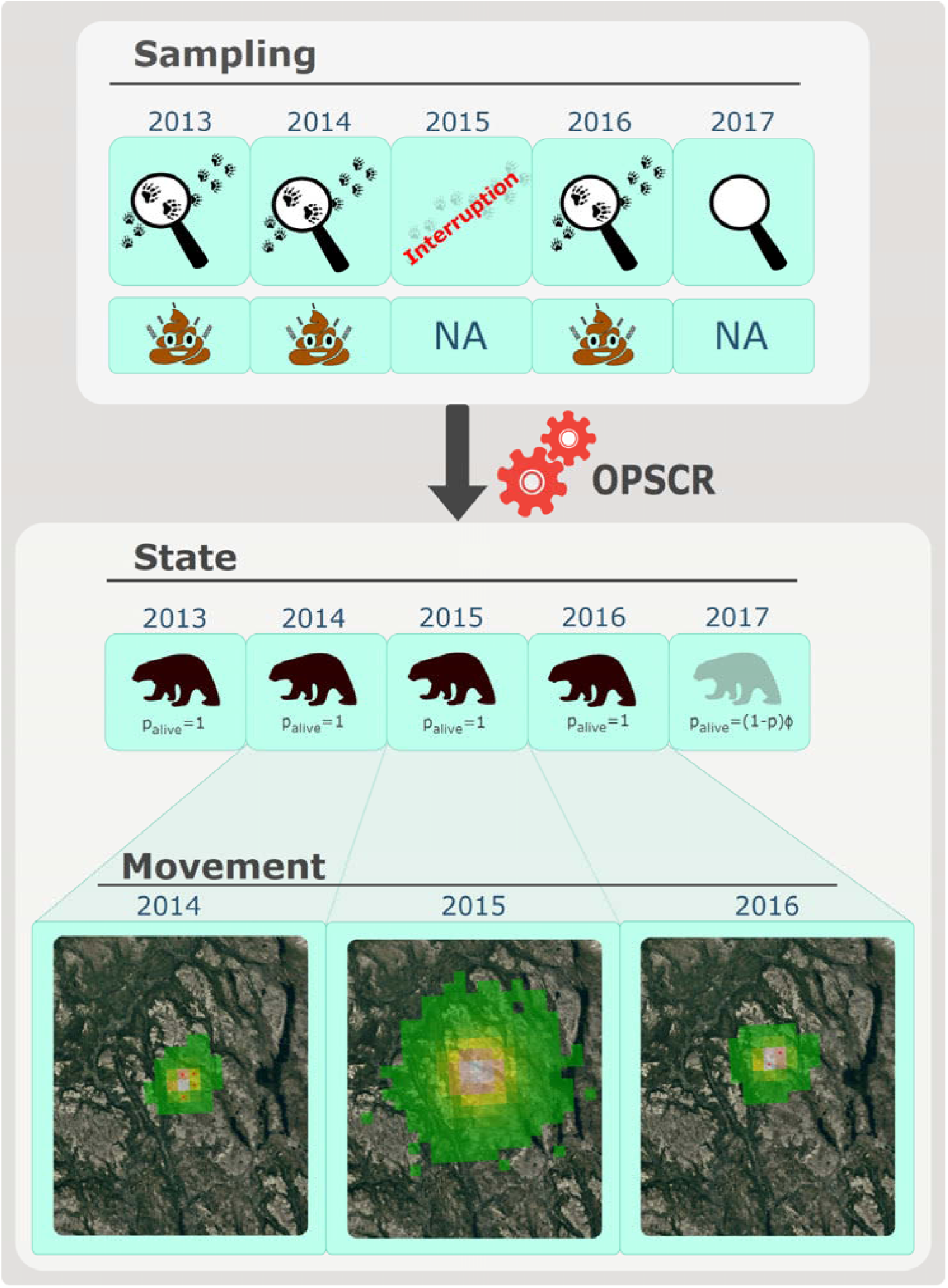
Illustration of the benefits of open population spatial capture recapture (OPSCR) models to estimate abundance when interruption in the sampling result in a gap in the data time series. The illustration is based on the detection history of one female wolverine during five winters (2013-2017) using scat-based non-invasive genetic monitoring in Norway. “Sampling” shows a timeline with a “scat emoji” at the occasion where the individual was detected and *NA* when not detected during the searches. In this illustration, we simulated a sampling interruption during the winter 2015 (i.e. all detections from all individuals were artificially removed during that occasion). “State” shows the individual state reconstruction during the interruption. When the individual was detected (2013, 2014, 2016), the individual was certain to be alive (black wolverine silhouette), as well as during the interruption (2015) because the individual was detected alive before and after the interruption. The probability of the individual being alive at all occasions between 2013 and 2016 (P_alive,_) equals to 1 from 2013 to 2016 (even for the occasion with interruption), because we could reconstruct with certainty the state of the individual to alive. At the last occasion, P_alive_ was estimated as : (1 − *p*) * *ϕ*, the probability for the individual to survive *ϕ* to the last occasion and not be detected (1 − *p*). “Movement” represents the movement process that models the individual’s activity center from one occasion to the other. The three maps (2014-2016) represent aerial photo of the study area, and green to grey colors show low to high probability of the AC being located in a given pixel, as predicted by the OPSCR model, respectively. During the interruption, the individual is certain to be alive, and the model uses population-level information about AC movement patterns to predict the most likely AC location of the individual. Individual detections are represented by red dots.

We built an OPSCR model to estimate abundance, recruitment, survival, and movement of individuals between primary sampling occasions. We then tested its reliability for inferring abundance in the presence of gaps in data collection when inferring abundance. We artificially generated sampling interruptions of various temporal configuration to assess their consequences for the precision and accuracy of abundance estimates. First, we introduced artificial sampling interruptions to simulated data sets for populations with different life history strategies (along the slow-fast continuum Stearns (1992)). Because of the low survival rate of species with a fast life history strategy, we expected sampling gaps to induce a more pronounced loss in precision compared with species with a slow life history. Most free-ranging populations are subject to demographic stochasticity in vital rates, which can be challenging to model in the presence of interruption. We therefore checked the effect of demographic stochasticity in vital rates on abundance estimates by simulating populations with and without temporal stochasticity in their vital rates. We then applied the OPSCR model to data from the non-invasive monitoring program of wolverines (*Gulo gulo*) in Norway as a real-life example, but with artificial gaps introduced. We provide recommendations for practitioners on how and under which conditions OPSCR can be used to obtain contiguous abundance estimates from non-contiguous monitoring data.

## 2. Material and methods

### 2.1. OPSCR model

We built a Bayesian OPSCR model that contained three main components: 1) an encounter model to estimate individual activity centers and account for imperfect detection of individuals (Royle et al. 2014), 2) a multi-state population dynamic model to estimate recruitment and survival (Seber 1965, Schwarz and Arnason 1996), and 3) a movement model to capture the movement of AC locations between years (Ergon and Gardner 2014). We used Markov Chain Monte Carlo (MCMC) and data augmentation to analyze OPSCR models and obtain estimates of abundance (Royle et al. 2007, 2009).

#### 2.1.1. The SCR model

The SCR model is the core element of our OPSCR model. SCR models use the spatial location of detections and non-detections at a set of detectors to estimate the latent locations of individual activity centers (ACs). SCR models are hierarchical state-space models combining 1) a point process model that describes the spatial distribution of individual ACs, and 2) a detection model conditional on the point process model, which describes the relationship between individual detection probability and distance to its AC. The half-normal detection model commonly used in SCR assumes that the probability *p* of detecting individual *i* at detector *j* and time *t* decreases with distance between the detector and the AC (*D*_*ijt*_):

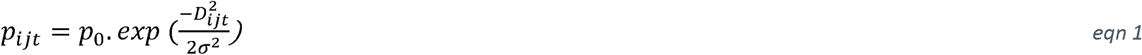

where p_0_ represents the detection probability at the location of the AC, and *σ* represents the width of the utilization distribution. The scale parameter *σ* is related to the extent of space used over the period of study.

#### 2.1.2. The multistate model

Individual state membership *z*_*it*_ takes the value 1 if “not yet entered”, 2 if “alive”, and 3 if “dead”. State *z* is the result of a Markovian process and changes with time according to a categorical distribution (Gimenez et al. 2007, Kery and Schaub 2011). During the first occasion, individuals can only be designated as “not yet entered” or “alive” so that *z*_*i*1_∼*dcat*(1 − *ψ, ψ*, 0), where *ψ* represents the inclusion probability.

For *t≥2, z*_*it*_ is conditional on the state of individual *i* at *t-1:*

- If *z*_*it*−1_ = 1, individual *i* is potentially available to be recruited (transition to state 2), so *z*_*it*_ ∼*dcat*(1 − *γ*_*t*_, *γ*_*t*_, 0), where *γ*_*t*_ is the recruitment parameter and is derived as:

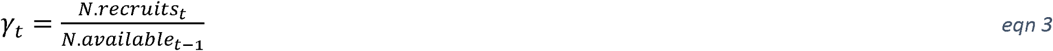

where *N. available* represents the number of augmented individuals with the state *not yet entered* (i.e. individuals available for transitioning to the *alive* state at each occasion), and *N. recruits* is the number of new individuals recruited into the population:

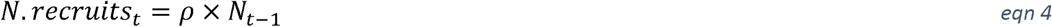

where *ρ* is the per capita recruitment parameter:

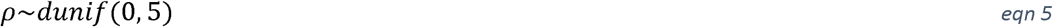
- If *z*_*it*−1_ = 2 individual *i* can either survive and remain *z*_*it*_ = 2 or die and transition to *z*_*it*_ = 3, so that *z*_*it*−1_∼*dcat*(0, *ϕ*, 1 − *ϕ*), where *ϕ* represent the survival probability.
- If *z*_*it*−1_ = 3, individual *i* is dead and will remain in this (absorbent) state.

Only individuals with the state “*alive*” can be detected. We therefore linked the encounter indicator *y* (detected, not detected) of individual *i* at detector *j* and time *t* with the individual’s state *z*_[*i,t*_]:

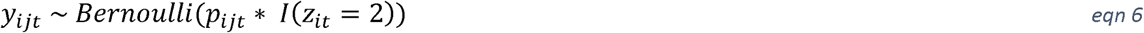

where *I* is an indicator function returning 1 for individuals in state 2, and 0 for individuals in state 1 or 3.

Estimates of abundance 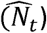 were obtained as:

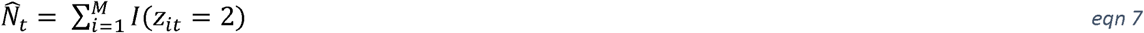

The state z_it_ of an individual is a latent variable, except at occasions when the individual was detected alive where it can be set to “alive”. In certain cases, it is also possible to reconstruct with certainty the state of individuals at occasion during which they were not detected (Figure 1). For example, an individual is known to be alive in years in which it was not detected, if that period is framed by alive detections.

#### 2.1.3. The Movement model

ACs at *t* = 1 were placed according to a homogenous Binomial point process (Illian et al. 2008). Under this model, AC positions were independently and uniformly distributed in the study area (S). In order to distinguish between temporary emigration and mortality, we integrated a movement model in the OPSCR model allowing shifts of individual activity centers between occasions. This is an important component of the OPSCR model as it can improve survival estimates and can take into account the impact of animals moving within and out of the sampled area (Ergon and Gardner 2014, Gardner et al. 2018). It is a particularly important feature of the model in the context of sampling interruption, as it helps propagating spatial locations of individual across occasions. Movement was modelled as a Markovian spatial point process. The outcome of each movement event was placed according to an inhomogeneous binomial point process (Illian et al. 2008) with only a single point (AC) simulated for each movement event. The functional form of the intensity surface that determined the location of the AC placement was a combination of an isotropic multivariate normal distribution centered around the source coordinates (location of the AC at previous occasion) with a standard deviation *τ*, and an intensity surface representing habitat quality. For simplicity, we considered homogenous habitat quality in this study (see Supporting information 1.1).

### 2.2. Simulations

We conducted a simulation study to evaluate the performance of our model under sampling interruptions of different magnitudes and configurations. We created a spatial domain of 40 × 40 distance units (du) within which we centered a 20du ×20du detector grid (with a minimum distance of 1.5 du between detectors). We released 50 individuals (N_1_) in the first occasion and sampled the location of their ACs uniformly within the spatial domain. During the subsequent occasions, we simulated individual movements as Markovian spatial point processes with the intensity surface being a multivariate normal distribution centered on the previous AC location. We simulated population dynamics assuming that the sampling occasion occurred just prior to reproduction. We drew the number of recruits (*ρ*) for each alive individual from a Poisson distribution. Note that if the sampling period does not start exactly after birth, *ρ* is a composite parameter of the number of offspring produced by an individual and their survival rate until the start of the sampling. Each alive individual had a probability *ϕ* to survive to the next sampling occasion.

#### 2.2.1. Population and survey characteristics

We simulated individual detections occurring at five consecutive primary occasions (e.g. for simplicity, we considered a one-year time interval between instantaneous occasions) using σ=2 and *p0=* 0.25, which led to an overall occasion-specific detectability of 71.2% (SD=6.46). We used a multivariate normal distribution with *τ* = 3 for the movement of ACs between occasions. We considered two stable populations (asymptotic growth rate=1) having contrasting life history characteristics along the slow-fast continuum (Stearns 1992) (Table 1). We simulated populations having a “slow” and “intermediate” life history strategy with *ϕ*=0.85 and ρ=0.15, and *ϕ*=0.65 and ρ=0.35 (Table 1), respectively. We did not consider a population having a faster life history strategy because the relative life span of individuals would be too short compared to the time interval between two consecutive occasions (a year). In addition of the stochastic realization of *z*_*it*_, we also considered scenarios with larger temporal stochasticity by drawing *ϕ*_*t*_ and *ρ*_*t*_ on a logit link from a normal distribution centered on the average values of the respective life history strategy and SD=0.2.

**Table 1.** Characteristics of the four simulated populations used to assess the consequences of sampling interruption on abundance estimates from open population spatial capture recapture models. Median survival time is expressed in years. Super population size represents the average number of individuals (from all simulated data sets) that were ever alive during the study. ρ and ψ are the per-capita recruitment and survival parameter, respectively. Average and min-max values represent the parameter set used from the 50 different datasets simulated for each population and scenario.

| Life History | Stochasticity | Median survival time | Asymptotic growth rate | SD growth rate | Average Super population size | Average $\rho$ [min-max] | Average $\psi$ [min-max] |
| --- | --- | --- | --- | --- | --- | --- | --- |
| Slow | Low | 4.9 | 1 | 0.07 | 80 | 0.15<br>[0.15-0.15] | 0.85<br>[0.85-0.85] |
|  | High | 4.9 | 1 | 0.10 | 80 | 0.15<br>[0.01-0.32] | 0.85<br>[0.68-1] |
| Intermediate | Low | 2.1 | 1 | 0.12 | 119 | 0.35<br>[0.35-0.35] | 0.65<br>[0.65-0.65] |
|  | High | 2.1 | 1 | 0.13 | 119 | 0.35<br>[0.23-0.49] | 0.65<br>[0.5-0.8] |

#### 2.2.2. Sampling interruption scenarios

We created 10 different sampling interruption scenarios (Figure 2) and a scenario without interruptions over five consecutive occasions (scenario 11111; Figure 2). When no sampling occurred during occasion *t*, we set *p*_*ijt*_ in the OPSCR model to 0 to specify that there was no possibility of detecting any individuals during that occasion.

**Figure 2.**
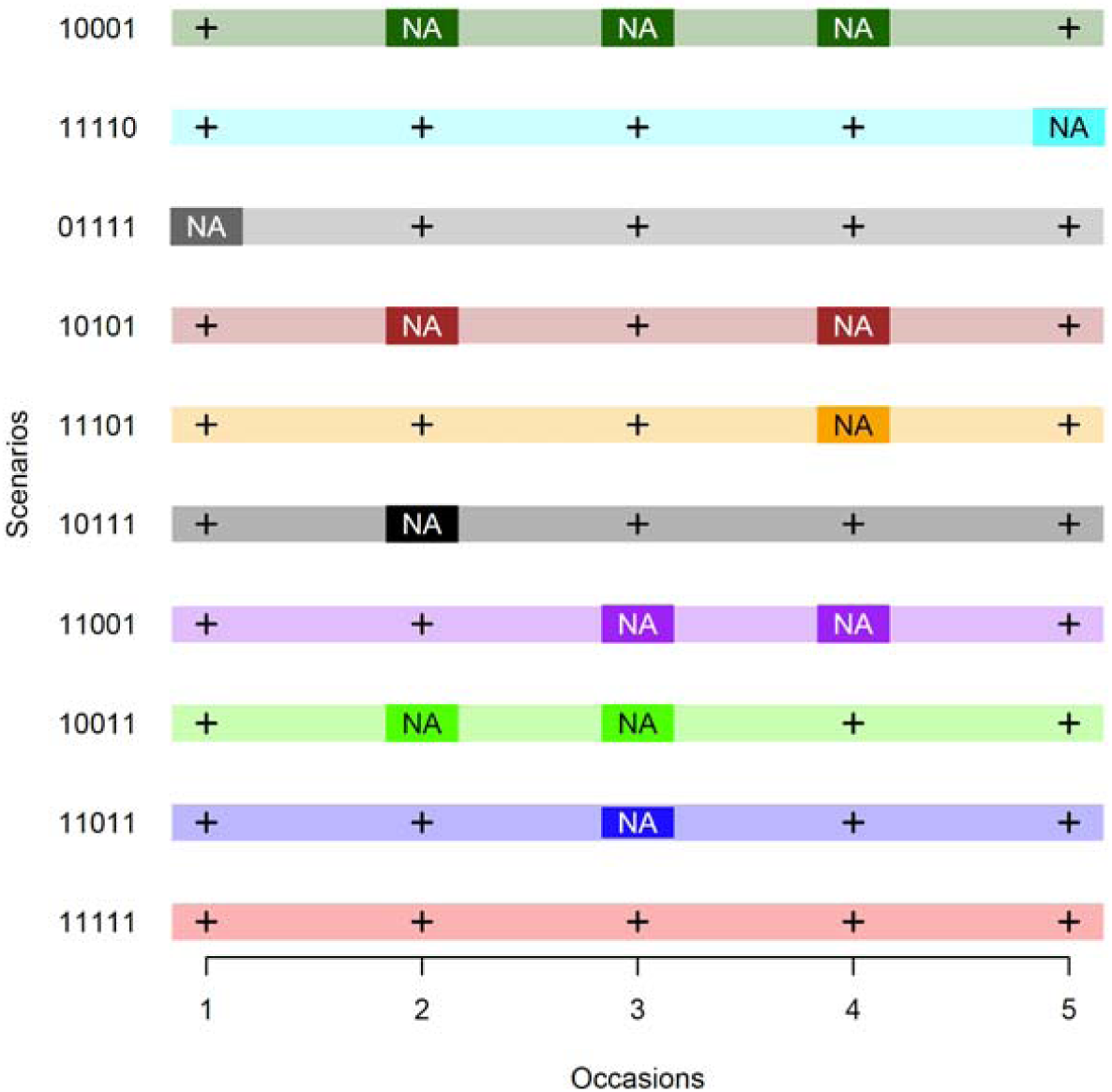
Visual representation of the 10 sampling interruption scenarios considered in the analysis. The x-axis denotes five consecutive sampling primary occasions. The 10 different scenarios are arranged along the y axis and are coded by binary values corresponding to whether sampling was performed (1) or not (0) during each occasion and is visually represented by “**+**” and “*NA*”, respectively.

### 2.3. Evaluation of model performance

We simulated 50 datasets for each of the 10 scenario and each of the four populations, resulting in 2000 simulated datasets. For each simulated data set, we calculated the relative bias 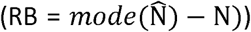 and the coefficient of variation 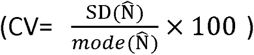, where SD is the standard deviation, 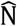 are the MCMC posterior samples of population size, and N is the true value of population size (Walther and Moore 2005). In addition, we calculated the 95% credible interval coverage as the percentage of simulations for which the credible interval contained the true value.

### 2.4. The wolverine data

We fit the OPSCR model to NGS data from the national monitoring program of wolverines in Norway (see description in (Flagstad et al. 2004, Brøseth et al. 2010, Bischof et al. 2016b, Gervasi et al. 2019)). We used data collected during five consecutive winters (January-May) between 2013 and 2017 in central Norway (Supporting information 1.2, Figure S1.2.1). The data consisted of 632 detections from 126 individually identified female wolverines. Samples were collected by field personnel from the management authorities (Norwegian Nature Inspectorate) using a search-encounter method on snow. During searches, the GPS coordinates of search-tracks were recorded. We used the partially aggregated binomial observation model (Milleret et al. 2018), which divides detectors into K subdetectors and models the frequency of subdetectors with more than one detection as a binomial response with a sample size of K. We located primary detectors in the center of grid cells (4km resolution) and subdetectors in the center of subdetector grid cells (800m resolution). We only placed subdetectors when search tracks overlapped with the subdetector grids. The configuration of active grid cells changed every year to account for spatial-temporal variation in searches. We also estimated year-specific *p0* to account for annual variation in sampling intensity. To increase computing efficiency, we used a local evaluation of the state-space to reduce the number of detectors considered for each individual during the model fit (Milleret et al. 2019). Searches were conducted continuously from 2013 to 2017, which allowed us to introduce different artificial gaps in the data time series, while having a reference point (scenario without gaps: 11111). We simulated sampling interruption by removing all detections from all individuals at the occasion(s) designated as interruption. We implemented the same 10 interruption scenarios as used in the simulations (Figure 2). We compared 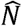 and its CV (i.e. obtained when excluding the buffer area, 63584km^2^) between the different scenarios.

### 2.5. Model fitting

We fitted the Bayesian OPSCR models using Markov chain Monte Carlo (MCMC) with nimble (Turek et al. 2016, de Valpine et al. 2017, NIMBLE Development Team 2019) in R version 3.3.3 (R core team 2017). NIMBLE provides a new implementation of the BUGS model language coupled with the capability to add new functions, distributions, and MCMC samplers to improve computing performance. We ran four chains with 40000 iterations each following a 2000-iteration burn-in. We considered models as converged when Rhat was ≤1.1 (Gelman and Rubin 1992) for all main parameters and by visually inspecting a sample of all repetitions of all scenarios. We re-ran models that did not reach convergence for 60000 iteration per chain following a 20000-iteration burn-in, and excluded them from the results if they still did not reach convergence. R and nimble codes for the OPSCR model, related custom functions, and simulations used are provided in Supporting Information S2, and list of priors used in Supporting Information 1.3 Table S1.3.1).

## 3. Results

### 3.1. Simulations

All models reached convergence, with the exception of scenario 10001 for species having an intermediate life history strategy (25% non-converged Supporting information 1.4, Table S1.4.1). We detected no systematic bias in 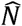, regardless of whether sampling interruption occurred or not (Supporting information 1.5, Table S1.5.1). However, the precision in 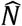 generally decreased towards the first and last occasions (e.g. Figure 1, scenario 11111). Regardless of when the interruption(s) occurred, the precision in 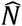 decreased for the affected occasion(s). For example, for the scenario 11011, CV of 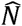 was on average 1.3 times higher during the third occasion (i.e. interruption) compared to the scenario without interruption in sampling (Figure3, Supporting information 1.5, Table S1.5.1). The increased uncertainty caused by interruptions also propagated to estimates for sampled occasions, especially for those adjacent to interruption(s). Precision of 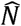 decreased as the number of interruptions increased. CV was on average 1.8 times higher for interruptions at the beginning or at the end of the study period, than for an interruption at the third occasion. Regardless of the interruption scenario, uncertainty in 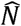 was larger for the intermediate life history scenario, but the presence of stochasticity in vital rates did not seem to amplify the depressing effect of interruptions on the precision of 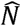.

**Figure 3.**
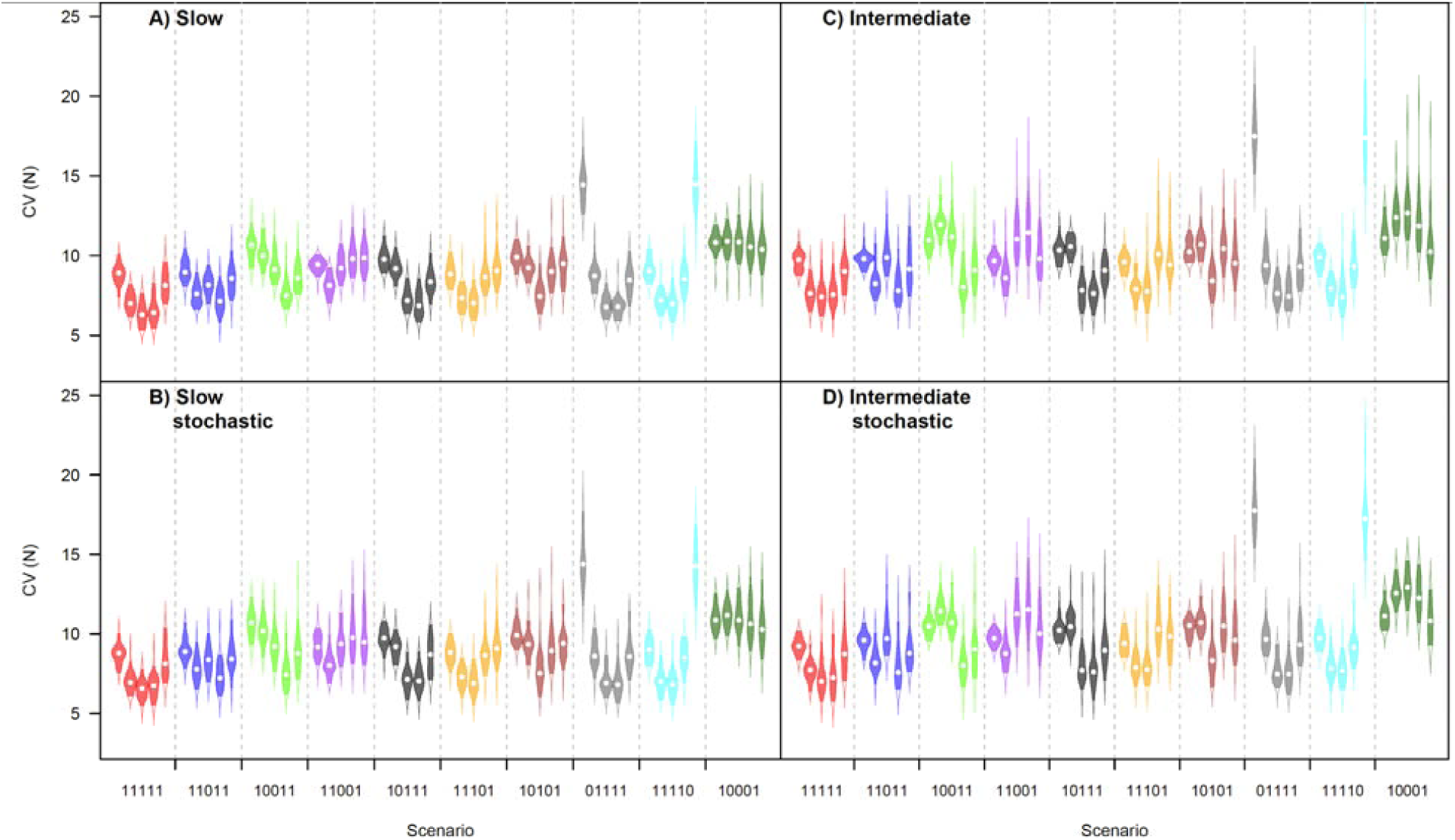
Violins plots (points: medians; solid colors: 95% credible interval) for the coefficient of variation (CV) of abundance estimates (N) obtained using an open population spatial capture recapture model fit to simulated datasets (50 repetitions for each scenario). Shown are results for simulations representing combinations of life history strategies (slow and intermediate), and with and without temporal stochasticity in vital rates. The five consecutive 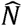 estimates (i.e corresponding to the five sampling occasions) are colored and grouped according to the sampling interruption scenario (x-axis). Sampling scenarios are presented by a series of 1s and 0s indicating whether sampling was considered to have occurred or not, respectively.

### 3.2. Wolverines

All models fit to the empirical wolverine data converged, except scenario 10001 for which the standard deviation of the Gaussian dispersal kernels (*τ*) did not reach the convergence criterion. Wolverine population size 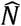 in the absence of sampling interruptions was relatively stable over the five consecutive years (>60 individuals, figure 4; 11111). We did not detect marked changes in 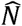 estimates when the data set was subjected to sampling interruptions (CI of all 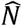 overlapped with each other regardless of the scenario, figure 4). However, patterns in CV of 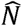 in response to sampling interruptions were similar to those observed for simulated data sets, with a higher uncertainty towards the first and last occasions and with a sampling interruption (Figure 4).

**Figure 4.**
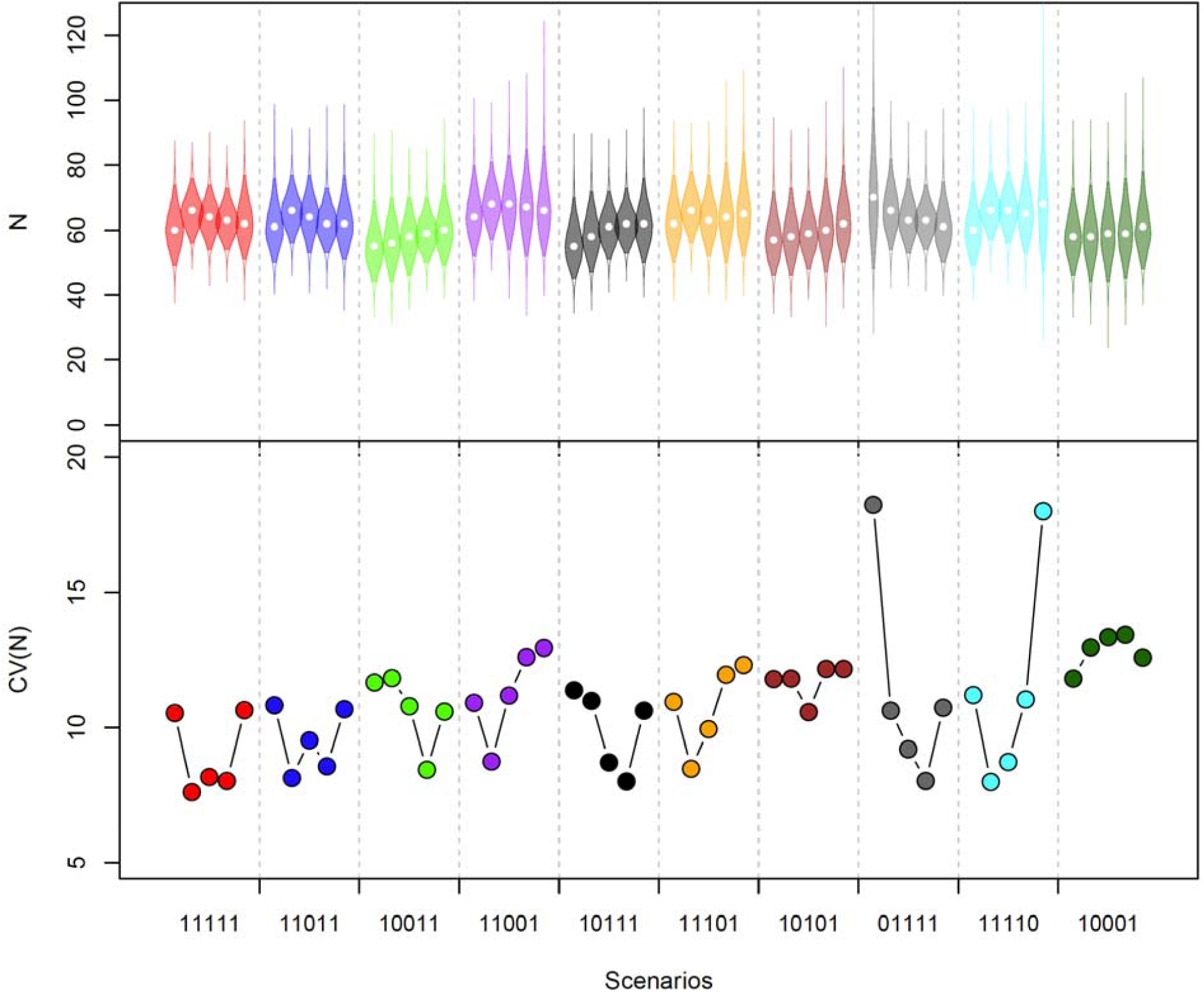
Violin plots (points: medians; solid colors: 95% credible interval) of the posterior distribution of abundance (N) (top panel) and its coefficient of variation (CV; bottom panel) obtained using an open population spatial capture recapture model on non-invasive genetic sampling data of wolverines collected in south-central Norway. The five consecutive annual 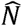 estimates and CV (2013-2017) are colored and grouped according to the sampling interruption scenario (x-axis). Sampling scenarios are presented by a series of 1s and 0s indicating whether sampling was considered to have occurred or not, respectively.

## 4. Discussion

Simulations and a case study on wolverines revealed that OPSCR models can be a valuable tool for abundance inferences when there are gaps in data time series. Although uncertainty in abundance estimates increased during occasions with a sampling interruption, the interruption did not seem to cause any systematic bias. Uncertainty in abundance estimates increased with the number of interruptions and the speed of the study species’ life history. Similarly, the simulated sampling interruptions in the wolverine example (a species with an intermediate life history strategy; *ϕ* = 0.7(95%*CI*: 0.62 − 0.78); *ρ* = 0.3 (95%*CI*: 0.21 − 0.39)) showed that interruptions caused higher uncertainty around abundance estimates, but that abundance estimates were relatively similar to those in the absence of interruptions (Figure 4). The effect of interruptions on precision was generally less pronounced when the gap in the time series was framed by several consecutive sampled occasions (11011). Although OPSCR models have already been used to infer abundance in the presence of interruptions (Chandler and Clark 2014, Augustine et al. 2019), our study is the first to explore the conditions under which reliable abundance inferences can be obtained when SCR data time series include temporal gaps in sampling.

Compared to a series of independent SCR models, OPSCR models use detections and model population dynamics and individual movement between several consecutive sampling occasions. As a result, individual detections in previous and/or subsequent occasions inform the Markovian model about the spatial location and demographic status of each individual and help determine its fate (Molinari et al. (2018); Figure 1). This explains the increase precision of the estimates for gaps framed by multiple occasions with data (Figure 3, scenario 11011). Despite a loss in precision of abundance estimates, the OPSCR model, and its Markovian structure, allows the reliable estimation of abundance in the presence of interruptions. However, the presence of sampling interruption pose a greater challenge to estimation when the lifetime of the species is short compared to the time interval between consecutive surveys. Indeed, we found that for species with intermediate life histories precision of abundance estimates was lower and models took longer to converge than for species with slow life histories.(Supporting information 1.4, Table S1.4.1.)

Movement of ACs between occasions is an important feature of OPSCR models and a miss-specified movement process can have important consequences for inferences (Ergon and Gardner 2014, Gardner et al. 2018). For the purpose of this study, we developed a Markovian movement model assuming distance between consecutive individual ACs being normally distributed. The movement model is essential to distinguish between mortality and emigration (Ergon and Gardner 2014) and assists the OPSCR in predicting the fate of individuals that are not detected during interruptions in sampling (Figure 1). Based on the locations of the AC at occasions prior to and following interruption(s), together with population level information about AC movement, the model makes prediction about the location of individuals ACs during occasions with missing data (e.g. prediction of the movement of individuals in and out of the study area). This is particularly useful as the OPSCR model not only yields population size estimates that bridge interruptions in sampling, but can also estimate density across the study area during years without sampling.

In this analysis, we considered that interruptions occurred at random and not because of a specific event (e.g. unfavorable climatic conditions) that could have, not only prevented the occurrence of sampling, but also affected the population. Independence of the probability of interruptions from biological processes affecting parameters of interest (Nakagawa and Freckleton 2008). When it is met, key parameters (e.g. σ, *ϕ, ρ*) are transferable between years and the model should return unbiased abundance estimates for gap years. Otherwise, investigators should use caution when drawing inferences for gap years, as the occasions with and without observations may be confounded with differences in biological processes.

The main goal of many wildlife monitoring programs is to obtain reliable estimates of population size and trends therein, but also to understand the mechanisms (e.g. recruitment, survival) involved in population size fluctuations when planning conservation and management actions. Although individual survival between occasions is informed through the reconstruction of individual states during interruptions, under some circumstances, parameter identifiability can be weak when parameters are allowed to vary over time (see Supporting information 1.6). In order to estimate survival and recruitment in the presence of sampling interruptions, it may be necessary to assume that these vital rates are constant over time, as we did in our example. However, estimation of time dependent vital rates, despite gaps in the data time series, may be facilitated through the use of random effects (e.g. year on survival or recruitment) or time-dependent covariates explaining temporal variation in vital rates (e.g. changes in environmental conditions, hunting intensity). In the simulations, we added un-modelled temporal stochasticity in vital rates, which did not have a marked impact on inferences. This suggests that OPSCR models are relatively robust to temporal stochasticity in vital rates, as long as its magnitude remain relatively low. Additionally, the integration of other types of data (e.g. unmarked individuals (Sollmann et al. 2013, Chandler and Clark 2014), and dead recoveries (Proffitt et al. 2015)), could be used to mitigate the loss of information due to sampling interruption.

## Conclusion

The framework described here allows ecologists to assess the impact of sampling interruptions – whether intentional or unintentional – on parameter estimates from OPSCR models. Based on our findings, we recommend that intentional interruption be restricted to species with life histories that are slow (relative to the monitoring interval) and to avoid multiple consecutive interruptions. Methods allowing the integration of different types of data (e.g. unmarked individuals, dead recoveries) into OPSCR models could help further mitigate the negative impact of interruptions on the precision of parameter estimates (see Chandler and Clark (2014) for an example). Previous studies testing the cost-efficiency of non-spatial CR surveys have focused on the importance of study duration, proportion of different individuals sampled, and detection probability (Lieury et al. 2017). Unless the study species requires close monitoring due to short response times for management interventions (e.g. endangered species), the use of OPSCR model for cases with periodic interruptions in sampling could be considered as an option to distribute sampling efforts over time and make long-term population-level monitoring cost-effective (Chandler and Clark 2014).

## Supporting information

Supporting informations

## ACKNOWLEDGMENT

This work was funded by the Norwegian Environment Agency (Miljødirektoratet), the Swedish Environmental Protection Agency (Naturvårdsverket) and the Research Council of Norway (NFR 286886). We thank the field staff from the State Nature Inspectorate and members of the public that collected the monitoring data for the large carnivore database Rovbase3.0 (rovbase.no).

## AUTHOR’S CONTRIBUTION

C.M, P.D, J.C. and R.B developed the concept and methodology with input from D.T, O.G, P.d.V. Wolverine data extraction and preparation were coordinated by H.B. C.M led the analysis with help from P.D, J.C, R.B, O.G, D.T and P.d.V. Point process model was developed by J.C. C.M led the writing with contributions from P.D, J.C, O.G, and R.B. All authors contributed critically to drafts of the manuscript and gave final approval for publication.

## DATA ACCESSIBILITY

R code to reproduce simulations is available in the supporting information and wolverine data will be uploaded on dryad repository upon acceptance.

## References

Augustine, B. C., M. Kéry, J. O. Marin, P. Mollet, G. Pasinelli, and C. Sutherland. 2019. Sex-specific population dynamics and demography of capercaillie (Tetrao urogallus L.) in a patchy environment. bioRxiv.

Bears, H., K. Martin, and G. C. White. 2009. Breeding in high-elevation habitat results in shift to slower life-history strategy within a single species. Journal of Animal Ecology 78:365–375.

Bischof, R., H. Brøseth, and O. Gimenez. 2016a. Wildlife in a Politically Divided World: Insularism Inflates Estimates of Brown Bear Abundance. Conservation Letters 9:122–130.

Bischof, R., E. R. Gregersen, H. Brøseth, H. Ellegren, and Ø. Flagstad. 2016b. Noninvasive genetic sampling reveals intrasex territoriality in wolverines. Ecology and Evolution 6:1527–1536.

Borchers, D. L., and M. G. Efford. 2008. Spatially Explicit Maximum Likelihood Methods for Capture-Recapture Studies. Biometrics 64:377–385.

Brøseth, H., Ø. Flagstad, C. Wärdig, M. Johansson, and H. Ellegren. 2010. Large-scale noninvasive genetic monitoring of wolverines using scats reveals density dependent adult survival. Biological Conservation 143:113–120.

Chandler, R. B., and J. D. Clark. 2014. Spatially explicit integrated population models. Methods in Ecology and Evolution 5:1351–1360.

Dawson, D. K., and M. G. Efford. 2009. Bird population density estimated from acoustic signals. Journal of Applied Ecology 46:1201–1209.

Efford, M. 2004. Density estimation in live-trapping studies. Oikos 106:598–610.

Efford, M. G., D. K. Dawson, and D. L. Borchers. 2009. Population density estimated from locations of individuals on a passive detector array. Ecology 90:2676–2682.

Ergon, T., and B. Gardner. 2014. Separating mortality and emigration: modelling space use, dispersal and survival with robust-design spatial capture-recapture data. Methods in Ecology and Evolution 5:1327–1336.

Flagstad, Ø., E. Hedmark, A. Landa, H. Brøseth, J. Persson, R. Andersen, P. Segerström, and H. Ellegren. 2004. Colonization History and Noninvasive Monitoring of a Reestablished Wolverine Population. Conservation Biology 18:676–688.

Gardner, B., R. Sollmann, N. S. Kumar, D. Jathanna, and K. U. Karanth. 2018. State space and movement specification in open population spatial capture-recapture models. Ecology and Evolution 0.

Gelman, A., and D. B. Rubin. 1992. Inference from iterative simulation using multiple sequences. Statistical Science 7:457–511.

Gervasi, V., J. D. C. Linnell, H. Brøseth, and O. Gimenez. 2019. Failure to coordinate management in transboundary populations hinders the achievement of national management goals: The case of wolverines in Scandinavia. Journal of Applied Ecology 0.

Gimenez, O., V. Rossi, R. Choquet, C. Dehais, B. Doris, H. Varella, J.-P. Vila, and R. Pradel. 2007. State-space modelling of data on marked individuals. Ecological Modelling 206:431–438.

Illian, J., A. Penttinen, H. Stoyan, and D. Stoyan. 2008. Statistical analysis and modelling of spatial point patterns. John Wiley & Sons.

Kery, M., and M. Schaub. 2011. Bayesian Population Analysis using WinBUGS: A Hierarchical Perspective. Elsevier Science.

Kindberg, J., J. E. Swenson, G. Ericsson, E. Bellemain, C. Miquel, and P. Taberlet. 2011. Estimating population size and trends of the Swedish brown bear Ursus arctos population. Wildlife Biology 17:114–123.

Lieury, N., S. Devillard, A. Besnard, O. Gimenez, O. Hameau, C. Ponchon, and A. Millon. 2017. Designing cost-effective capture-recapture surveys for improving the monitoring of survival in bird populations. Biological Conservation 214:233–241.

Lindenmayer, D. B., and G. E. Likens. 2009. Adaptive monitoring: a new paradigm for long-term research and monitoring. Trends in Ecology & Evolution 24:482–486.

López-Bao, J. V., R. Godinho, C. Pacheco, F. J. Lema, E. Garcia, L. Llaneza, V. Palacios, and J. Jiménez. 2018. Toward reliable population estimates of wolves by combining spatial capture-recapture models and non-invasive DNA monitoring. Scientific reports 8:2177.

Milleret, C., P. Dupont, C. Bonenfant, H. Brøseth, Ø. Flagstad, C. Sutherland, and R. Bischof. 2019. A local evaluation of the individual state-space to scale up Bayesian spatial capture–recapture. Ecology and Evolution 9:352–363.

Milleret, C., P. Dupont, H. Brøseth, J. Kindberg, A. Royle J., and R. Bischof. 2018. Using partial aggregation in spatial capture recapture. Methods in Ecology and Evolution 0.

Molinari-Jobin, A., M. Kéry, E. Marboutin, F. Marucco, F. Zimmermann, P. Molinari, H. Frick, C. Fuxjäger, S. Wölfl, F. Bled, C. Breitenmoser-Würsten, I. Kos, M. Wölfl, R. Cerne, O. Müller, and U. Breitenmoser. 2018. Mapping range dynamics from opportunistic data: spatiotemporal modelling of the lynx distribution in the Alps over 21 years. Animal Conservation 21:168–180.

Nakagawa, S., and R. P. Freckleton. 2008. Missing inaction: the dangers of ignoring missing data. Trends in Ecology & Evolution 23:592–596.

NIMBLE Development Team. 2019. NIMBLE: MCMC, Particle Filtering, and Programmable Hierarchical Modeling. https://cran.r-project.org/package=nimble.

Plummer, M. 2003. JAGS: A program for analysis of Bayesian graphical models using Gibbs sampling. Page 125Proceedings of the 3rd international workshop on distributed statistical computing.

Proffitt, K. M., J. F. Goldberg, M. Hebblewhite, R. Russell, B. S. Jimenez, H. S. Robinson, K. Pilgrim, and M. K. Schwartz. 2015. Integrating resource selection into spatial capture-recapture models for large carnivores. Ecosphere 6:art239.

Royle, J. A., R. B. Chandler, R. Sollmann, and B. Gardner. 2014. Spatial Capture-Recapture. Academic Press.

Royle, J. A., R. M. Dorazio, and Link W.A. 2007. Analysis of multinomial models with unknown index using data augmentation. Journal of Computational and Graphical Statistics 16(1):67–85.

Royle, J. A., K. U. Karanth, A. M. Gopalaswamy, and N. S. Kumar. 2009. Bayesian inference in camera trapping studies for a class of spatial capture–recapture models. Ecology 90:3233–3244.

Royle, J. A., A. J. Magoun, B. Gardner, P. Valkenburg, and R. E. Lowell. 2011. Density estimation in a wolverine population using spatial capture–recapture models. The Journal of Wildlife Management 75:604–611.

Royle, J. A., and K. V Young. 2008. A hierarchical model for spatial capture–recapture data. Ecology 89:2281–2289.

Sanz–Aguilar, A., R. Pradel, and G. Tavecchia. 2019. Age–dependent capture–recapture models and unequal time intervals. Animal Biodiversity and Conservation 42.1:91–98.

Schmidt, B. R., M. Schaub, and S. Steinfartz. 2007. Apparent survival of the salamander Salamandra salamandra is low because of high migratory activity. Frontiers in Zoology 4:19.

Schwarz, C. J., and A. N. Arnason. 1996. A General Methodology for the Analysis of Capture-Recapture Experiments in Open Populations. Biometrics 52:860–873.

Seber, G. A. F. 1965. A Note on the Multiple-Recapture Census. Biometrika 52:249–259.

Seber, G. A. F. 1982. The Estimation of Animal Abundance and Related Parameters (2nd ed.). London: Edward Arnold.

Sollmann, R., B. Gardner, A. W. Parsons, J. J. Stocking, B. T. McClintock, T. R. Simons, K. H. Pollock, and A. F. O’Connell. 2013. A spatial mark–resight model augmented with telemetry data. Ecology 94:553–559.

Stearns, S. C. 1992. The evolution of life histories.

Swenson, J. E., M. Schneider, A. Zedrosser, A. Söderberg, R. Franzén, and J. Kindberg. 2017. Challenges of managing a European brown bear population; lessons from Sweden, 1943--2013. Wildlife Biology 2017:wlb--00251.

Turek, D., P. de Valpine, and C. J. Paciorek. 2016. Efficient Markov chain Monte Carlo sampling for hierarchical hidden Markov models. Environmental and Ecological Statistics 23:549–564.

de Valpine, P., D. Turek, C. J. Paciorek, C. Anderson-Bergman, D. T. Lang, and R. Bodik. 2017. Programming with models: writing statistical algorithms for general model structures with NIMBLE. Journal of Computational and Graphical Statistics 26:403–413.

Walther, B. A., and J. L. Moore. 2005. The concepts of bias, precision and accuracy, and their use in testing the performance of species richness estimators, with a literature review of estimator performance. Ecography 28:815–829.

Zabala, J., I. Zuberogoitia, J. A. Martínez-Climent, and J. Etxezarreta. 2011. Do long lived seabirds reduce the negative effects of acute pollution on adult survival by skipping breeding? A study with European storm petrels (Hydrobates pelagicus) during the “Prestige” oil-spill. Marine Pollution Bulletin 62:109–115.

Zuberogoitia, I., J. Zabala, J. Etxezarreta, A. Crespo, G. Burgos, and J. Arizaga. 2016. Assessing the impact of extreme adverse weather on the biological traits of a European storm petrel colony. Population Ecology 58:303–313.

