## Supporting informations for "Estimating abundance with interruptions in data collection using open population spatial capture-recapture models": SourceNimbleObservationModel.R

```
##########################################
## OBSERVATION MODEL NIMBLE FUNCTIONS ####
##########################################
## ==== 1. calculateDistance ==== 
#' @title NIMBLE Function to calculate the square distance between an AC location and a vector of detectors locations.
#'
#' @description
#' \code{calculateDistance} is a NIMBLE Function to calculate the square distance between an AC location and several detectors.
#' 
#' @param sxy \code{Vector} of length 2 containing location X and Y of the activity center of one individual . 
#' @param detector.xy \code{Matrix} with nrow = n.detectors and ncol the X and Y coordinates.
#' @param indicator \code{numeric} denotes whether the individual is considered alive (1) or dead (0) and whether the  square distance should be calculated or not. .
#'
#' @examples
#' d2[i,1:n.detectors,t] <- calculateDistance(sxy[i,1:2,t], detector.xy[1:n.detectors,1:2], z[i,t]==2)

calculateDistance <- nimbleFunction(run = function( sxy = double(1)
                                                    , detector.xy = double(2)
                                                    , indicator = double(0, default = 1.0)){
  # Return type declaration
  returnType(double(1))
  
  # Check input dimensions
  n.detectors <- dim(detector.xy)[1]
  if(indicator == 0){
    return(rep(0,n.detectors))
  }
  # Calculate distance vector (Output)
  d2 <- pow(detector.xy[1:n.detectors,1] - sxy[1], 2) + pow(detector.xy[1:n.detectors,2] - sxy[2], 2)
  # Return square distance
  return(d2)
})


## ==== 2. dbin_LESS_vector ==== 
#' @title Function to create a NIMBLE custom distribution for faster SCR model runs.
#'
#' @description
#' \code{dbin_LESS} returns the likelihood of a given individual & spatial binary detection history y[i,1:n.detectors] 
#' 
#' @param x \code{Vector} of length n.detectors containing detection/non detections 
#' @param pZero \code{Numeric} variable denoting the baseline detection parameter of the half-normal detection function.
#' @param sigma \code{Numeric} variable denoting the scale parameter of the half-normal detection function.
#' @param sxy A \code{Vector} of length 2 with individual activity center x and y coordinates.
#' @param d2 A \code{Vector}  of dimensions n.detectors with the square distances from the activity centers to each detector obtained from the \code{calculateDistance}.
#' @param maxDist A \code{Numeric} with the maximum distance to the AC where detections are considered possible. This applies a local evaluation of the state space (Milleret et al. 2018 Ecology and Evolution)
#' @param indicator A \code{Logical}; if == 0 no detections possible; saves time. 
#' @param log A \code{integer} indicates whether the log-probability or the probabilty should be returned (default to normal)
#'
#' @examples
#' y[i,1:n.detectors,t] ~ dbern_LESS(p0, sigma, d2[i,1:n.detectors,t], maxDist,  z[i,t]==2)

## ==== 2.1.Density function ====
dbin_LESS <- nimbleFunction(run = function( x = double(1)
                                            , pZero = double(0)
                                            , sigma = double(0)
                                            , trials = double(1)
                                            , d2 = double(1)
                                            , maxDist = double(0, default = 0.0)
                                            , indicator = double(0, default = 1.0)
                                            , log = integer(0, default = 0)){
  # Return type declaration
  returnType(double(0))
  
  ## Check input dimensions
  n.detectors <- length(x)
  
  ## Shortcut if individual is not available for detection
  if(indicator == 0){
    if(sum(x[1:n.detectors]) == 0){
      if(log == 0) return(1.0)
      else return(0.0)
    }else{
      if(log == 0) return(0.0)
      else return(-Inf)
    }
  }
  
  
  ## Calculate the likelihood of the detection observations
  alpha <- -1.0 / (2.0 * sigma * sigma)
  detectLogLikeli <- numeric(n.detectors, init = FALSE)
  p <- numeric(n.detectors, init = FALSE)
  
  maxDist_squared <- maxDist*maxDist
  for(j in 1:n.detectors){
    # Calculate the detection probability (negative distances repesent zero detection probability)
    if(d2[j] <= (maxDist_squared)){p[j] <- pZero * exp(alpha * d2[j])} else {p[j] <- 0.0000000}
  }#j
  
  
  logProb <- sum(dbinom(x, prob = p, size = trials, log = TRUE))
  if(log)return(logProb)
  return(exp(logProb))
  
})

## ==== 2.2 Sampling function ====
rbin_LESS <- nimbleFunction(run = function( n = integer(0)
                                            , pZero = double(0)
                                            , sigma = double(0)
                                            , trials = double(1)
                                            , d2 = double(1)
                                            , maxDist = double(0, default = 0.0)
                                            , indicator = double(0, default = 1.0)){
  # Return type declaration
  returnType(double(1))
  
  # Check input dimensions
  n.detectors <- length(d2)
  
  ## Shortcut if individual is not available for detection
  if(indicator == 0){return(rep(0.0, n.detectors))}
  
  
  ## Simulate the detections using the calculated detection distances ----
  alpha <- -1.0 / (2.0 * sigma * sigma)
  # Initialise a detections output vector
  detectOut <- rep(0, n.detectors)
  for(j in 1:n.detectors){
    # Calculate the detection probability (negative distances repesent zero detection probability)
    if(d2[j] <= (maxDist*maxDist)){
      p <- pZero * exp(alpha * d2[j])
      # Draw from a Bernoulli distribution with the calculated probability
      detectOut[j] <- rbinom(1, trials[j], p)
    }#if
  }#j
  
  # Output
  return(detectOut)
})

## ==== 2.3. Registration ====
registerDistributions(list(
  dbin_LESS = list(
    BUGSdist = "dbin_LESS(pZero, sigma, trials, d2, maxDist, indicator)",
    types = c( "value = double(1)", "pZero = double(0)", "sigma = double(0)", "trials = double(1)", "d2 = double(1)", 
               "maxDist = double(0)" ,"indicator = double(0)"),
    pqAvail = FALSE)))
```
