## Supporting informations for "Estimating abundance with interruptions in data collection using open population spatial capture-recapture models": SourceNimblePointProcess.R

```
## 0. ------ DEREGISTER ANY PREVIOUSLY REGISTERED DISTRIBUTIONS ------
# In some versions of NIMBLE the distribution registeration proceedures can return an error if the distributions
# have already been previously registered.  This little section of code deregisters the functions to ensure no
# errors are thrown if this file is sourced more than once.
if(exists("distributions", nimbleUserNamespace)) {
   # List of distributions defined in this source file
   distributionNames <- c(
      "dbinomPP",
      "dbinomPPSingle",
      "dbinomMNormSourcePP",
      "dbinomMNormSourcePPSingle")
   # Find those distributions that are defined in this source fiel and see if they are already registered
   isDefinedDist <- distributionNames %in% nimbleUserNamespace$distributions$namesVector
   if(any(isDefinedDist)) {
      # If the distributions are registered then deregister them
      deregisterDistributions(distributionNames[isDefinedDist])
   }
}

## 1. ------ DEFINE NIMBLE CUSTOM UTILITY FUNCTIONS USED IN THE POINT PROCESS DISTRIBUTIONS ------

### 1.1. ==== Calcualte the size of the observation window ====
calcObsWindowSize <- nimbleFunction(
   run = function(
      lowerCoords = double(2),
      upperCoords = double(2),
      areAreas = double(0, default = 1)
   ) {
      ## 1.1.1. Specify the return type dimensionality ----
      # Return type declaration
      returnType(double(1))
      ## 1.1.2. Sanity test the inputs ----
      # Assess the dimensionality of the input coordinates
      dimCoords <- dim(lowerCoords)[2]
      # Ensure that the dimensionality is valid
      if(dimCoords <= 0) {
         stop("invalid dimension structure for the input coordinates")
      }
      # Assess the number of observation windows
      numObsWindows <- dim(lowerCoords)[1]
      if(numObsWindows <= 0) {
         stop("invalid number of observation windows")
      } else if(numObsWindows != dim(upperCoords)[1]) {
         stop("number of observation windows not consistent between lower and upper coordinates")
      }
      if(dimCoords != dim(upperCoords)[2]) {
         stop("lower and/or upper coordinates have an incorrect dimension structure")
      }
      ## 1.1.3. Calculate the area/length of the observation window ----
      obsWindowSize <- rep(0, numObsWindows)
      if(areAreas) {
         # The observation windows are areas/volumes and therefore calculate their area/volume
         # accordingly
         obsWindowSize <- obsWindowSize + 1.0
         for(dimIter in 1:dimCoords) {
            obsWindowSize <- obsWindowSize * (upperCoords[1:numObsWindows, dimIter] - lowerCoords[1:numObsWindows, dimIter])
            if(sum(obsWindowSize <= 0.0) > 0) {
               # Ensure that upper and lower coordinates are valid
               stop("window area is zero for at least one observation window")
            }
         }
      } else {
         # The observation windows are transects and therefore calculate their length accordingly
         for(dimIter in 1:dimCoords) {
            coordDist <- upperCoords[1:numObsWindows, dimIter] - lowerCoords[1:numObsWindows, dimIter]
            obsWindowSize <- obsWindowSize + coordDist * coordDist
         }
         if(sum(obsWindowSize <= 0.0) > 0) {
            # Ensure that upper and lower coordinates are valid
            stop("window length is zero for at least one observation window")
         }
         obsWindowSize <- sqrt(obsWindowSize)
      }
      return(obsWindowSize)
   }
)

### 1.2. ==== Calculate the normalisation integral of the multivariate normal source-point model ====
calcMNormSourceInt <- nimbleFunction(
   run = function(
      lowerCoords = double(2),
      upperCoords = double(2),
      sourceCoords = double(1),                    # The coordinates of the source location (the origin of the decay kernel)
      normSD = double(0),                          # The standard deviation of the isotropic multivariate normal distribution decay kernel
      areAreas = double(0, default = 1)
   ) {
      ## 1.2.1. Specify the return type dimensionality ----
      # Return type declaration
      returnType(double(1))
      ## 1.2.2. Sanity test the inputs ----
      # Assess the dimensionality of the input coordinates
      dimCoords <- dim(lowerCoords)[2]
      # Ensure that the dimensionality is valid
      if(dimCoords <= 0) {
         stop("invalid dimension structure for the input coordinates")
      }
      # Assess the number of observation windows
      numObsWindows <- dim(lowerCoords)[1]
      if(numObsWindows <= 0) {
         stop("invalid number of observation windows")
      } else if(numObsWindows != dim(upperCoords)[1]) {
         stop("number of observation windows not consistent between lower and upper coordinates")
      }
      if(dimCoords != dim(upperCoords)[2]) {
         stop("lower and/or upper coordinates have an incorrect dimension structure")
      }
      # Ensure that the source coordinates have the correct dimensionality
      if(length(sourceCoords) != dimCoords) {
         stop("source coordinates do match the coordinates of the observation windows")
      }
      # Ensure that the standard deviation of the isotropic multivariate normal distribution
      # has a valid value
      if(normSD <= 0.0) {
         stop("invalid value given for the standard deviation of the isotropic normal distribution")
      }
      ## 1.2.3. Calculate the normalisation integral ----
      # Initialise a vector to store the output
      normIntegral <- numeric(length = numObsWindows, value = 0.0, recycle = TRUE)
      if(areAreas) {
         # The observation windows are areas/volumes
         normIntegral <- normIntegral + pow(2.0 * pi * normSD * normSD, dimCoords / 2.0)
         # Iterate over the number of dimensions
         for(dimIter in 1:dimCoords) {
            if(sum(upperCoords[1:numObsWindows, dimIter] <= lowerCoords[1:numObsWindows, dimIter]) > 0) {
               # Ensure that upper and lower coordinates are valid
               stop("window area is zero for at least one observation window")
            }
            # Calculate the multivariate-normal normalisation integral for the current dimension
            normIntegral <- normIntegral * (pnorm((upperCoords[1:numObsWindows, dimIter] - sourceCoords[dimIter]) / normSD, 0.0, 1.0) - pnorm((lowerCoords[1:numObsWindows, dimIter] - sourceCoords[dimIter]) / normSD, 0.0, 1.0))
         }
      } else {
         # The observation windows are transects
         # Currently not supported
         stop("currently transect observation windows are not supported for multivariate normal source point models")
      }
      # Return the normalisation interal
      return(normIntegral)
   }
)

### 1.3. ==== Calculate whether a point within a set of observation windows ====
isWithinWindow <- nimbleFunction(
   run = function(
      curCoords = double(1),
      lowerCoords = double(2),
      upperCoords = double(2),
      tolDist = double(0),
      areAreas = double(0, default = 1)
   ) {
      ## 1.3.1. Specify the return type dimensionality ----
      # Return type declaration
      returnType(double(1))
      ## 1.3.2. Sanity test the inputs ----
      # Assess the dimensionality of the input coordinates
      dimCoords <- dim(lowerCoords)[2]
      # Ensure that the dimensionality is valid
      if(dimCoords <= 0) {
         stop("invalid dimension structure for the input coordinates")
      }
      # Assess the number of observation windows
      numObsWindows <- dim(lowerCoords)[1]
      if(numObsWindows <= 0) {
         stop("invalid number of observation windows")
      } else if(numObsWindows != dim(upperCoords)[1]) {
         stop("number of observation windows not consistent between lower and upper coordinates")
      }
      if(dimCoords != dim(upperCoords)[2]) {
         stop("lower and/or upper coordinates have an incorrect dimension structure")
      }
      # Ensure that the source coordinates have the correct dimensionality
      if(length(curCoords) != dimCoords) {
         stop("point coordinates do match the coordinates of the observation windows")
      }
      # Ensure that the tolerance distance is correctly set
      if(tolDist < 0.0) {
         stop("tolerance distance cannot be negative")
      }
      ## 1.3.3. Assess intersection between the points and the observation windows ----
      isInObsWindow <- numeric(length = numObsWindows, value = 1.0, recycle = TRUE)
      if(areAreas) {
         # The observation window is an area/volume so assess that it falls within
         # that volume
         for(dimIter in 1:dimCoords) {
            if(sum(lowerCoords[1:numObsWindows, dimIter] >= upperCoords[1:numObsWindows, dimIter]) > 0) {
               stop("upper coordinates must be greater than lower coordinates in each dimension")
            }
            isInObsWindow <- isInObsWindow * 
               numeric(value = rep(curCoords[dimIter], numObsWindows) >= lowerCoords[1:numObsWindows, dimIter] &
                   rep(curCoords[dimIter], numObsWindows) < upperCoords[1:numObsWindows, dimIter], length = numObsWindows)
         }
      } else {
         # The observation window is a line so test to see if the point falls on that
         # line
         isInObsWindow <- numeric(value = rep(curCoords[1], numObsWindows) >= lowerCoords[1:numObsWindows, 1] &
            rep(curCoords[1], numObsWindows) < upperCoords[1:numObsWindows, 1], length = numObsWindows)
         if(dimCoords > 1) {
            for(dimIter in 2:dimCoords) {
               isInObsWindow <- isInObsWindow * numeric(value = abs(rep(curCoords[dimIter], numObsWindows) - (
                  lowerCoords[1:numObsWindows, dimIter] + (upperCoords[1:numObsWindows, dimIter] - lowerCoords[1:numObsWindows, dimIter]) * (curCoords[1] - lowerCoords[1:numObsWindows, 1]) / (upperCoords[1:numObsWindows, 1] - lowerCoords[1:numObsWindows, 1])
               )) <= tolDist, length = numObsWindows)
            }
         }
      }
      return(isInObsWindow)
   }
)

### 1.4. ==== Find the nearest point in each observation window to a source point ====
nearestObsWindowPoint <- nimbleFunction(
   run = function(
      lowerCoords = double(2),
      upperCoords = double(2),
      sourceCoords = double(1),                    # The coordinates of the source location (the origin of the decay kernel)
      areAreas = double(0, default = 1)
   ) {
      ## 1.4.1. Specify the return type dimensionality ----
      # Return type declaration
      returnType(double(2))
      ## 1.4.2. Sanity test the inputs ----
      # Assess the dimensionality of the input coordinates
      dimCoords <- dim(lowerCoords)[2]
      # Ensure that the dimensionality is valid
      if(dimCoords <= 0) {
         stop("invalid dimension structure for the input coordinates")
      }
      # Assess the number of observation windows
      numObsWindows <- dim(lowerCoords)[1]
      if(numObsWindows <= 0) {
         stop("invalid number of observation windows")
      } else if(numObsWindows != dim(upperCoords)[1]) {
         stop("number of observation windows not consistent between lower and upper coordinates")
      }
      if(dimCoords != dim(upperCoords)[2]) {
         stop("lower and/or upper coordinates have an incorrect dimension structure")
      }
      # Ensure that the source coordinates have the correct dimensionality
      if(length(sourceCoords) != dimCoords) {
         stop("source coordinates do match the coordinates of the observation windows")
      }
      ## 1.4.3. Calculate the set of nearest point to the source coordinates for each observation window ----
      nearCoords <- matrix(value = 0.0, nrow = numObsWindows, ncol = dimCoords, recycle = TRUE)
      if(areAreas) {
         # Observation windows are areas/volumes so calculate the nearest point appropriately
         for(dimIter in 1:dimCoords) {
            nearCoords[1:numObsWindows, dimIter] <- pmin(
               pmax(rep(sourceCoords[dimIter], numObsWindows), lowerCoords[1:numObsWindows, dimIter]),
               upperCoords[1:numObsWindows, dimIter])
         }
      } else {
         # Observation windows are transects so calculate the nearest point appropriately
         # Currently not supported
         stop("currently transect observation windows are not supported for multivariate normal source point models")
      }
      return(nearCoords)
   }
)

### 1.5. ==== Calculate the integral of the normal distribution function ====
# Utility function to calculate the integral of the normal distribution function
normalDistIntegral <- nimbleFunction(
   run = function(
      curVals = double(1)
   ) {
      ## 1.5.1. Specify the return type dimensionality ----
      returnType(double(1))
      ## 1.5.2. Calculate the integrated density ----
      outVals <- curVals * pnorm(curVals, 0.0, 1.0) + dnorm(curVals, 0.0, 1.0)
      return(outVals)
   }
)

### 1.7. ==== Lookup the index of an observation window that contains a set of source coordinates ====
# Function to input a matrix of source coordinates and return a vcetor of the indeces of the observation
# windows that the observation window falls within.
obsWindowIndex <- nimbleFunction(
   run = function(
      lowerCoords = double(2),
      upperCoords = double(2),
      sourceCoords = double(2)
   ) {
      ## 1.7.1. Specify the return type dimensionality ----
      returnType(double(1))
      ## 1.7.2. Sanity test the inputs ----
      # Ensure that the number of coordinates is valid
      dimCoords <- dim(lowerCoords)[2]
      if(dimCoords <= 0) {
         stop("invalid dimensionality of the coordinate system")
      }
      # Ensure that the number of windows is valid
      numWindows <- dim(lowerCoords)[1]
      if(numWindows <= 0) {
         stop("invalid number of observation windows")
      }
      # Ensure that the number of coordinates is consistent between the inputs
      if(dim(upperCoords)[2] != dimCoords | dim(sourceCoords)[2] != dimCoords) {
         stop("dimensionality of the inputs is not consisitent")
      }
      if(dim(upperCoords)[1] != numWindows) {
         stop("number of observation windows in the inputs is not consistent")
      }
      # Ensure that the number of points to get the indeces for is valid
      numPoints <- dim(sourceCoords)[1]
      if(numPoints <= 0) {
         stop("invalid number of source points")
      }
      ## 1.7.3. Calculate the indeces ----
      # Initialise a vector of output indeces
      outIndeces <- rep(0, numPoints)
      # Iterate over each of the source points
      for(pointIter in 1:numPoints) {
         # Initialise a flag denoting whether the corrct index has been found
         notFound <- 1
         # Iterate over the observation windows
         while(outIndeces[pointIter] < numWindows & notFound == 1) {
            outIndeces[pointIter] <- outIndeces[pointIter] + 1
            numAboveLowerCoords <- sum(sourceCoords[pointIter, 1:dimCoords] >= lowerCoords[outIndeces[pointIter], 1:dimCoords])
            numBelowUpperCoords <- sum(sourceCoords[pointIter, 1:dimCoords] < upperCoords[outIndeces[pointIter], 1:dimCoords])
            # Test to see if the source point falls within the current observation window
            if(numAboveLowerCoords == dimCoords & numBelowUpperCoords == dimCoords) {
               notFound <- 0
            }
         }
         if(notFound == 1) {
            # Index not found: the source point does not fall within an observation window
            outIndeces[pointIter] <- 0
         }
      }
      return(outIndeces)
   }
)

### 1.8. ==== Function to reduce the number of observation windows further than a given distance from a source coordinate ====
# Reduce the number of observation windows according to a distance criterion (can be used to speed up integration calculations)
obsWindowReduce <- nimbleFunction(
   run = function(
      lowerCoords = double(2),
      upperCoords = double(2),
      sourceCoords = double(1),
      maxDist = double(0),
      areAreas = double(0, default = 1)
   ) {
      ## 1.8.1. Specify the return type dimensionality ----
      returnType(double(2))
      ## 1.8.2. Sanity test the inputs ----
      # Ensure the number of coordinates
      dimCoords <- dim(lowerCoords)[2]
      if(dimCoords <= 0) {
         stop("invalid dimensionality of the coordinate system")
      }
      # Ensure that the number of windows is valid
      numWindows <- dim(lowerCoords)[1]
      if(numWindows <= 0) {
         stop("invalid number of observation windows")
      }
      # Ensure that the number of coordinates is consistent between the inputs
      if(dim(upperCoords)[2] != dimCoords | length(sourceCoords) != dimCoords) {
         stop("dimensionality of the inputs is not consisitent")
      }
      if(dim(upperCoords)[1] != numWindows) {
         stop("number of observation windows in the inputs is not consistent")
      }
      ## 1.8.3. Perform distance check ----
      # Initialise a usage vector
      useRow <- integer(value = c(1), length = numWindows)
      if(maxDist > 0.0) {
         # Calculate the distance from the source coordinate to the nearest point on the observation window
         nearestPoint <- nearestObsWindowPoint(lowerCoords, upperCoords, sourceCoords, areAreas)
         # Iterate over the nearest points and see if they fall within the Euclidean distance
         for(windowIter in 1:numWindows) {
            # Calculate the Euclidean distance
            distVal <- pow(sum(pow(sourceCoords[1:dimCoords] - nearestPoint[windowIter, 1:dimCoords], 2.0)), 0.5)
            # Check to see if the distance is less than the maximum-permissable distance
            if(distVal > maxDist) {
               useRow[windowIter] <- 0
            }
         }
      }
      ## 1.8.4. Create the reduced observation window matrix ----
      # Get the number of rows within the maximum specified distance
      numNewWindows <- sum(useRow)
      # Initalise an output matrix
      outputMat <- matrix(value = 0, nrow = numNewWindows, ncol = dimCoords * 2)
      newWindowIter <- 0
      # Fill the initialised matrix with the rows that fall within the designated distance
      for(windowIter in 1:numWindows) {
         if(useRow[windowIter]) {
            # If the row is to be used then copy the lower and upper coordinates into the relevant places
            # of the matrix
            newWindowIter <- newWindowIter + 1
            outputMat[newWindowIter, 1:dimCoords] <- lowerCoords[windowIter, 1:dimCoords]
            outputMat[newWindowIter, dimCoords + 1:dimCoords] <- upperCoords[windowIter, 1:dimCoords]
         }
      }
      # Return the output matrix
      return(outputMat)
   }
)

## 2. ------ DEFINE A NIMBLE CUSTOM DISTRIBUTION FOR THE (IN)HOMOGENOUS BINOMIAL POINT PROCESS ------

### 2.1. ==== Define the density function ====
# Define a function for the density function for the (in)homogenous binomial process
dbinomPP <- nimbleFunction(
   run = function(
      x = double(2),                               # Coordinate values to calculate the density
      numPoints = double(0),                       # The number of points in the binomial process
      lowerCoords = double(2),                     # The lower coordinate values of the observation windows
      upperCoords = double(2),                     # The upper coordinate values of the observation windows
      intensityWeights = double(1, default = 1),   # Values used in the intensity surface (by default a homogeous process is produced)
      areAreas = double(0, default = 1),           # Flag denoting whether the lower and upper coordinates are areas or transects
      numWindows = double(0, default = -1),        # Number of observation windows (if negative the number of rows in lowerCoords is used to define this value)
      log = integer(0, default = 0)                # If not 0 then return the log density
   ) {
      # Maximum allowable tolerance for testing to see if points fall on lines (only used when areAreas = 0)
      tolDist <- 6.661338e-16
      ## 2.1.1. Specify the return type dimensionality ----
      # Return type declaration
      returnType(double(0))
      ## 2.1.2. Sanity test the inputs ----
      # Assess the dimensionality of the input coordinates
      dimCoords <- dim(x)[2]
      # Ensure that the dimensionality is valid
      if(dimCoords <= 0) {
         stop("invalid dimension structure for the input coordinates")
      }
      # Ensure that the number of points corresponds to the dimensionality of the input coordinates
      if(dim(x)[1] < numPoints) {
         # If the number of rows in the output is less than the number of points simulated by the
         # binomial process: return a zero likelihood.  This is often not required as in nearly every
         # application of the binomial process these values will be the same but we need to handle the
         # occasional exception
         if(log) {
            return(-Inf)
         } else {
            return(0.0)
         }
      }
      if(numPoints == 0) {
         # If no points are simulated then the output is certain
         if(log) {
            return(0.0)
         } else {
            return(1.0)
         }
      } else if(numPoints < 0) {
         # Invalid number of points: set likelihood to zero
         if(log) {
            return(-Inf)
         } else {
            return(0.0)
         }
      }
      # Assess the number of observation windows
      numObsWindows <- trunc(numWindows)
      if(numObsWindows <= 0) {
         numObsWindows <- dim(lowerCoords)[1]
      }
      # Ensure that the number of observation windows is valid
      if(numObsWindows <= 0) {
         stop("invalid number of observation windows")
      }
      # Check that the number of observation windows is consistent (lower coordinates)
      if(numObsWindows > dim(lowerCoords)[1]) {
         stop("number of observation windows not consistent between lower and upper coordinates (or the 'number of windows' parameter)")
      }
      # Check that the number of observation windows is consistent (upper coordinates)
      if(numObsWindows > dim(upperCoords)[1]) {
         stop("number of observation windows not consistent between lower and upper coordinates (or the 'number of windows' parameter)")
      }
      if(dim(lowerCoords)[2] != dimCoords | dim(upperCoords)[2] != dimCoords) {
         stop("lower and/or upper coordinates have an incorrect dimension structure")
      }
      # Calculate the area/length of the observation windows
      obsWindowSize <- calcObsWindowSize(
         matrix(lowerCoords[1:numObsWindows, 1:dimCoords], nrow = numObsWindows, ncol = dimCoords),
         matrix(upperCoords[1:numObsWindows, 1:dimCoords], nrow = numObsWindows, ncol = dimCoords),
         areAreas)
      # Recycle the intensity weights to match the number of observation windows
      recIntensityWeights <- numeric(length = numObsWindows, value = intensityWeights, recycle = TRUE)
      # Ensure that the intensity weights are valid
      if(sum(recIntensityWeights < 0.0) > 0) {
         # Invalid values for the intensity weights: set likelihood to zero
         if(log) {
            return(-Inf)
         } else {
            return(0.0)
         }
      }
      # Find the sum of the product of the intensity weights and area
      sumIntensity <- sum(recIntensityWeights * obsWindowSize)
      if(sumIntensity <= 0.0) {
         # Invalid values for the intensity weights: set likelihood to zero
         if(log) {
            return(-Inf)
         } else {
            return(0.0)
         }
      }
      ## 2.1.3. Calculate the likelihood of each point ----
      logPointDens <- rep(0.0, numPoints)
      for(pointIter in 1:numPoints) {
         curCoords <- x[pointIter, 1:dimCoords]
         # Retrieve the observation windows which the point falls within
         isInObsWindow <- isWithinWindow(curCoords,
            matrix(lowerCoords[1:numObsWindows, 1:dimCoords], nrow = numObsWindows, ncol = dimCoords),
            matrix(upperCoords[1:numObsWindows, 1:dimCoords], nrow = numObsWindows, ncol = dimCoords),
            tolDist, areAreas)
         # Calculate the sum of the intensity
         pointSumIntensity <- sum(isInObsWindow * recIntensityWeights)
         if(pointSumIntensity <= 0.0) {
            if(log) {
               # Return the log scale density if requested
               return(-Inf)
            } else {
               # Return the natural scale density if requested
               return(0.0)
            }
         }
         logPointDens[pointIter] <- log(pointSumIntensity)
      }
      ## 2.1.4. Return the retrieved density ----
      outProb <- sum(logPointDens) - numPoints * log(sumIntensity)
      if(log == 0) {
         # Export the output probability on the natural scale if requested
         outProb <- exp(outProb)
      }
      return(outProb)
   }
)

### 2.2. ==== Define the sampling function ====
# Define a function to draw random coordinates from an (in)homogenous binomial process
rbinomPP <- nimbleFunction(
   run = function(
      n = integer(0),                              # Number of samples to draw from the distribution
      numPoints = double(0),                       # The number of points in the binomial process
      lowerCoords = double(2),                     # The lower coordinate values of the observation windows
      upperCoords = double(2),                     # The upper coordinate values of the observation windows
      intensityWeights = double(1, default = 1),   # Values used in the intensity surface (by default a homogeous process is produced)
      areAreas = double(0, default = 1),           # Flag denoting whether the lower and upper coordinates are areas or transects
      numWindows = double(0, default = -1)         # Number of observation windows (if negative the number of rows in lowerCoords is used to define this value)
   ) {
      ## 2.2.1. Specify the return type dimensionality ----
      # Return type declaration
      returnType(double(2))
      ## 2.2.2. Sanity test the inputs ----
      # Ensure that only one sample is requested
      if(n <= 0) {
         stop("the number of requested samples must be above zero")
      } else if(n > 1) {
         print("rbinomPP only allows n = 1; using n = 1")
      }
      # Assess the dimensionality of the input coordinates
      dimCoords <- dim(lowerCoords)[2]
      # Ensure that the dimensionality is valid
      if(dimCoords <= 0) {
         stop("invalid dimension structure for the input coordinates")
      }
      # Test the values for the number of points
      if(numPoints < 0) {
         stop("invalid number of points to simulate")
      } else if(numPoints == 0) {
         # Return an empty matrix (0 rows and dimCoords columns)
         return(matrix(nrow = 0, ncol = dimCoords, type = "double"))
      }
      # Asses the number of observation windows
      numObsWindows <- trunc(numWindows)
      if(numObsWindows <= 0) {
         numObsWindows <- dim(lowerCoords)[1]
      }
      # Ensure that the number of observation windows is valid
      if(numObsWindows <= 0) {
         stop("invalid number of observation windows")
      }
      # Check that the number of observation windows is consistent (lower coordinates)
      if(numObsWindows > dim(lowerCoords)[1]) {
         stop("number of observation windows not consistent between lower and upper coordinates (or the 'number of windows' parameter)")
      }
      # Check that the number of observation windows is consistent (upper coordinates)
      if(numObsWindows > dim(upperCoords)[1]) {
         stop("number of observation windows not consistent between lower and upper coordinates (or the 'number of windows' parameter)")
      }
      if(dim(upperCoords)[2] != dimCoords) {
         stop("lower and/or upper coordinates have an incorrect dimension structure")
      }
      # Calculate the area/length of the observation windows
      obsWindowSize <- calcObsWindowSize(
         matrix(lowerCoords[1:numObsWindows, 1:dimCoords], nrow = numObsWindows, ncol = dimCoords),
         matrix(upperCoords[1:numObsWindows, 1:dimCoords], nrow = numObsWindows, ncol = dimCoords),
         areAreas)
      # Recycle the intensity weights to match the number of observation windows
      recIntensityWeights <- numeric(length = numObsWindows, value = intensityWeights, recycle = TRUE)
      # Ensure that the intensity weights are valid
      if(sum(recIntensityWeights < 0.0) > 0) {
         recIntensityWeights <- recIntensityWeights * numeric(value = recIntensityWeights > 0.0, length = numObsWindows)
      }
      # Weight the areas by the intensity weights
      areaIntensityWeights <- recIntensityWeights * obsWindowSize
      # Find the sum of the product of the intensity weights and area
      sumIntensity <- sum(areaIntensityWeights)
      if(sumIntensity <= 0.0) {
         # If the area weights sum to zero then instead set them all to their relative sizee
         areaIntensityWeights <- obsWindowSize
         sumIntensity <- sum(areaIntensityWeights)
      }
      ## 2.2.3. Generate random points ----
      # Generate observation window indeces for the output points
      # Currently this can't be done as a single call to rcat due to the way NIMBLE implements the categorical distribution
      obsWindowInd <- rep(0, numPoints)
      for(indIter in 1:numPoints) {
         obsWindowInd[indIter] <- rcat(1, areaIntensityWeights)
      }
      # Initialise an output matrix for the coordinates
      outCoordinates <- matrix(nrow = numPoints, ncol = dimCoords)
      if(areAreas){
         # The observation windows are areas/volumes so generate a set of random
         # coordinates within these volumes
         for(pointIter in 1:numPoints) {
            for(dimIter in 1:dimCoords) {
               outCoordinates[pointIter, dimIter] <- runif(1, min = lowerCoords[obsWindowInd[pointIter], dimIter], max = upperCoords[obsWindowInd[pointIter], dimIter])
            }
         }
      } else {
         # The observation windows are transects so generate a set of random coordinates
         # along their lengths
         for(pointIter in 1:numPoints) {
            coordProp <- runif(1, min = 0, max = 1)
            for(dimIter in 1:dimCoords) {
               outCoordinates[pointIter, dimIter] <- lowerCoords[obsWindowInd[pointIter], dimIter] + 
                  (upperCoords[obsWindowInd[pointIter], dimIter] - lowerCoords[obsWindowInd[pointIter], dimIter]) * coordProp
            }
         }
      }
      ## 2.2.4. Return the generated coordinates ----
      return(outCoordinates)
   }
)

### 2.3. ==== Define the density function for the single data-point version ====
dbinomPPSingle <- nimbleFunction(
	run = function(
		x = double(1),                               # Coordinate values to calculate the density
		lowerCoords = double(2),                     # The lower coordinate values of the observation windows
		upperCoords = double(2),                     # The upper coordinate values of the observation windows
		intensityWeights = double(1, default = 1),   # Values used in the intensity surface (by default a homogeous process is produced)
		areAreas = double(0, default = 1),           # Flag denoting whether the lower and upper coordinates are areas or transects
		numWindows = double(0, default = -1),        # Number of observation windows (if negative the number of rows in lowerCoords is used to define this value)
		log = integer(0, default = 0)                # If not 0 then return the log density
	) {
		## 2.3.1. Specify the return type dimensionality ----
		returnType(double(0))
		## 2.3.2. Create a temporary input matrix ----
		temporaryInput <- matrix(x, ncol = length(x), nrow = 1)
		## 2.3.3. Call the matrix-version of dbinomPP ----
		return(dbinomPP(temporaryInput, 1, lowerCoords, upperCoords, intensityWeights, areAreas, numWindows, log))
	}
)

### 2.4. ==== Define the sampling function for the single data-point version ====
rbinomPPSingle <- nimbleFunction(
	run = function(
		n = integer(0),                              # Number of samples to draw from the distribution
		lowerCoords = double(2),                     # The lower coordinate values of the observation windows
		upperCoords = double(2),                     # The upper coordinate values of the observation windows
		intensityWeights = double(1, default = 1),   # Values used in the intensity surface (by default a homogeous process is produced)
		areAreas = double(0, default = 1),           # Flag denoting whether the lower and upper coordinates are areas or transects
		numWindows = double(0, default = -1)         # Number of observation windows (if negative the number of rows in lowerCoords is used to define this value)
	) {
		## 2.4.1. Specify the return dimensionality ----
		returnType(double(1))
		## 2.4.2. Create a temporary output matrix ----
		# Retrieve the number of dimensions
		dimCoords <- dim(lowerCoords)[2]
		# Call the matrix-version of rbinomPP
		temporaryOutput <- rbinomPP(n, 1, lowerCoords, upperCoords, intensityWeights, areAreas, numWindows)
		## 2.4.3. Return a slice of the output matrix ----
		return(temporaryOutput[1, 1:dimCoords])
	}
)

## 3. ------ DEFINE A NIMBLE CUSTOM DISTRIBUTION FOR THE MULTIVARIATE-NORMAL SOURCE POINT INHOMOGENOUS BINOMIAL POINT PROCESS (AND DERIVATIVES) ------

### 3.1. ==== Define the density function ====
# Define a function for the density function for the multivariate-normal source point inhomogenous binomial process
dbinomMNormSourcePP <- nimbleFunction(
   run = function(
      x = double(2),                               # Coordinate values to calculate the density
      numPoints = double(0),                       # The number of points in the binomial process
      lowerCoords = double(2),                     # The lower coordinate values of the observation windows
      upperCoords = double(2),                     # The upper coordinate values of the observation windows
      sourceCoords = double(1),                    # The coordinates of the source location (the origin of the decay kernel)
      normSD = double(0),                          # The standard deviation of the isotropic multivariate normal distribution decay kernel
      intensityWeights = double(1, default = 1),   # Extra intensity weights for the different observation windows (by default a pure multivariate source point is assumed)
      areAreas = double(0, default = 1),           # Flag denoting whether the lower and upper coordinates are areas or transects
      numWindows = double(0, default = -1),        # Number of observation windows (if negative the number of rows in lowerCoords is used to define this value)
      localEvalParam = double(0, default = -1),    # Parameter that controls the maximum distance that an observation window can be from x to be considered as a possible destination (-1 is interpredted as +Inf)
      log = integer(0, default = 0)                # If not 0 then return the log density
   ) {
      # Maximum allowable tolerance for testing to see if points fall on lines (only used when areAreas = 0)
      tolDist <- 6.661338e-16
      ## 3.1.1. Specify the return type dimensionality ----
      # Return type declaration
      returnType(double(0))
      ## 3.1.2. Sanity test the inputs ----
      # Assess the dimensionality of the input coordinates
      dimCoords <- dim(x)[2]
      # Ensure that the dimensionality is valid
      if(dimCoords <= 0) {
         stop("invalid dimension structure for the input coordinates")
      } else if(dimCoords != 2) {
         stop("currently the bivariate-normal source point inhomogenous point process model is only defined for 2-dimensional processes")
      }
      # Ensure that the number of points corresponds to the dimensionality of the input coordinates
      if(dim(x)[1] < numPoints) {
         # If the number of rows in the output is lower than the number of points simulated by the
         # binomial process: return a zero likelihood.  This is often not required as in nearly every
         # application of the binomial process these values will be the same but we need to handle the
         # occasional
         if(log) {
            # Return the log scale density if requested
            return(-Inf)
         } else {
            # Return the natural scale density if requested
            return(0.0)
         }
      }
      if(numPoints == 0) {
         # If no points are simulated then the output is certain
         if(log) {
            # Return the log scale density if requested
            return(0.0)
         } else {
            # Return the natural scale density if requested
            return(1.0)
         }
      } else if(numPoints < 0) {
         # Invalid number of points: set likelihood to zero
         if(log) {
            # Return the log scale density if requested
            return(-Inf)
         } else {
            # Return the natural scale density if requested
            return(0.0)
         }
      }
      # Asses the number of observation windows
      numObsWindows <- trunc(numWindows)
      if(numObsWindows <= 0) {
         numObsWindows <- dim(lowerCoords)[1]
      }
      # Ensure that the number of observation windows is valid
      if(numObsWindows <= 0) {
         stop("invalid number of observation windows")
      }
      # Check that the number of observation windows is consistent (lower coordinates)
      if(numObsWindows > dim(lowerCoords)[1]) {
         stop("number of observation windows not consistent between lower and upper coordinates (or the 'number of windows' parameter)")
      }
      # Check that the number of observation windows is consistent (upper coordinates)
      if(numObsWindows > dim(upperCoords)[1]) {
         stop("number of observation windows not consistent between lower and upper coordinates (or the 'number of windows' parameter)")
      }
      # Ensure that the lower and upper coordinates have the correct dimension structure
      if(dim(lowerCoords)[2] != dimCoords | dim(upperCoords)[2] != dimCoords) {
         stop("lower and/or upper coordinates have an incorrect dimension structure")
      }
      # Perform the window reduction according to the local evaluation criterion
      redMatrix <- obsWindowReduce(lowerCoords, upperCoords, sourceCoords, localEvalParam, areAreas)
      numObsWindows <- dim(redMatrix)[1]
      if(numObsWindows <= 0) {
         # If there are no observation windows after the reduction process then return a zero likelihood
         if(log) {
            # Return the log scale density if requested
            return(-Inf)
         } else {
            # Return the natural scale density if requested
            return(0.0)
         }
      }
      inLowerCoords <- matrix(nrow = numObsWindows, ncol = dimCoords)
      inUpperCoords <- matrix(nrow = numObsWindows, ncol = dimCoords)
      inLowerCoords[1:numObsWindows, 1:dimCoords] <- redMatrix[1:numObsWindows, 1:dimCoords]                              # Retrieve the lower coordinates of the reduced windows
      inUpperCoords[1:numObsWindows, 1:dimCoords] <- redMatrix[1:numObsWindows, (dimCoords + 1):(dimCoords + dimCoords)]  # Retrieve the upper coordinates of the reduced windows
      # Recycle the intensity weights to match the number of observation windows
      recIntensityWeights <- numeric(length = numObsWindows, value = intensityWeights, recycle = TRUE)
      # Ensure that the intensity weights are valid
      if(sum(recIntensityWeights < 0.0) > 0) {
         # Invalid values for the intensity weights: set likelihood to zero
         if(log) {
            # Return the log scale density if requested
            return(-Inf)
         } else {
            # Return the natural scale density if requested
            return(0.0)
         }
      }
      # Test the validity of the source coordinates
      if(length(sourceCoords) != dimCoords) {
         stop("dimensioality of source coordinates are not consistent")
      }
      # Test the validiity of the decay parameter
      if(normSD <= 0.0) {
         if(log) {
            # Return the log scale density if requested
            return(-Inf)
         } else {
            # Return the natural scale density if requested
            return(0.0)
         }
      }
      ## 3.1.3. Calculate the likelihood of each point ----
      # Calculate the integration of the decay kernel (weighted by intensity) for each observation window
      obsWindowNorm <- calcMNormSourceInt(
         matrix(inLowerCoords[1:numObsWindows, 1:dimCoords], nrow = numObsWindows, ncol = dimCoords),
         matrix(inUpperCoords[1:numObsWindows, 1:dimCoords], nrow = numObsWindows, ncol = dimCoords),
         sourceCoords, normSD, areAreas) * recIntensityWeights
      totalNorm <- sum(obsWindowNorm)
      logPointDens <- rep(0.0, numPoints)
      if(totalNorm <= 0.0) {
         # Return a zero likelihood if the intensity values sum to zero
         if(log) {
            # Return the log scale density if requested
            return(-Inf)
         } else {
            # Return the natural scale density if requested
            return(0.0)
         }
      } else {
         # Calculate the likelihood for each point
         for(pointIter in 1:numPoints) {
            curCoords <- x[pointIter, 1:dimCoords]
            # Retrieve the observation windows which the point falls within
            isInObsWindow <- isWithinWindow(curCoords,
               matrix(inLowerCoords[1:numObsWindows, 1:dimCoords], nrow = numObsWindows, ncol = dimCoords),
               matrix(inUpperCoords[1:numObsWindows, 1:dimCoords], nrow = numObsWindows, ncol = dimCoords),
               tolDist, areAreas)
            # Calculate the distance decay component of the current point from the source (the exponent
            # of the isotropic multivariate normal diistribution)
            distDecay <- 0.0
            for(dimIter in 1:dimCoords) {
               distDecay <- distDecay + (curCoords[dimIter] - sourceCoords[dimIter]) * (curCoords[dimIter] - sourceCoords[dimIter])
            }
            distDecay <- distDecay * (-0.5 / (normSD * normSD))
            # Calculate the sum of the intensity for each point
            pointSumIntensity <- sum(isInObsWindow * recIntensityWeights * exp(distDecay))
            if(pointSumIntensity <= 0.0) {
               if(log) {
                  # Return the log scale density if requested
                  return(-Inf)
               } else {
                  # Return the natural scale density if requested
                  return(0.0)
               }
            }
            logPointDens[pointIter] <- log(pointSumIntensity)
         }
      }
      ## 3.1.4. Return the retrieved density ----
      outProb <- sum(logPointDens) - numPoints * log(totalNorm)
      if(log == 0) {
         # Export the output probability on the natural scale if requested
         outProb <- exp(outProb)
      }
      return(outProb)
   }
)

### 3.2. ==== Define the sampling function ====
# Define a function to draw random coordinate from the multivariate-normal source point inhomogenous binomial process
rbinomMNormSourcePP <- nimbleFunction(
   run = function(
      n = integer(0),                              # Number of samples to draw from the distribution
      numPoints = double(0),                       # The number of points in the binomial process
      lowerCoords = double(2),                     # The lower coordinate values of the observation windows
      upperCoords = double(2),                     # The upper coordinate values of the observation windows
      sourceCoords = double(1),                    # The coordinates of the source location (the origin of the decay kernel)
      normSD = double(0),                          # The standard deviation of the isotropic multivariate normal distribution decay kernel
      intensityWeights = double(1, default = 1),   # Extra intensity weights for the different observation windows (by default a pure multivariate source point is assumed)
      areAreas = double(0, default = 1),           # Flag denoting whether the lower and upper coordinates are areas or transects
      numWindows = double(0, default = -1),        # Number of observation windows (if negative the number of rows in lowerCoords is used to define this value)
      localEvalParam = double(0, default = -1)     # Parameter that controls the maximum distance that an observation window can be from x to be considered as a possible destination (-1 is interpredted as +Inf)
   ) {
      ## 3.2.1. Specify the return type dimensionality ----
      # Return type declaration
      returnType(double(2))
      ## 3.2.2. Sanity test the inputs ----
      # Ensure that only one sample is requested
      if(n <= 0) {
         stop("the number of requested samples must be above zero")
      } else if(n > 1) {
         print("rbinomMNormSourcePP only allows n = 1; using n = 1")
      }
      # Assess the dimensionality of the input coordinates
      dimCoords <- dim(lowerCoords)[2]
      # Ensure that the dimensionality is valid
      if(dimCoords <= 0) {
         stop("invalid dimension structure for the input coordinates")
      }
      # Test the values for the number of points
      if(numPoints < 0) {
         stop("invalid number of points to simulate")
      } else if(numPoints == 0) {
         # Return an empty matrix (0 rows and dimCoords columns)
         return(matrix(nrow = 0, ncol = dimCoords, type = "double"))
      }
      # Assess the number of observation windows
      numObsWindows <- trunc(numWindows)
      if(numObsWindows <= 0) {
         numObsWindows <- dim(lowerCoords)[1]
      }
      # Ensure that the number of observation windows is valid
      if(numObsWindows <= 0) {
         stop("invalid number of observation windows")
      }
      # Check that the number of observation windows is consistent (lower coordinates)
      if(numObsWindows > dim(lowerCoords)[1]) {
         stop("number of observation windows not consistent between lower and upper coordinates (or the 'number of windows' parameter)")
      }
      # Check that the number of observation windows is consistent (upper coordinates)
      if(numObsWindows > dim(upperCoords)[1]) {
         stop("number of observation windows not consistent between lower and upper coordinates (or the 'number of windows' parameter)")
      }
      if(dim(upperCoords)[2] != dimCoords) {
         stop("lower and/or upper coordinates have an incorrect dimension structure")
      }
      # Recycle the intensity weights to match the number of observation windows
      recIntensityWeights <- numeric(length = numObsWindows, value = intensityWeights, recycle = TRUE)
      # Ensure that the intensity weights are valid
      if(sum(recIntensityWeights < 0.0) > 0) {
         recIntensityWeights <- recIntensityWeights * numeric(value = recIntensityWeights > 0.0, length = numObsWindows)
      }
      # Test the validity of the source coordinates
      if(length(sourceCoords) != dimCoords) {
         stop("dimensionality of source coordinates are not consistent")
      }
      # Test the validiity of the decay parameter
      if(normSD <= 0.0) {
         stop("invalid values for the decay parameter")
      }
      # Print error message if the local evaluation parameters are used in the simulation function
      if(localEvalParam > 0.0) {
         print("local evaluation parameters not used in the simulation function")
      }
      ## 3.2.3. Generate random points ----
      # Current a stratified rejection sampling approach is employed to generate random variables from this process.
      # This can become inefficient if the observation windows because very large compared to the range of the decay
      # kernel.  This could be optimised better in future versions.
      obsWindowInd <- rep(1, numPoints)
      if(numObsWindows > 1) {
         # Weight the observation windows by their intensity weights multiplied by the integral of the decay function
         integratedSource <- calcMNormSourceInt(
            matrix(lowerCoords[1:numObsWindows, 1:dimCoords], nrow = numObsWindows, ncol = dimCoords),
            matrix(upperCoords[1:numObsWindows, 1:dimCoords], nrow = numObsWindows, ncol = dimCoords),
            sourceCoords, normSD, areAreas)
         areaIntensityWeights <- recIntensityWeights * integratedSource
         # Find the sum of the product of the intensity weights and area
         sumIntensity <- sum(areaIntensityWeights)
         if(sumIntensity <= 0.0) {
            areaIntensityWeights <- integratedSource
            sumIntensity <- sum(areaIntensityWeights)
         }
         # Generate observation window indeces for the output
         # Currently this can't be done as a single call to rcat due to the way NIMBLE implements the categorical distribution
         for(indIter in 1:numPoints) {
            obsWindowInd[indIter] <- rcat(1, areaIntensityWeights)
         }
      }
      # Initialise an output matrix for the coordinates
      outCoordinates <- matrix(nrow = numPoints, ncol = dimCoords)
      # Calculate the nearest point to the source coordinates in each observation window
      nearestPoints <- nearestObsWindowPoint(
         matrix(lowerCoords[1:numObsWindows, 1:dimCoords], nrow = numObsWindows, ncol = dimCoords),
         matrix(upperCoords[1:numObsWindows, 1:dimCoords], nrow = numObsWindows, ncol = dimCoords),
         sourceCoords, areAreas)
      # Calculate the value of the decay kernel at the nearest point in each observation window
      decayNearest <- rep(0.0, numObsWindows)
      for(dimIter in 1:dimCoords) {
         decayNearest <- decayNearest + (nearestPoints[1:numObsWindows, dimIter] - sourceCoords[dimIter]) * (nearestPoints[1:numObsWindows, dimIter] - sourceCoords[dimIter])
      }
      decayNearest <- (decayNearest * -0.5) / (normSD * normSD)
      decayNearest <- exp(decayNearest)
      if(areAreas) {
         # The observation windows are areas/volumes so generate a set of random
         # coordinates within these volumes
         for(pointIter in 1:numPoints) {
            # Create a set of test coordinates
            testCoords <- runif(dimCoords, min = lowerCoords[obsWindowInd[pointIter], 1:dimCoords], max = upperCoords[obsWindowInd[pointIter], 1:dimCoords])
            # Assess the value of the decay parameter
            decayTest <- (testCoords - sourceCoords) * (testCoords - sourceCoords)
            decayVal <- exp((-0.5 * sum(decayTest)) / (normSD * normSD))
            randVal <- runif(1, min = 0, max = 1)
            while(randVal > (decayVal / decayNearest[obsWindowInd[pointIter]])) {
               # Create a set of test coordinates
               testCoords <- runif(dimCoords, min = lowerCoords[obsWindowInd[pointIter], 1:dimCoords], max = upperCoords[obsWindowInd[pointIter], 1:dimCoords])
               # Assess the value of the decay parameter
               decayTest <- (testCoords - sourceCoords) * (testCoords - sourceCoords)
               decayVal <- exp((-0.5 * sum(decayTest)) / (normSD * normSD))
               randVal <- runif(1, min = 0, max = 1)
            }
            # Save the test coordinates once they have been accepted
            outCoordinates[pointIter, 1:dimCoords] <- testCoords
         }
      } else {
         # The observation windows are transects so generate a set of random coordinates
         # along their lengths
         for(pointIter in 1:numPoints) {
            # Create a set of test coordinates
            testProp <- runif(1, min = 0, max = 1)
            testCoords <- lowerCoords[obsWindowInd[pointIter], 1:dimCoords] + (upperCoords[obsWindowInd[pointIter], 1:dimCoords] - lowerCoords[obsWindowInd[pointIter], 1:dimCoords]) * testProp
            # Assess the value of the decay parameter
            decayTest <- (testCoords - sourceCoords) * (testCoords - sourceCoords)
            decayVal <- exp((-0.5 * sum(decayTest)) / (normSD * normSD))
            randVal <- runif(1, min = 0, max = 1)
            while(randVal > (decayVal / decayNearest[obsWindowInd[pointIter]])) {
               # Create a set of test coordinates
               testProp <- runif(1, min = 0, max = 1)
               testCoords <- lowerCoords[obsWindowInd[pointIter], 1:dimCoords] + (upperCoords[obsWindowInd[pointIter], 1:dimCoords] - lowerCoords[obsWindowInd[pointIter], 1:dimCoords]) * testProp
               # Assess the value of the decay parameter
               decayTest <- (testCoords - sourceCoords) * (testCoords - sourceCoords)
               decayVal <- exp((-0.5 * sum(decayTest)) / (normSD * normSD))
               randVal <- runif(1, min = 0, max = 1)
            }
            # Save the test coordinates once they have been accepted
            outCoordinates[pointIter, 1:dimCoords] <- testCoords
         }
      }
      ## 3.2.4. Return the generated coordinates ----
      return(outCoordinates)
   }
)

### 3.3. ==== Define the density function for the single data-point version ====
dbinomMNormSourcePPSingle <- nimbleFunction(
   run = function(
      x = double(1),                               # Coordinate values to calculate the density
      lowerCoords = double(2),                     # The lower coordinate values of the observation windows
      upperCoords = double(2),                     # The upper coordinate values of the observation windows
      sourceCoords = double(1),                    # The coordinates of the source location (the origin of the decay kernel)
      normSD = double(0),                          # The standard deviation of the isotropic multivariate normal distribution decay kernel
      intensityWeights = double(1, default = 1),   # Extra intensity weights for the different observation windows (by default a pure multivariate source point is assumed)
      areAreas = double(0, default = 1),           # Flag denoting whether the lower and upper coordinates are areas or transects
      numWindows = double(0, default = -1),        # Number of observation windows (if negative the number of rows in lowerCoords is used to define this value)
      localEvalParam = double(0, default = -1),    # Parameter that controls the maximum distance that an observation window can be from x to be considered as a possible destination (-1 is interpredted as +Inf)
      log = integer(0, default = 0)                # If not 0 then return the log density
   ) {
		## 3.3.1. Specify the return type dimensionality ----
      returnType(double(0))
      ## 3.3.2. Create a temporary input matrix ----
      temporaryInput <- matrix(x, ncol = length(x), nrow = 1)
      ## 3.3.3. Call the matrix-version of dbinomMNormSourcePP ----
      return(dbinomMNormSourcePP(temporaryInput, 1, lowerCoords, upperCoords, sourceCoords, normSD, intensityWeights, areAreas, numWindows, localEvalParam, log))
   }
)

### 3.4. ==== Define the sampling function for the single data-point version ====
rbinomMNormSourcePPSingle <- nimbleFunction(
   run = function(
      n = integer(0),                              # Number of samples to draw from the distribution
      lowerCoords = double(2),                     # The lower coordinate values of the observation windows
      upperCoords = double(2),                     # The upper coordinate values of the observation windows
      sourceCoords = double(1),                    # The coordinates of the source location (the origin of the decay kernel)
      normSD = double(0),                          # The standard deviation of the isotropic multivariate normal distribution decay kernel
      intensityWeights = double(1, default = 1),   # Extra intensity weights for the different observation windows (by default a pure multivariate source point is assumed)
      areAreas = double(0, default = 1),           # Flag denoting whether the lower and upper coordinates are areas or transects
      numWindows = double(0, default = -1),        # Number of observation windows (if negative the number of rows in lowerCoords is used to define this value)
      localEvalParam = double(0, default = -1)     # Parameter that controls the maximum distance that an observation window can be from x to be considered as a possible destination (-1 is interpredted as +Inf)
   ) {
      ## 3.4.1. Specify the return type dimensionality ----
      returnType(double(1))
      ## 3.4.2. Sanity test the inputs ----
      # Ensure that only one sample is requested
      if(n <= 0) {
         stop("the number of requested samples must be above zero")
      } else if(n > 1) {
         print("rbinomMNormSourcePPSingle only allows n = 1; using n = 1")
      }
      ## 3.4.3. Create a temporary output matrix ----
      # Retrieve the number of coordinates
      dimCoords <- dim(lowerCoords)[2]
      # Call the matrix-version of rbinomMNormSourcePP
      temporaryOutput <- rbinomMNormSourcePP(1, 1, lowerCoords, upperCoords, sourceCoords, normSD, intensityWeights, areAreas, numWindows, localEvalParam)
      ## 3.4.3. Return a slice of the output matrix ----
      return(temporaryOutput[1, 1:dimCoords])
   }
)


## 6. ------ REGISTER THE DISTRIBUTIONS ------
# Register the distributions with NIMBLE
registerDistributions(list(
   ### 6.1. ==== Register the dbinomPP distribution ====
   dbinomPP = list(
      ## 6.1.1. Define the BUGS code to call the distribution ----
      BUGSdist = "dbinomPP(numPoints, lowerCoords, upperCoords, intensityWeights, areAreas, numWindows)",
      ## 6.1.2. Set the input and output types and dimension structure ----
      types = c(
         "value = double(2)", "numPoints = double(0)", "lowerCoords = double(2)",
         "upperCoords = double(2)", "intensityWeights = double(1)", "areAreas = double(0)", "numWindows = double(0)"),
      ## 6.1.3. Define the cumulative probability and quantile function availability ----
      pqAvail = FALSE
   ),
   ### 6.2. ==== Register the dbinomPPSingle distribution ====
   dbinomPPSingle = list(
      ## 6.2.1. Define the BUGS code to call the distribution ----
      BUGSdist = "dbinomPPSingle(lowerCoords, upperCoords, intensityWeights, areAreas, numWindows)",
      ## 6.2.2. Set the input and output types and dimension structure ----
      types = c(
         "value = double(1)", "lowerCoords = double(2)", "upperCoords = double(2)",
         "intensityWeights = double(1)", "areAreas = double(0)", "numWindows = double(0)"),
      ## 6.2.3. Define the cumulative probability and quantile function availability ----
      pqAvail = FALSE
   ),
   ### 6.3. ==== Register the dbinomMNormSourcePP distribution ====
   dbinomMNormSourcePP = list(
      ## 6.3.1. Define the BUGS code to call the distribution ----
      BUGSdist = "dbinomMNormSourcePP(numPoints, lowerCoords, upperCoords, sourceCoords, normSD, intensityWeights, areAreas, numWindows, localEvalParam)",
      ## 6.3.2. Set the input and output types and dimension structure ----
      types = c(
         "value = double(2)", "numPoints = double(0)", "lowerCoords = double(2)",
         "upperCoords = double(2)", "sourceCoords = double(1)", "normSD = double(0)",
         "intensityWeights = double(1)", "areAreas = double(0)", "numWindows = double(0)",
         "localEvalParam = double(0)"),
      ## 6.3.3. Define the cumulative probability and quantile function availability ----
      pqAvail = FALSE
   ),
   ### 6.4. ==== Register the dbinomMNormSourcePPSingle distribution ====
   dbinomMNormSourcePPSingle = list(
      ## 6.4.1. Define the BUGS code to call the distribution ----
      BUGSdist = "dbinomMNormSourcePPSingle(lowerCoords, upperCoords, sourceCoords, normSD, intensityWeights, areAreas, numWindows, localEvalParam)",
      ## 6.4.2. Set the input and output types and dimension structure ----
      types = c(
         "value = double(1)", "lowerCoords = double(2)", "upperCoords = double(2)",
         "sourceCoords = double(1)", "normSD = double(0)", "intensityWeights = double(1)",
         "areAreas = double(0)", "numWindows = double(0)", "localEvalParam = double(0)"),
      ## 6.4.3. Define the cumulative probability and quantile function availability ----
      pqAvail = FALSE
   )

))
```
