## Supporting informations for "Estimating abundance with interruptions in data collection using open population spatial capture-recapture models": SourceRFunctions.R

```
## ==== I. SimulateZ ====
#' @title Data set simulation function.
#'
#' @description
#' \code{SimulateZ} returns a \code{list} object with the composition of the simulated population,
#'  its pop size and the corresponding z matrix for CMR models.
#'
#' @param N0 A \code{numeric} object containig the original number of individuals in the pop to be simulated (alternative to z0)
#'	@param n.occasions A \code{numeric} object containing the number of years tobe simulated.
#' @param PHI A \code{matrix} object containing the state-specific survival probabilities.
#' @param REPRO A \code{matrix} object containing the state-specific reproduction probabilities.
#' @param FEC A \code{vector} object containing the state-specific fecundity (= mean litter size).
#' @param INIT.STATE A \code{vector} object containing the initial-state probabilities (= prob to enter the data set in each state).
#' 
#' @return A \code{list} object with the population attributes 
#' pop.ls: a list of population composition for each time step
#' N: a list of pop sizes
#' z: the state matrix to be used in CMR models
#' 

SimulateZ <- function( N0 = NULL
                       , n.occasions
                       , PHI
                       , REPRO
                       , FEC  
                       , init.state = 1)
{
   ## ---- CREATE POPULATION LIST ----
   POP <- POP.SIZE <- list()
   POP[[1]] <- data.frame(id=1:N0, z=init.state)
   
   ## ---- CREATE POPULATION SIZE VECTOR ----
   N <- vector()
   N[1] <- N0
   ## ----CONVERT TO ARRAY AND MATRICES
   if(length(dim(PHI))!=3){
      PHI <- array(PHI, c(dim(PHI),n.occasions))
   }
   if(length(dim(REPRO))!=3){
      REPRO <- array(REPRO, c(dim(REPRO),n.occasions))
   }
   if(length(FEC)==dim(PHI)[1]){
      FEC <- matrix(FEC, nrow=dim(PHI)[1],ncol=n.occasions)
   }
   
   #-----------------------------------------------------------------------
   ## ---- LOOP POPULATION PROCESS OVER n.occasions ----
   for(t in 2:(n.occasions+1)){
      Z <- R <- B <- Z.new <-  NULL
        for (i in 1:(dim(POP[[t-1]])[1])){
            R[i] <- rbinom(1, 1, REPRO[init.state,POP[[t-1]]$z[i], t-1 ])
            if(R[i]==0){B[i] <-  0
            }else{
               B[i] <- rpois(1, FEC[POP[[t-1]]$z[i], t-1 ])
            }
            Z[i] <- which(rmultinom(1, 1, PHI[,POP[[t-1]]$z[i], t-1 ])==1)
         }#i

      
      N.new <- sum(B[])
      if(N.new>=1){
         if(is.null(init.state)){init.state <- c(1,rep(0,dim(PHI)[2]-1))}
         for (i in 1:N.new){
            Z.new[i] <- which(rmultinom(1, 1, init.state)==1)
         }
         Z <- c(Z,Z.new)
      }
      N[t] <- length(Z)
      POP[[t]] <- data.frame(id=1:N[t], z = Z)
   }
   #-----------------------------------------------------------------------
   ## ---- GENERATE z BASED ON POPULATION LIST
   z <- matrix(NA, N[n.occasions+1], n.occasions+1)
   for (t in 1:(n.occasions+1))
   {
      z[1:N[t],t] <- POP[[t]]$z
      pop.size <- vector()
      for (s in 1:dim(PHI)[1]){
         pop.size[s] <- length(which(z[ ,t]==s))
      }
      POP.SIZE[[t]] <- c(pop.size)
   }
   
   #----------------------------------------------------------------------
   return (list(pop.ls = POP, N.stage = POP.SIZE, N.tot=N, z = z))
}

## ==== II. MakeInitsXY ====
#' @title Function to set initial values of XY coordinates of ACS
#'
#' @description
#' \code{MakeInitsXY} returns a matrix object with with the x coordinates ([,1]) and y coordinates  ([,2]). it returns the location of a detection for a detected individual and a random location for augmented individuals.
#' for individuals that remained undetected during one or several years, the detections from before/after are used as the centroid of the buffer (radius determinde by the dist move).
#' 
#' @param y \code{matrix} or \code{array}  with individual detections. row=Ids, col= detectors and if arrray=  [,,t] time. 
#' @param detector.xy \code{matrix} or \code{array} with coordinates of detectors. if array, the 3rd dimension corresponds to years
#' @param habitat.r \code{raster} with the raster of the habitat (0/1)
#' @param dist.move \code{numeric} with a radius value of how much an individual can move from year to year. sxy t+1 will be drawn from a buffer centered on average sxy t
#' @param ydead \code{array}  with individual dead recoveries row=Ids, col= detectors and if arrray=  [,,t] time. 
#' @param detector.xyDead \code{matrix} or \code{array} with coordinates of detectors. if array, the 3rd dimension corresponds to years
#'
#'
#' @examples
#' # Generate simulated detection histories within an SCR framework:
#' MakeInitsXY(   y = y
#'              , detector.xy = data$detector.xy
#'              , habitat.r = habitat.r
#'              , dist.move = dist.move )

MakeInitsXY <- function(    y = y
                           , 
                           detector.xy = detector.xy
                           ,
                           habitat.r =  myHabitat$habitat.r
                           ,
                           dist.move = dist.move
                           ,
                           ydead = NULL
                           ,
                           detector.xyDead = NULL
                           ,
                           seed = FALSE
){
   
   ##PREPARE INPUT  
   n.years <- ifelse(length(dim(y))>=3, dim(y)[3], 1)
   n.individuals <- dim(y)[1]
   n.detectors <- dim(detector.xy)[1]
   if(length(dim(y)) == 2){y <- array(y, c(n.individuals, n.detectors, n.years))}
   if(length(dim(detector.xy)) == 2){detector.xy <- array(detector.xy, c(n.detectors, 2, n.years))}
   sxy <- array(NA, c(n.individuals, 2, n.years))
   
   # CHECK WHO/WHEN IDS ARE DETECTED 
   detected <- apply(y, c(1,3), function(x) any(x >= 1))
   id.detec <- which(apply(detected, 1, function(x) sum(x==FALSE)!=n.years))
   
   if(!is.null(ydead)){
      detected.dead <-  ydead > 0
      id.dead <- which(apply(ydead,1, function(x) any(x > 0)) ==T)
      id.detec <- unique(c(id.detec, id.dead ))
   }
   
   # CREATE THE HABITAT POLYGON
   buffered.habitat.poly <- aggregate(rasterToPolygons(habitat.r, fun=function(x){x>0}))
   crs <- CRS(proj4string(buffered.habitat.poly))
   
   ## DETECTED INDIVIDUALS 
   for(i in id.detec){
      
      # IF AT LEAST ONE DETECTION ALIVE
      if(sum(detected[i,])>0){
         # YEAR WITH DETECTIONS 
         detected.years <- which(detected[i,]==TRUE)
         #GET AVERAGE DETECTIONS
         for(t in detected.years){
            tmpsxy <- detector.xy[which(y[i,,t]>0),,t] 
            sxy[i,,t] <- if(is.null(dim(tmpsxy))){
               tmpsxy }else{ colMeans(tmpsxy)}  
         }
         # YEAR WITH NODETECTIONS
         if(sum(detected[i,])!=n.years){
             notdetected.years <- which(detected[i,]==FALSE)
            
             for(t in notdetected.years){
                if(t < min(detected.years)){
                   sxy[i,,t] <- sxy[i,,min(detected.years)] 
                }
                if(t > max(detected.years)){
                   sxy[i,,t] <- sxy[i,,max(detected.years)]  
                }
                if(t < max(detected.years) & t > min(detected.years) ){
                      sxy[i,,t] <- sxy[i,,t-1]  
                }
                
                 sp <- SpatialPoints(rbind(sxy[i,,t]), proj4string = crs)
                 buffer <- gBuffer(sp, width = dist.move, byid = T )
                 int <- intersect(buffer, buffered.habitat.poly)
                 if((seed)){
                    set.seed(i + seed)
                 }
                 sxy[i,,t] <- t(coordinates(spsample(int, n=1, type="random", iter=500)))
                 
             }
             

            #CALCULATE AVERAGE DETECTIONS OF ALL YEARS 
            # notdetected.years <- which(detected[i,]==FALSE)
            # averagesxy <- sxy[i,,detected.years] 
            # averagesxy <-  if(is.null(dim(averagesxy))){
            #    averagesxy }else{ rowMeans(averagesxy)}  
            # DRAW A BUFFER
            # sp <- SpatialPoints(rbind(averagesxy), proj4string = crs)
            # buffer <- gBuffer(sp, width = dist.move )
            # int <- intersect(buffer, buffered.habitat.poly)
            # DRAW SXY
         
         }
      }
      
      ## IF DEAD
      if(!is.null(ydead)){
         # IF AT LEAST ONE DETECTION DEAD
         if(sum(detected.dead[i,])>0){
            # YEAR WITH DETECTIONS 
            detected.years <- which(detected.dead[i,]==TRUE)
            #GET SXY DEAD 
            for(t in detected.years){
               sxy[i,,t] <- detector.xyDead[ydead[i,t],,t] 
            }
            #IF INDIVIDUAL ONLY DETECTED DEAD 
            if(sum(detected[i,])==0){
               #CALCULATE AVERAGE DETECTIONS OF ALL YEARS 
               notdetected.years <- which((detected[i,]+detected.dead[i,])==0)
               averagesxy <- sxy[i,,detected.years] 
               # DRAW A BUFFER
               sp <- SpatialPoints(rbind(averagesxy), proj4string = crs)
               buffer <- gBuffer(sp, width = dist.move )
               int <- intersect(buffer, buffered.habitat.poly)
               if(!is.null(seed)){
                  set.seed(i + seed)
               }
               # DRAW SXY
               sxy[i,,notdetected.years] <- t(coordinates(spsample(int, n=length(notdetected.years), type="random", iter=500)))
            }
         }
      }
   }
   
   #AUGMENTED IDS 
   id.notdetec <- c(1:n.individuals)[(c(1:n.individuals) %in% id.detec)==FALSE]
   if(length(id.notdetec)!=0){
   start.loca <- spsample(buffered.habitat.poly, n=length(id.notdetec),type="random",iter=500)
   # DRAW A BUFFER
   buffer.augm <- gBuffer(start.loca,width = dist.move, byid = T)
   buffered.habitat.poly <- gBuffer(buffered.habitat.poly, width = -1)
   int.buff <- intersect(buffered.habitat.poly, buffer.augm)
   
   for(i in 1:length(id.notdetec)){
      #DRAW COORDINATES FROM WITHIN THE BUFFER 
      if(!is.null(seed)){
         set.seed(i + seed)
      }
      coords <- spsample(int.buff[i,], n=n.years, type="random",iter=500)
      sxy[id.notdetec[i],,] <- t(coordinates(coords))
      #model$habitat.mx[trunc(thisSXY[2])+1, trunc(thisSXY[1])+1]    ## coming from habitat array
      
   }
   }
   # ADD SOME RANDOM NOISE TO AVOID NON-ZERO MOVEMENT
   for(t in 1:n.years){
      if(!is.null(seed)){
         set.seed(t + seed)
      }
      sxy[,2,t] <-  sxy[,2,t] +jitter(0,factor = 0.01)
   }
   return(sxy)
   
}


#' @title Plot trace plots and posterior density of parameters
#'
#' @description
#' \code{PlotJagsParams} Plot trace plots, posterior density of parameters, and a vertical red line for the true parameter value (if known)
#'
#' @param jags.samples A \code{mcmc.list} object from coda.samples()functions in rjags
#' @param params A \code{vector} of strings with the variable names for plots should be computed
#' @param sim.values A \code{vector} with the true values of the parameters in \code{params}
#' @param trace.plot A \code{logial} if traceplot should be plotted
#' @param density.plot A \code{logial} if posterior density should be plotted
#'
#'
#' @return Plot trace plots and posterior density of parameters  
#' @example 
#' 


## ==== III. PlotJagsParams ====

PlotJagsParams <- function( jags.samples 
                          , params = NULL
                          , sim.values = NULL
                          , trace.plot = TRUE
                          , density.plot = TRUE)
{
if(is.null(params)){params <- colnames(jags.samples[[1]])}
for(i in 1:length(params))
   {
   if(trace.plot & density.plot){par(mfrow=c(1,2))}
   if(trace.plot){
      traceplot(jags.samples[ ,params[i]])
      if(!is.null(sim.values)){abline(h = sim.values[i], col = "black", lwd = 3, lty = 2)}}
   if(density.plot){
      plot(density(unlist(jags.samples[ ,params[i]])), main = params[i])
      if(!is.null(sim.values)){abline(v = sim.values[i], col = "red", lwd = 2)}}
   }
}
```
