## Supporting informations for "Estimating abundance with interruptions in data collection using open population spatial capture-recapture models": SupportingInformation1.docx

**Supporting information 1.1** Description of the point process model

We used a derivation of the homogenous binomial point process for the placement of activity centers (ACs) and it was this formulation that formed the basis for the placement of the ACs in the first time period, *s_i_*_1_. Under this model, each AC is independently and uniformly placed in the available habitat. A natural extension of the homogenous binomial point process is to allow the density of activity centers to vary spatially according to a spatial intensity function, *λ*(***x***), where ***x*** is a vector of spatial coordinates. Under this inhomogeneous extension, the number of ACs falling within an area *U* (here defined as K) follows

$$K\sim\text{Binomial}\left( n,\frac{\Lambda\left( U \right)}{\Lambda\left( S \right)} \right)$$

where *n* is the number of activity centers within the entire region of interest (*S*) and

$$\Lambda\left( U \right)=\iint_{U} \lambda\left( \boldsymbol{x} \right)d\boldsymbol{x}$$

In this study, we define the intensity function of the inhomogeneous binomial process to condition the placement of the activity center of individual *i* at time period *t,* at coordinates $s_{i_{t}}= \left( s_{i_{x,t}},s_{i_{y,t}} \right)$, on the location of its activity center in the previous time period, at coordinates $s_{i_{t-1}}= \left( s_{i_{x,t-1}},s_{i_{y,t-1}} \right),$ such that

$$\lambda\left( s_{i_{t}},\tau,s_{i_{t-1}} \right)=e^{-\frac{1}{2\tau^{2}}\left( s_{i_{x,t}}-s_{i_{x,t-1}} \right)^{2}+\left( s_{i_{y,t}}-s_{i_{y,t-1}} \right)^{2}}$$

Under this specification, the dynamics of the activity center of each individual is modelled as a series of inhomogeneous binomial point processes, each with *n* = 1. The spatial intensity function describes movement as a series of isotropic Gaussian random walks (similar to that applied in (Gardner et al. 2018)) with a standard deviation set equal to *τ*. *τ* therefore regulates the distance that individuals are likely to move between time periods.

**Supporting information 1.2** Maps of the study area

**Figure S1.2.1.** Map of the study area (spatial domain, dark grey polygon) in Norway (A), and annual maps (B-F) representing wolverine detections (black points) and detector locations (red points) within the same spatial domain.

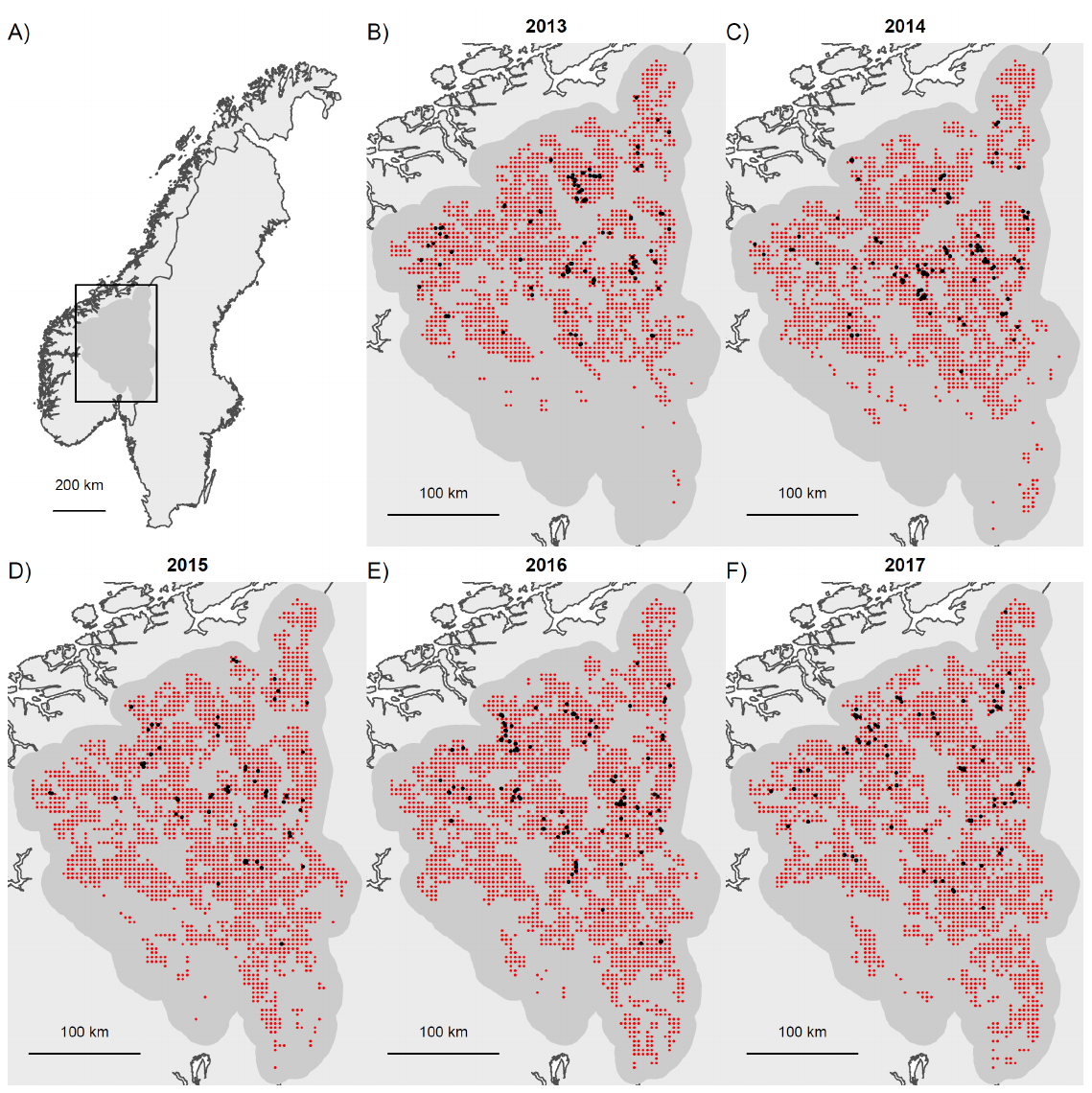

**Supporting information 1.3** List of parameters and priors used

**Table S1.3.1** Description of prior distributions for parameters estimated by the Bayesian OPSCR model.

| **Parameter** | **Description** | **Prior** |
| --- | --- | --- |
| $\tau$ | Standard deviation of the Gaussian dispersal kernels describing the movement of AC locations between years | dgamma(0.01,0.01) |
| $\phi$ | Survival probability | dunif(0,1) |
| ψ | Inclusion probability during the first year | dunif(0,1) |
| $\rho$ | Per capita recruitment | dunif(0,5) |
| σ | Scale parameter of the halfnormal detection function | dunif(0,5) |
| p0 | Baseline detection probability | dunif(0,1) |

**Supporting information 1.4** Summary of the parameter convergence

**Table S1.4.1** Number of models that reached convergence (Rhat ≥1.1) for at least one of the parameters of the OPSCR models (N, σ, ρ,$\phi$, $\tau$, p0). Models that did not converge after 40000 iterations (2000 burn-in) were ran again with 60000 iterations (20000 burn-in). In all cases where a model did not reach convergence after 60000 iterations (49 models), it was due the parameter “dispersal sigma” failing to reach the criterion (Rhat <1.1). OPSCR models were fitted to 50 different data sets that were simulated for each life history and interruption scenario.

|  | **Interruption scenarios** | | | | | | | | | |
| --- | --- | --- | --- | --- | --- | --- | --- | --- | --- | --- |
|  | **1**  **(11111)** | **2**  **(11011)** | **3**  **(10011)** | **4**  **(11001)** | **5**  **(10111)** | **6**  **(11101)** | **7 (10101)** | **8 (01111)** | **9**  **(11110)** | **10**  **(10001)** |
| **Life history** | **Number of models not converged (40000 iterations)** | | | | | | | | | |
| **Slow** | 46 | 47 | 49 | 48 | 47 | 48 | 48 | 47 | 50 | 48 |
| **Slow stochastic** | 50 | 48 | 47 | 46 | 50 | 46 | 46 | 45 | 50 | 38 |
| **Intermediate** | 44 | 48 | 45 | 44 | 46 | 49 | 46 | 44 | 49 | 17 |
| **Intermediate stochastic** | 46 | 46 | 46 | 45 | 46 | 45 | 47 | 46 | 48 | 16 |
| **Number of models not converged (60000 iterations)** | | | | | | | | | | |
| **Slow** | 50 | 50 | 50 | 50 | 50 | 50 | 50 | 50 | 50 | 50 |
| **Slow stochastic** | 50 | 50 | 50 | 50 | 50 | 50 | 50 | 50 | 50 | 50 |
| **Intermediate** | 50 | 50 | 50 | 50 | 50 | 50 | 50 | 50 | 50 | 30 |
| **Intermediate stochastic** | 50 | 50 | 50 | 50 | 50 | 50 | 50 | 50 | 50 | 21 |

**Supporting information 1.5** Summary of the parameter estimates

**Table S1.5.1** Average relative bias (RB), coefficient of variation (CV) and coverage of the 95% confidence interval (%) from the 50 simulated datasets for all combinations of sampling interruption scenario, life history strategy (Slow and Intermediate), and presence/absence of stochasticity in annual vital rates ((survival and recruitment).

|  | **Slow** | | | **Slow stochastic** | | | **Intermediate** | | | **Intermediate stochastic** | | |
| --- | --- | --- | --- | --- | --- | --- | --- | --- | --- | --- | --- | --- |
|  | **RB** | **CV** | **Coverage** | **RB** | **CV** | **Coverage** | **RB** | **CV** | **Coverage** | **RB** | **CV** | **Coverage** |
| **Scenario 1 (11111)** | | | | | | | | | | | | |
|  | -0.02 | 8.84 | 98 | -0.03 | 8.74 | 98 | -0.03 | 9.66 | 100 | 0.02 | 9.26 | 98 |
|  | -0.01 | 7.12 | 98 | -0.02 | 7.01 | 92 | -0.02 | 7.74 | 94 | 0 | 7.69 | 100 |
|  | -0.01 | 6.36 | 94 | -0.01 | 6.53 | 98 | 0 | 7.53 | 98 | 0 | 7.3 | 94 |
|  | -0.02 | 6.57 | 98 | -0.03 | 6.72 | 96 | 0.02 | 7.65 | 96 | -0.01 | 7.44 | 90 |
|  | 0 | 8.25 | 92 | -0.03 | 8.35 | 100 | 0.02 | 8.98 | 96 | -0.01 | 8.95 | 90 |
| **Scenario 2 (11011)** | | | | | | | | | | | | |
|  | 0 | 9.12 | 94 | -0.02 | 9.03 | 98 | -0.03 | 9.81 | 96 | -0.01 | 9.61 | 90 |
|  | 0 | 7.58 | 90 | 0 | 7.81 | 94 | -0.01 | 8.47 | 94 | 0 | 8.34 | 98 |
|  | 0 | 8.17 | 98 | 0.01 | 8.37 | 92 | 0 | 10.12 | 98 | 0.01 | 10.04 | 98 |
|  | 0 | 7.2 | 100 | -0.01 | 7.39 | 92 | -0.01 | 8.05 | 100 | 0 | 8.03 | 96 |
|  | -0.01 | 8.56 | 100 | -0.01 | 8.64 | 94 | -0.02 | 9.37 | 96 | -0.01 | 9.19 | 94 |
| **Scenario 3 (10011)** | | | | | | | | | | | | |
|  | -0.01 | 10.54 | 90 | -0.01 | 10.66 | 98 | -0.04 | 11.03 | 92 | 0.02 | 10.52 | 100 |
|  | 0.02 | 10.2 | 90 | -0.01 | 10.2 | 96 | -0.01 | 12.06 | 94 | 0.04 | 11.58 | 96 |
|  | 0.02 | 9.31 | 98 | -0.01 | 9.27 | 90 | -0.01 | 11.28 | 88 | 0.04 | 10.89 | 92 |
|  | 0 | 7.65 | 96 | -0.01 | 7.62 | 98 | -0.01 | 8.24 | 96 | -0.01 | 8.02 | 94 |
|  | 0 | 8.72 | 92 | 0 | 8.81 | 88 | 0 | 9.3 | 96 | 0 | 9.02 | 96 |
| **Scenario 4 (11001)** | | | | | | | | | | | | |
|  | -0.02 | 9.39 | 98 | 0 | 9.26 | 96 | -0.01 | 9.7 | 98 | 0.01 | 9.76 | 100 |
|  | -0.01 | 8.13 | 96 | 0 | 8.15 | 94 | -0.01 | 8.75 | 100 | 0 | 8.76 | 94 |
|  | -0.01 | 9.39 | 100 | 0.02 | 9.46 | 100 | 0.02 | 11.28 | 100 | 0.02 | 11.4 | 96 |
|  | -0.01 | 9.91 | 100 | 0 | 10 | 100 | 0.02 | 11.62 | 96 | 0.02 | 11.82 | 90 |
|  | -0.01 | 9.88 | 98 | -0.02 | 9.9 | 96 | 0.01 | 10.08 | 96 | -0.02 | 10.31 | 92 |
| **Scenario 5 (10111)** | | | | | | | | | | | | |
|  | -0.02 | 9.86 | 98 | 0 | 9.77 | 92 | -0.03 | 10.32 | 98 | -0.01 | 10.29 | 100 |
|  | -0.02 | 9.26 | 98 | -0.01 | 9.13 | 96 | -0.01 | 10.58 | 96 | 0.01 | 10.66 | 92 |
|  | -0.01 | 7.29 | 98 | -0.03 | 7.17 | 94 | 0 | 7.77 | 92 | 0.01 | 7.97 | 100 |
|  | -0.01 | 7 | 92 | -0.02 | 7.16 | 96 | 0 | 7.59 | 92 | 0 | 7.82 | 92 |
|  | 0 | 8.42 | 92 | -0.02 | 8.71 | 98 | -0.01 | 9.03 | 96 | 0 | 9.17 | 98 |
| **Scenario 6 (11101)** | | | | | | | | | | | | |
|  | 0 | 8.92 | 90 | 0 | 8.83 | 98 | -0.01 | 9.61 | 98 | 0.02 | 9.48 | 92 |
|  | 0 | 7.37 | 92 | -0.01 | 7.29 | 96 | 0.01 | 7.97 | 96 | 0.01 | 8.03 | 88 |
|  | -0.01 | 7.05 | 94 | -0.01 | 7.01 | 98 | 0 | 7.94 | 94 | -0.01 | 8.04 | 94 |
|  | -0.02 | 8.8 | 92 | 0.01 | 8.71 | 96 | 0.02 | 10.54 | 92 | -0.01 | 10.46 | 96 |
|  | -0.02 | 9.24 | 98 | -0.01 | 9.25 | 96 | -0.04 | 9.95 | 92 | -0.02 | 9.84 | 94 |
| **Scenario 7 (10101)** | | | | | | | | | | | | |
|  | -0.01 | 9.98 | 94 | -0.02 | 10.03 | 98 | 0 | 10.39 | 96 | 0 | 10.39 | 98 |
|  | 0 | 9.28 | 88 | -0.02 | 9.4 | 90 | 0 | 10.79 | 96 | 0.05 | 10.79 | 90 |
|  | 0 | 7.63 | 100 | -0.03 | 7.78 | 96 | -0.01 | 8.49 | 88 | 0.02 | 8.39 | 94 |
|  | -0.01 | 9.03 | 98 | -0.01 | 9.08 | 94 | -0.01 | 10.57 | 96 | 0.02 | 10.59 | 92 |
|  | -0.02 | 9.42 | 98 | -0.02 | 9.38 | 96 | -0.02 | 9.89 | 98 | 0 | 9.89 | 100 |
| **Scenario 8 (01111)** | | | | | | | | | | | | |
|  | 0.01 | 14.41 | 100 | -0.02 | 14.62 | 96 | -0.02 | 17.71 | 94 | -0.04 | 17.89 | 94 |
|  | 0 | 8.76 | 96 | -0.01 | 8.78 | 100 | -0.01 | 9.58 | 96 | -0.01 | 9.74 | 92 |
|  | 0 | 6.91 | 96 | -0.01 | 7.09 | 98 | 0 | 7.77 | 100 | -0.01 | 7.73 | 92 |
|  | -0.01 | 6.79 | 100 | -0.01 | 7 | 100 | 0 | 7.66 | 96 | -0.01 | 7.6 | 94 |
|  | -0.02 | 8.52 | 100 | -0.03 | 8.76 | 98 | 0.01 | 9.32 | 92 | 0 | 9.57 | 96 |
| **Scenario 9 (11110)** | | | | | | | | | | | | |
|  | 0 | 9.19 | 98 | 0 | 9 | 98 | 0.01 | 9.8 | 96 | -0.02 | 9.79 | 94 |
|  | -0.01 | 7.2 | 96 | 0 | 7.05 | 96 | 0 | 7.91 | 98 | -0.02 | 7.89 | 88 |
|  | -0.01 | 7.01 | 90 | -0.01 | 6.83 | 98 | 0 | 7.67 | 96 | 0 | 7.7 | 94 |
|  | -0.03 | 8.62 | 92 | -0.01 | 8.55 | 94 | 0.01 | 9.46 | 98 | -0.01 | 9.25 | 100 |
|  | -0.02 | 14.56 | 92 | 0.01 | 14.3 | 88 | 0 | 17.72 | 100 | -0.01 | 17.43 | 96 |
| **Scenario 10 (10001)** | | | | | | | | | | | | |
|  | 0 | 10.87 | 96 | -0.01 | 10.98 | 98 | 0 | 11.3 | 100 | -0.03 | 11.16 | 100 |
|  | 0 | 10.89 | 98 | 0.01 | 11.18 | 96 | 0.02 | 12.52 | 93.3 | 0 | 12.68 | 94.4 |
|  | 0.02 | 10.81 | 94 | 0.01 | 11.11 | 98 | 0 | 12.76 | 90 | -0.04 | 13.06 | 94.4 |
|  | 0.02 | 10.65 | 94 | 0.04 | 10.88 | 90 | -0.01 | 12.16 | 86.7 | -0.05 | 12.47 | 88.9 |
|  | 0.01 | 10.41 | 94 | 0.02 | 10.42 | 96 | 0.01 | 10.7 | 86.7 | -0.01 | 10.83 | 100 |

**Supporting information 1.6**

**P**arameter identifiability in the presence of interruption**.**

**Methods**

Occasions with interruptions in sampling yield no individuals detections. Inferences are nonetheless possible, primarily due to the Markovian structure of the model and the reconstructed *z* matrix (Figure 1), as well as due to the assumption of parameter exchangeability among years (i.e. constant survival/recruitment rates assumed in our study). Because the reconstructed *z* should contain some information about individual survival, we used simulations to test whether annual estimates of survival ($\phi_{t}$) are identifiable in the presence of interruptions. For simplicity, we only chose scenario 11011. We simulated detection datasets as described in 2.2 in the main text, and fitted the OPSCR model (as described in sections 2.1 and 2.5), with the exception $\phi_{t}$ being estimated independently for each transition. Because the interruption occurred at the 3^rd^ occasion, we expect $\phi_{2}$(survival between occasion 2 and 3) and $\phi_{3}$ (survival between occasion 3 and 4) to be weakly identifiable. Weak identifiability arises when data provides little information about parameters (Gimenez et al. 2009). In the case of interruption and low survival rates of individuals, no detections available and few information in the reconstructed *z*, means that a low $\phi_{2}$ and high $\phi_{3}$, or a high $\phi_{2}$ and a low $\phi_{3}$ could be equally likely, and therefore pose convergence and/or parameter identifiability issues (Gimenez et al. 2009). We simulated 25 datasets for species with a slow and an intermediate life history (see table 1 of the main text for a definition), and an initial population size of 50 and 100 individuals. We expected estimation of $\phi_{t}$ to be more challenging for species with a fast life history, as it is more difficult to reconstruct *z* during the interruption and for populations with a smaller population size, as it leads to smaller sample size.

We applied two simple measures to identify weakly identifiable parameters as described by Gimenez et al. (2009). We computed a measure of prior-posterior distribution overlap. A ≥35% overlap has been suggested as an indicator of weak identifiability; and 2) examined the correlation matrix of the posterior distribution of $\phi_{t}$. We then reported the average and standard deviation of prior-posterior distribution overlap and of Pearson’s correlation between the different parameters across the 25 data set.

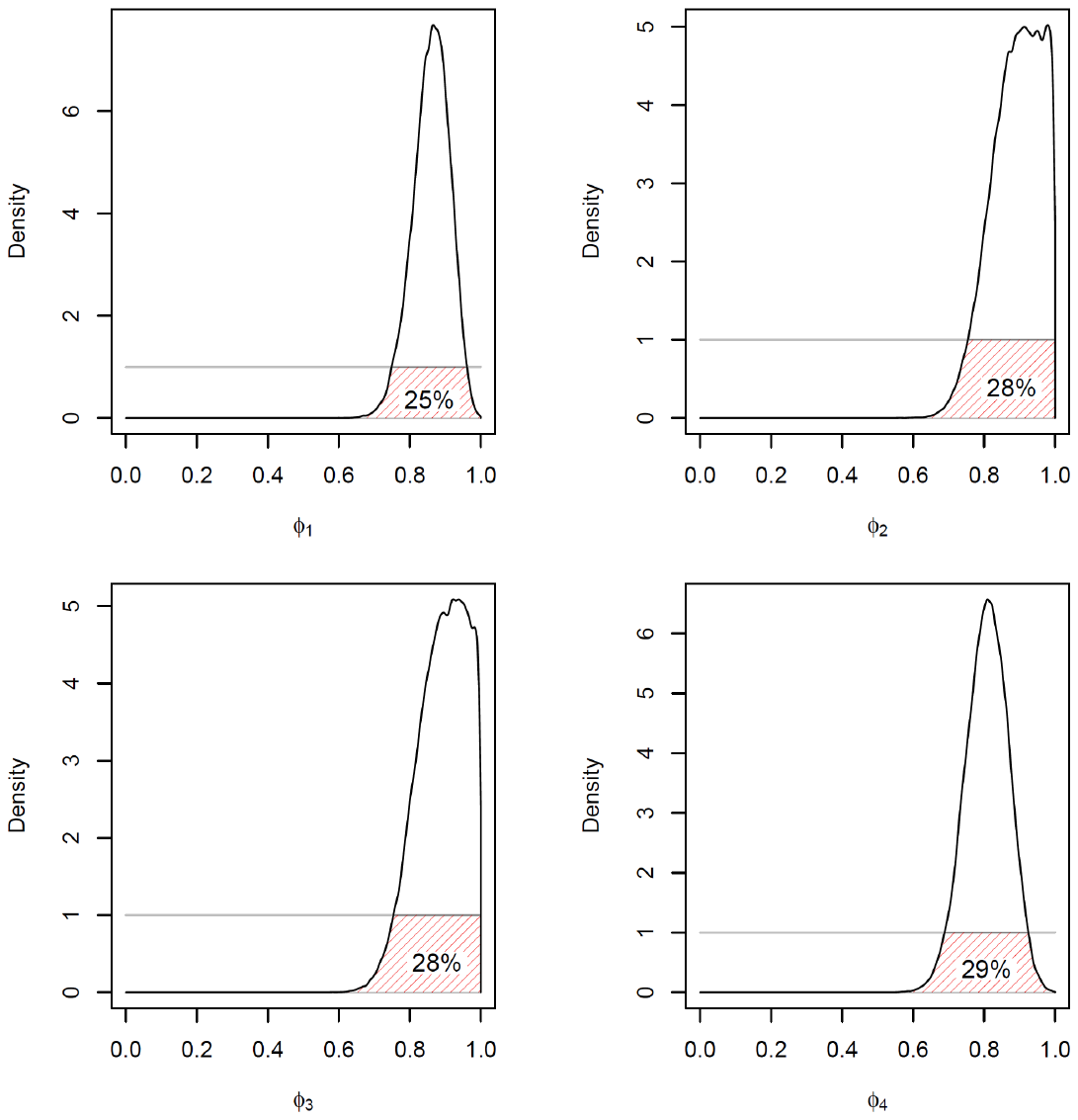

**Figure S1.6.1**. Comparison between prior (grey flat line)-posterior (black line) distributions for the survival parameter ($\phi_{t}$) from an open population spatial capture recapture model fitted to simulated data with sampling interruption (11011; **See Figure 2 main text**) for a species with a slow speed of life history and an initial population size =100. Values represent the percentage of overlap between prior and posterior distributions. The figure illustrates the prior/posterior overlap results for one simulated data set.

**Results and discussion**

As expected, we found parameter identifiability of $\phi_{t}$ to be higher for species with slower-paced life histories and for larger populations. While the degree of prior-posterior overlap was generally ≤35% (for all $\phi_{t}$) for species with a slow life history , it was larger for intermediate species, especially for $\phi_{2}$ and $\phi_{3}$( <50% Table S1.6.1). However, there was a negative correlation (p < -0.6) between $\phi_{2}$ and $\phi_{3}$for both life history strategies and both initial population sizes (Table S1.6.2). Together, these results suggests that the reconstructed *z* contains some information about individual survival, especially for species with a slow life history and large sample size. However, the correlation between $\phi_{2}$ and $\phi_{3}$ suggest poor identifiability and the need to assume and use constant parameters across years, to use covariates, or use other sources of information (e.g. dead recovery) to provide ancillary information for gaps in the monitoring data.

**Table S1.6.1** Average (sd) of prior-posterior overlap distributions between survival estimates ($\phi_{t}$) using an open population spatial capture recapture model on 25 simulated data sets with sampling interruption (11011; **See Figure 2 main text**) for species with a slow and intermediate life history, and for populations with an initial population size (N_1_) of 50 and 100 individuals.

.

|  | $\boldsymbol{\phi}_{\boldsymbol{1}}$ | $\boldsymbol{\phi}_{\boldsymbol{2}}$ | $\boldsymbol{\phi}_{\boldsymbol{3}}$ | $\boldsymbol{\phi}_{\boldsymbol{4}}$ |
| --- | --- | --- | --- | --- |
| N_1_ | **Slow** | | | |
| 50 | 34.7 (0.73) | 31.2 (0.89) | 32.4 (4.07) | 36.1 (1.11) |
| 100 | 24.9 (0.30) | 28.4 (0.21) | 28.4 (0.22) | 28.4 (0.28) |
|  | **Intermediate** | | | |
| 50 | 42.2 (0.22) | 59.2 (1.76) | 53.8 (5.79) | 40.0 (0.17) |
| 100 | 30.3 (0.67) | 62.4 (4.24) | 58.9 (9.14) | 30.9 (0.16) |

**Table S1.6.2** Average (sd) Pearson *r* correlation coefficients between survival estimates ($\phi_{t}$) and per capita recruitment ($\boldsymbol{\rho)}$ using an open population spatial capture recapture model. Results are presented for 25 simulated data sets with sampling interruption (11011; **See Figure 2 main text**) for species with slow and intermediate life history speeds, and for initial population sizes (N_1_) of 50 and 100 individuals.

|  | $\boldsymbol{\phi}_{\boldsymbol{1}}$ | $\boldsymbol{\phi}_{\boldsymbol{2}}$ | $\boldsymbol{\phi}_{\boldsymbol{3}}$ | $\boldsymbol{\phi}_{\boldsymbol{4}}$ | $\boldsymbol{\rho}$ |
| --- | --- | --- | --- | --- | --- |
| Slow | | | | | |
| *N_1_=50* | | | | | |
| $\boldsymbol{\phi}_{\boldsymbol{1}}$ | 1 | -0.08 (0.007) | -0.01 (0.057) | 0.00 (0.007) | -0.13 (0.000) |
| $\boldsymbol{\phi}_{\boldsymbol{2}}$ |  | 1 | -0.54 (0.191) | 0.01 (0.064) | 0.26 (0.410) |
| $\boldsymbol{\phi}_{\boldsymbol{3}}$ |  |  | 1 | -0.08 (0.064) | -0.46 (0.410) |
| $\boldsymbol{\phi}_{\boldsymbol{4}}$ |  |  |  | 1 | 0.02 (0.078) |
| $\boldsymbol{\rho}$ |  |  |  |  | 1 |
| *N_1_=100* | | | | | |
| $\boldsymbol{\phi}_{\boldsymbol{1}}$ | 1 | -0.06 (0.007) | -0.05 (0.014) | 0.00 (0.007) | -0.13 (0.014) |
| $\boldsymbol{\phi}_{\boldsymbol{2}}$ |  | 1 | -0.60 (0.021) | -0.04 (0.007) | -0.06 (0.014) |
| $\boldsymbol{\phi}_{\boldsymbol{3}}$ |  |  | 1 | -0.04 (0.007) | -0.06 (0.007) |
| $\boldsymbol{\phi}_{\boldsymbol{4}}$ |  |  |  | 1 | -0.04 (0.000) |
| $\boldsymbol{\rho}$ |  |  |  |  | 1 |
| Intermediate | | | | | |
| *N_1_=50* | | | | | |
| $\boldsymbol{\phi}_{\boldsymbol{1}}$ | 1 | -0.06 (0.028) | 0.00 (0.078) | 0.00 (0.007) | -0.16 (0.049) |
| $\boldsymbol{\phi}_{\boldsymbol{2}}$ |  | 1 | -0.60 (0.064) | -0.02 (0.021) | -0.07 (0.198) |
| $\boldsymbol{\phi}_{\boldsymbol{3}}$ |  |  | 1 | -0.02 (0.099) | -0.25 (0.297) |
| $\boldsymbol{\phi}_{\boldsymbol{4}}$ |  |  |  | 1 | -0.15 (0.035) |
| $\boldsymbol{\rho}$ |  |  |  |  | 1 |
| *N_1_=100* | | | | | |
| $\boldsymbol{\phi}_{\boldsymbol{1}}$ | 1 | -0.04 (0.064) | 0.05 (0.113) | 0.01 (0.014) | -0.22 (0.106) |
| $\boldsymbol{\phi}_{\boldsymbol{2}}$ |  | 1 | -0.80 (0.078) | -0.02 (0.028) | 0.00 (0.163) |
| $\boldsymbol{\phi}_{\boldsymbol{3}}$ |  |  | 1 | 0.00 (0.035) | -0.26 (0.304) |
| $\boldsymbol{\phi}_{\boldsymbol{4}}$ |  |  |  | 1 | -0.08 (0.078) |
| $\boldsymbol{\rho}$ |  |  |  |  | 1 |

**References**

Gardner, B., R. Sollmann, N. S. Kumar, D. Jathanna, and K. U. Karanth. 2018. State space and movement specification in open population spatial capture-recapture models. Ecology and Evolution 0.

Gimenez, O., B. J. T. Morgan, and S. P. Brooks. 2009. Weak identifiability in models for mark-recapture-recovery data. Pages 1055–1067 Modeling demographic processes in marked populations. Springer.
