## Supporting informations for "Estimating abundance with interruptions in data collection using open population spatial capture-recapture models": SupportingInformation2.pdf

This script performs a data set simulation and OPSCR modelling in the presence of sampling interuptions with NIMBLE (NIMBLE Development Team 2019; Valpine et al. 2017). All details about the procedure are provided in Milleret et al. Estimating abundance with interruption in data collection: Simulations and case studies using open population spatial capture-recapture models.

#### I.LOAD LIBRARIES AND SET WORKING DIRECTORY

```
rm(list = ls())
library(rgdal)
library(raster)
library(rgeos)
library(sp)
library(nimble)
library(abind)
library(boot)
library(coda)
```

Set working directory where the *SourceFunctions.R*, *SourceNimblePointProcess.R* and *SourceNimbleObservationModel.R* are located and source the files.

```
setwd("YourWorkingdirectory")
source("SourceRFunctions.R")
source("SourceNimblePointProcess.R")
source("SourceNimbleObservationModel.R")
```

```
## Registering the following user-provided distributions: dbinomPP dbinomPPSingle dbinomMNormSourcePP d

## Registering the following user-provided distributions: dbin_LESS .
```

#### II.SET SIMULATION PARAMETERS

```
# HABITAT EXTENT
buffer <- 5
grid.size <- 30
# DETECTOR SPACING
detector.spacing <- 1.5
# DETECTION FUNCTION SURVEY CHARACTERISTICS
p0 <- 0.25  # p0 FOR THE HALFNORMAL DETECTION FUNCTION
sigma <- 2  # SIGMA FOR THE HALFNORMAL DETECTION FUNCTION
n.occasions <- 5  # NB OCCASIONS

# POPULATION CHARACTERISTICS
N1 <- 50  # N INDIVIDUALS AT FIRST OCCASION
phi <- 0.85  # SURVIVAL
rho <- 0.15  # PER CAPITA RECRUITMENT
```

```
sd.phi <- 0  # SD OF THE SURVIVAL (IF STOCHASTICITY)
sd.rho <- 0  # SD OF THE PER CAPITA (IF STOCHASTICITY)
p.repro <- 1  # PROBABILITY OF INDIVIDUALS REPRODUCING (if = 1, ASSUME ALL INDIVIDUALS REPRODUCE)
dispSigma <- 3  # TAU(DISPERSAL SIGMA)

# SAMPLING INTERUPTION
toggle.interuption <- c(1, 1, 1, 1, 1)  # WHETHER INTERUPTION OCCUR (1) OR NOT (0) AT EACH OCCASION
# LEVEL OF AUGMENTATION
augmentation <- 1.2  # DATASET IS AUGMENTED BY N AUGMENTED INDIVIDUALS THAT EQUALS TO:
# SUPERPOPULATION SIZE * augmentation

# NIMBLE RUN CHARACTERISTICS
nburnin <- 1000  # BURN-IN
niter <- 10000  # N ITERATIONS
nchains <- 3  # N CHAINS
```

#### III.CREATE SIMULATED DATASET

```
### ==== 1. CREATE A SQUARE SPATIAL DOMAIN ====
coords <- matrix(c(0, 0, grid.size + buffer * 2, 0, grid.size +
    buffer * 2, grid.size + buffer * 2, 0, grid.size + buffer *
    2, 0, 0), ncol = 2, byrow = TRUE)

P1 <- Polygon(coords)
myStudyArea <- SpatialPolygons(list(Polygons(list(P1), ID = "a")),
    proj4string = CRS("+proj=utm +zone=33 +ellps=WGS84 +datum=WGS84 +units=m +no_defs"))
### ==== 1.2. HABITAT RASTER ====
r <- raster(nrow = 2, ncol = 2, xmn = 0, xmx = grid.size + buffer *
    2, ymn = 0, ymx = grid.size + buffer * 2)
# HABITAT QUALITY VALUES, WE ASSUME HOMOGENEOUS HABITAT
# QUALITY
habitatQuality <- r[] <- 1
proj4string(r) <- CRS(proj4string(myStudyArea))
resolution <- res(r)
lowerCoords <- coordinates(r) - resolution/2
upperCoords <- coordinates(r) + resolution/2
habitatQuality <- r[]

### ==== 2.GENERATE DETECTORS ====
co <- seq(buffer, grid.size + buffer, by = detector.spacing)
x <- rep(co, length(co))
y <- sort(rep(co, length(co)), decreasing = T)
detectors.xy <- cbind(x, y)
detectors.sp <- SpatialPoints(detectors.xy, proj4string = CRS(proj4string(myStudyArea)))
# PLOT CHECK
plot(r)
plot(myStudyArea, add = T)
points(detectors.sp, col = "black", pch = 16)
```

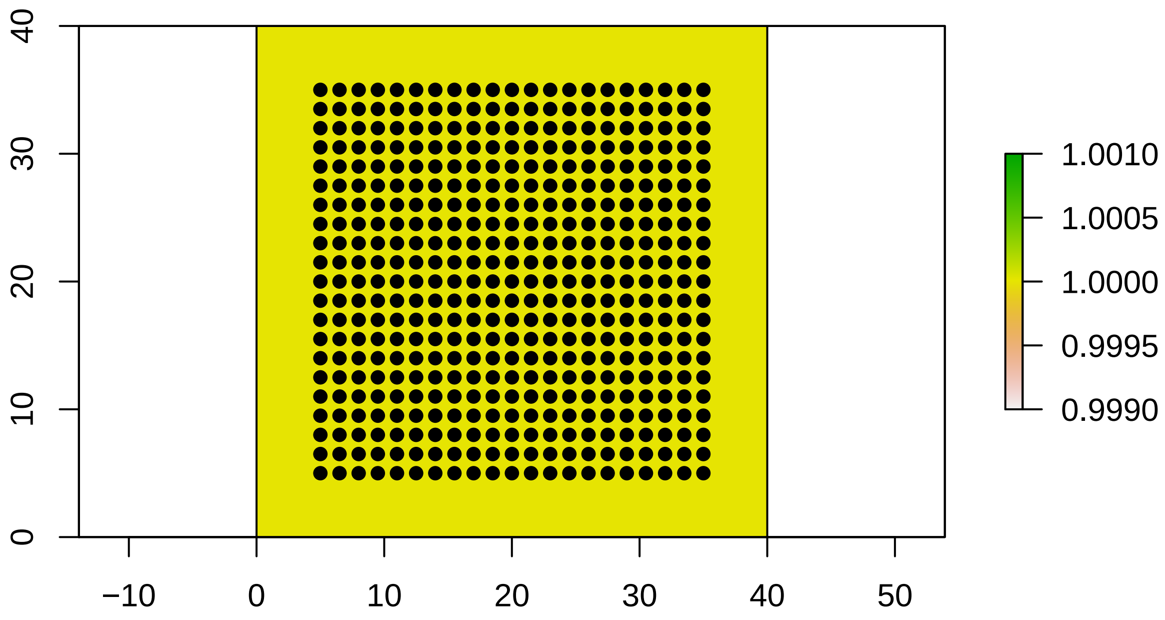


```r
### ==== 3.SIMULATE INDIVIDUAL STATE MATRIX : z.mx ==== ====
### 3.1 DEFINE PHI AND RHO ARRAYS ==== CREATE ARRAYS SO PHI AND
### FEC CAN VARRY OVER TIME AND STOCHASTICITY IN VITAL RATES
### CAN BE ADDED (t)
PHI.arr <- array(NA, c(2, 2, n.occasions))
RHO.arr <- array(NA, c(2, 2, n.occasions))
FEC.mat <- matrix(NA, nrow = 2, ncol = n.occasions)
phit <- 0
fect <- 0

for (t in 1:n.occasions) {
    # DEFINE THE PHI MATRIX two states: ALIVE/DEAD DRAW PHI FROM
    # NORMAL DISTRIB AND APPLY LOGIT
    phit[t] <- inv.logit(rnorm(1, mean = logit(phi), sd = sd.phi))
    PHI.arr[, , t] <- matrix(c(phit[t], 0, 1 - phit[t], 1), ncol = 2,
        nrow = 2, byrow = TRUE)
    # DEFINE REPRO MATRIX (p of reproducing) THIS DEFINES THE
    # PROBABILITY OF INDIVIDUALS TO REPRODUCE (ASSUME ALL
    # INDIVIDUALS REPRODUCE HERE)
    RHO.arr[, , t] = matrix(c(p.repro, 0, 0, 0), ncol = 2, nrow = 2,
        byrow = TRUE)
    # DEFINE PER CAPITA RECRUITMENT. GIVEN THAT INDIVIDUAL
    # REPRODUCES, HOW MANY ARE RECRUITED
    fect[t] <- inv.logit(rnorm(1, mean = logit(rho), sd = sd.rho))
    FEC.mat[, t] = c(fect[t], 0)
}
```

```r
### ==== 3.2 SIMULATE Z ====
z.mx <- SimulateZ(N0 = N1, n.occasions = n.occasions - 1, PHI = PHI.arr,
    REPRO = RHO.arr, FEC = FEC.mat, init.state = 1)
myZ <- z.mx$z
# ADD THE NOT ENTERED STATE 1: NOT ENTERERD 2: ALIVE 3: DEAD
myZ <- myZ + 1
myZ[is.na(myZ)] <- 1

# GET POP COMPOSITION AND PLOT IT
z.levels <- unique(na.omit(unlist(apply(myZ, 1, unique))))
z.levels <- z.levels[order(z.levels)]
Pop.Compo <- list()
for (l in 1:length(z.levels)) {
    Pop.Compo[[l]] <- apply(myZ, 2, function(x) {
        length(which(x == z.levels[l]))
    })
}  #l

# PLOT CHECK
plot(1:dim(myZ)[2], Pop.Compo[[1]], type = "l", col = 1, lwd = 2,
    ylim = c(0, max(unlist(Pop.Compo))), xlab = "Years", ylab = "# of Individuals")

for (l in 2:length(Pop.Compo)) {
    points(1:dim(myZ)[2], Pop.Compo[[l]], type = "l", col = l,
        lwd = 2)
}  #l
legend("bottomleft", legend = c(unlist(lapply(z.levels, function(x) {
    paste(" State", x)
}))), col = 1:3, lwd = 1, cex = 0.6, inset = 0.01)
```

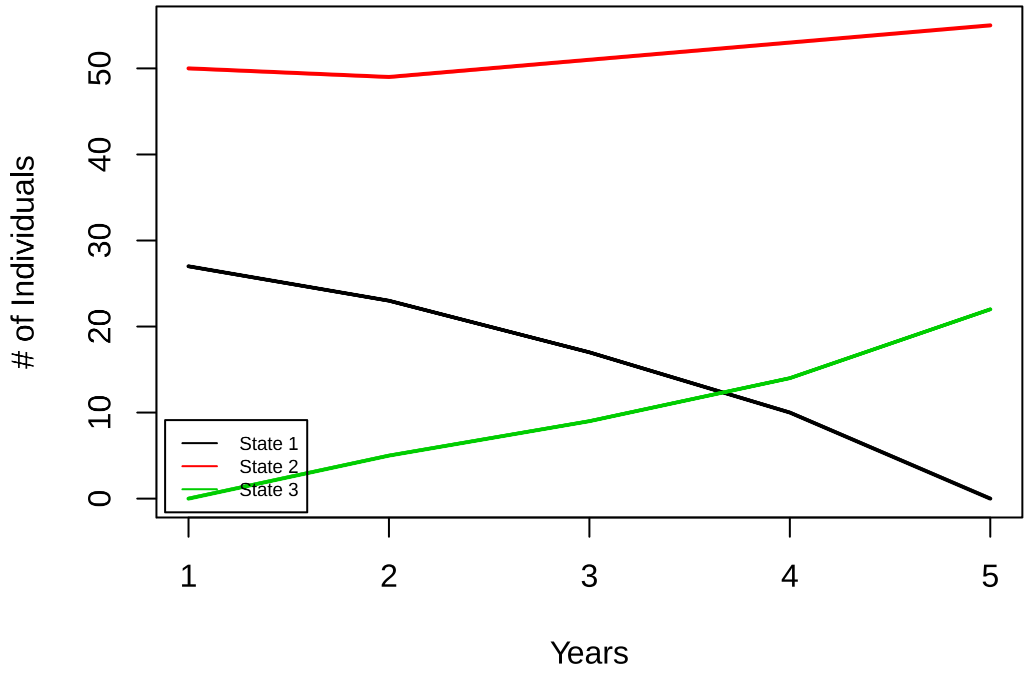


```
### ==== 4.SIMULATE INDIVIDUAL AC LOCATIONS ==== ==== 4.1 FIRST
### OCCASION ==== CREATE EMPTY OBJECTS
mySimulatedACs <- list()
tempCoords <- matrix(NA, nrow = dim(myZ)[1], ncol = 2)

# SIMULATE UNIFORM LOCATION OF ACS
for (i in 1:dim(myZ)[1]) {
    tempCoords[i, ] <- rbinomPPSingle(n = 1, lowerCoords = lowerCoords,
        upperCoords = upperCoords, intensityWeights = habitatQuality,
        areAreas = 1, numWindows = nrow(lowerCoords))
}  #i

# STORE ACS IN A SP OBJECT
mySimulatedACs[[1]] <- SpatialPointsDataFrame(tempCoords, data.frame(x = tempCoords[,
    1], y = tempCoords[, 2], Id = 1:nrow(tempCoords)), proj4string = CRS(proj4string(r)))
### ==== 4.2 FOLLWOWING YEARS ==== DRAW SUBSEQUENT INDIVIDUAL
### ACS FROM A NORMAL DISTRIBUTION CENTERED AROUND THE SOURCE
### COORDINATE WITH A SD ('normSD') EQUAL TO THE DISPERSAL
### SIGMA ('dispSigma') THERE IS THE POSSIBILITY TO DEFINE A
### HABITAT QUALITY SURFACE FOR THE PUPORSE OF THE STUDY,
### HABITAT QUALITY WAS UNIFORM (SET TO 1 EVERYWHERE)
for (t in 2:dim(myZ)[2]) {
    for (i in 1:nrow(tempCoords)) {
        tempCoords[i, ] <- rbinomMNormSourcePPSingle(n = 1, lowerCoords = lowerCoords,
            upperCoords = upperCoords, sourceCoords = tempCoords[i,
                ], normSD = dispSigma, intensityWeights = habitatQuality,
```

```
            areAreas = 1, numWindows = nrow(lowerCoords))
    }
    # STORE ACS IN A SP OBJECT
    mySimulatedACs[[t]] <- SpatialPointsDataFrame(tempCoords,
        data.frame(x = tempCoords[, 1], y = tempCoords[, 2],
            Id = 1:nrow(tempCoords)), proj4string = CRS(proj4string(mySimulatedACs[[t -
            1]])))
}  #t

# PLOT CHECK
plot(aggregate(rasterToPolygons(r, fun = function(x) {
    x > 0
})))
points(detectors.sp, pch = 16, cex = 0.5)
col <- rainbow(length(mySimulatedACs[[1]]))
points(mySimulatedACs[[1]], pch = 21, bg = col)

for (t in 2:length(mySimulatedACs)) {
    points(mySimulatedACs[[t]], pch = 21, bg = col)
    arrows(x0 = coordinates(mySimulatedACs[[t]])[, 1], x1 = coordinates(mySimulatedACs[[t -
        1]])[, 1], y0 = coordinates(mySimulatedACs[[t]])[, 2],
        y1 = coordinates(mySimulatedACs[[t - 1]])[, 2], col = col,
        length = 0.08)
}  #t
```

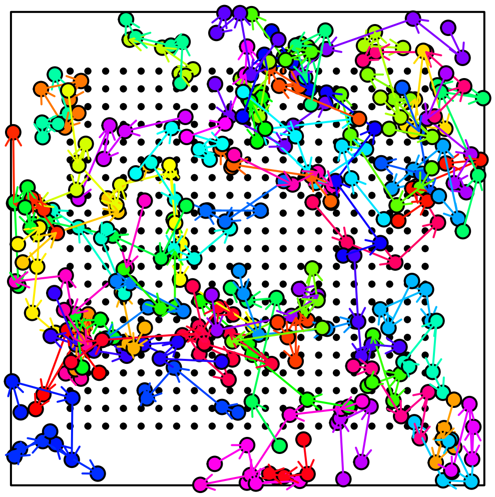

```
### ==== 5.SIMULATE DETECTION ==== ==== 5.1 DETECTION ARRAY
### ====
z.not.alive <- apply(myZ, 2, function(x) {
    which(x %in% c(1, 3))
})
Y <- array(NA, c(dim(myZ)[1], length(detectors.sp), dim(myZ)[2]))

for (t in 1:dim(myZ)[2]) {
    D <- gDistance(detectors.sp, mySimulatedACs[[t]], byid = TRUE)
    # OBTAIN Y DETECTION MATRIX USING HALF NORMAL DETECTION
    # FUNCTION (EQN 7 IN MAIN TEXT)
    fixed.effects <- rep(log(p0), length(detectors.sp))
    pzero <- exp(fixed.effects)
    p <- pzero * exp(-D * D/(2 * sigma * sigma))
    Y[, , t] <- apply(p, c(1, 2), function(x) rbinom(1, 1, x))
    # individuals not alive can't be detected
    Y[z.not.alive[[t]], , t] <- 0

}  #t
### ==== 5.1 PLOT CHECK ====
par(mfrow = c(2, 3), mar = c(0, 0, 3, 0))
for (t in 1:dim(myZ)[2]) {
    plot(myStudyArea)
    title(t)
    points(detectors.sp, pch = 16, cex = 0.6)
    detections <- apply(Y[, , t], 1, function(x) which(x > 0))
    col <- rainbow(dim(Y)[1])[sample(dim(Y)[1])]
    for (i in 1:length(detections)) {
        if (!i %in% z.not.alive[[t]])
            {
                if (length(detectors.sp[detections[[i]], ]) ==
                  0) {
                  points(mySimulatedACs[[t]][i, ], bg = col[i],
                    pch = 21, cex = 0.5)
                  points(mySimulatedACs[[t]][i, ], col = col[i],
                    pch = 4)

                } else {
                  points(detectors.sp[detections[[i]], ], col = col[i],
                    pch = 16, cex = 0.7)
                  ac <- coordinates(mySimulatedACs[[t]][i, ])
                  dets <- coordinates(detectors.sp[detections[[i]],
                    ])
                  segments(x0 = ac[1, 1], x1 = dets[, 1], y0 = ac[1,
                    2], y1 = dets[, 2], col = col[i])
                  points(mySimulatedACs[[t]][i, ], bg = col[i],
                    pch = 21)

                }  #else
            }  #if
    }  #i
}  #t
```

```r
### ==== 6.AUGMENT DATA SET ==== REMOVE UNDETECTED IDS
detected.time <- apply(Y, c(1, 3), function(x) any(x >= 1))
detected <- apply(detected.time, c(1), function(x) sum(x == 1) >
    0)
Y <- Y[detected, , ]

# AUGMENTATION= 'augmentation' x N OF THE SUPER POPULATION
y.aug <- array(0, c(nrow(myZ) * augmentation - dim(Y)[1], dim(Y)[2:3]))
y <- abind(Y, y.aug, along = 1)
dim(y)
```

```
## [1]  92 441   5
```

```r
### ==== 7.ADD INTERUPTION ====
interuptions <- which(toggle.interuption == 0)
if (length(interuptions) > 0) {
    # ZERO DETECTIONS DURING INTERUPTIONS
    y[, , interuptions] <- 0
}

### ==== 8.RECONSTRUCT z VALUES ====
z <- apply(y, c(1, 3), function(x) any(x > 0))
z <- ifelse(z, 2, NA)

z <- t(apply(z, 1, function(zz) {
    if (any(!is.na(zz))) {
        range.det <- range(which(!is.na(zz)))
        zz[range.det[1]:range.det[2]] <- 2
    }
    return(zz)
}))

### ==== 9.GENERATE z INITIAL values====
z.init <- t(apply(z, 1, function(zz) {
    out <- zz
    out[] <- 1
    if (any(!is.na(zz))) {
        range.det <- range(which(!is.na(zz)))
        if (range.det[1] > 1)
            zz[1:(range.det[1] - 1)] <- 1
        if (range.det[2] < length(zz))
            zz[(range.det[2] + 1):length(zz)] <- 3
        out[] <- zz
    }
    return(out)
}))

z.init <- ifelse(!is.na(z), NA, z.init)


## -------------------------
```

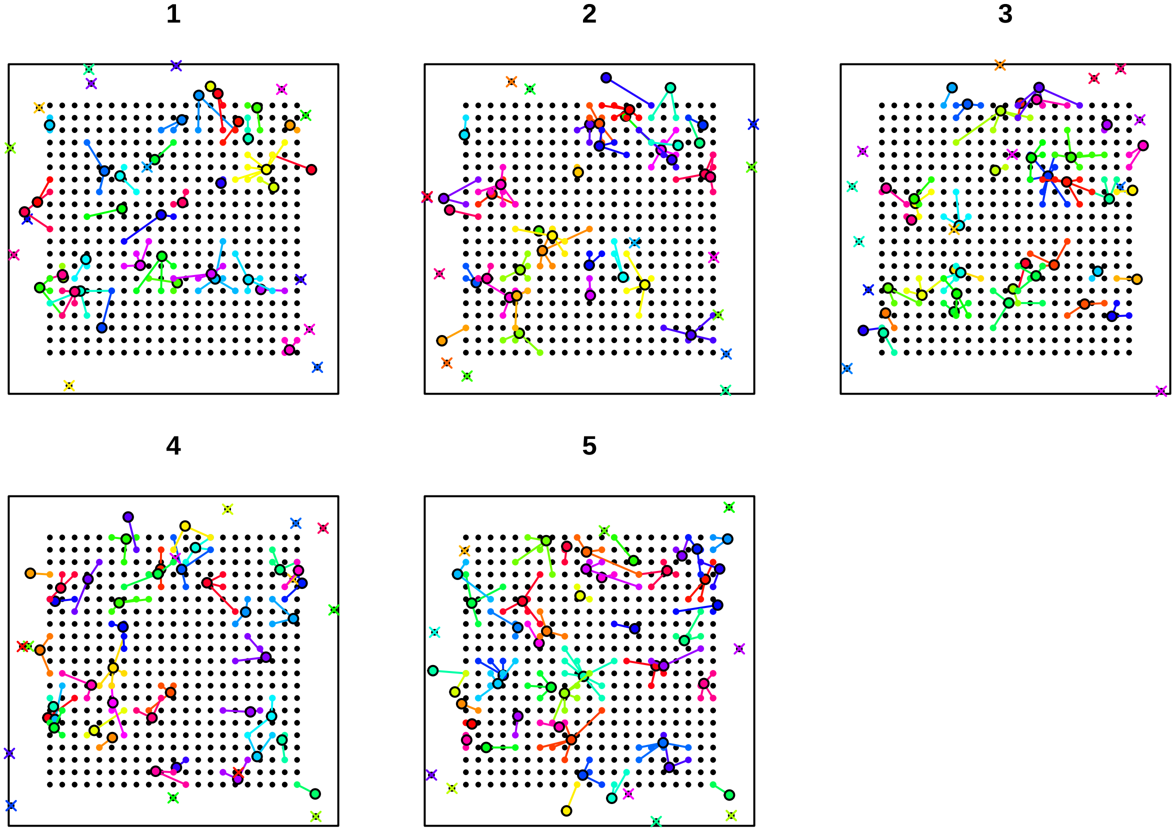


### IV.NIMBLE MODELS AND MCMC RUN

```
### ==== 1.NIMBLE MODEL ====
modelCode <- nimbleCode({
    ##-----------------------------##
    ##------ SPATIAL PROCESS ------##
    ##-----------------------------##
    dispSigma ~ dgamma(0.001, 0.001)

    for (i in 1:n.individuals) {
        sxy[i, 1:2, 1] ~ dbinomPPSingle(lowerHabCoords[1:n.cells,
            1:2], upperHabCoords[1:n.cells, 1:2], mu[1:n.cells],
            1, n.cells)
        for (t in 2:n.years) {
            sxy[i, 1:2, t] ~ dbinomMNormSourcePPSingle(lowerHabCoords[1:n.cells,
                1:2], upperHabCoords[1:n.cells, 1:2], sxy[i,
                1:2, t - 1], dispSigma, mu[1:n.cells], 1, n.cells,
                -1)
        }  #t
    }  #i

    ##-------------------------------##
    ##----- DEMOGRAPHIC PROCESS -----##
    ##-------------------------------##
```

```
psi ~ dunif(0, 1)
phi ~ dunif(0, 1)
rho ~ dunif(0, 5)

# calculate gamma from rho as a function of n.available.
for (t in 2:n.years) {
    gamma[t - 1] <- (N[t - 1] * rho)/n.available[t - 1]
}  #t

# initial gamma
omeg1[1] <- 1 - psi
omeg1[2] <- psi
omeg1[3] <- 0

for (t in 1:(nyears1)) {
    # NOT ENTERED
    omega[1, 1, t] <- 1 - gamma[t]
    omega[1, 2, t] <- gamma[t]
    omega[1, 3, t] <- 0

    # ALIVE
    omega[2, 1, t] <- 0
    omega[2, 2, t] <- phi
    omega[2, 3, t] <- 1 - phi

    # DEAD
    omega[3, 1, t] <- 0
    omega[3, 2, t] <- 0
    omega[3, 3, t] <- 1
}  #t

for (i in 1:n.individuals) {
    z[i, 1] ~ dcat(omeg1[1:3])
    for (t in 1:(nyears1)) {
        z[i, t + 1] ~ dcat(omega[z[i, t], 1:3, t])
    }  #t \t\t\t\t\t\t\t\t\t\t
}  #i

##-----------------------------##
##----- DETECTION PROCESS -----##
##-----------------------------##
sigma ~ dunif(0, 5)
p0 ~ dunif(0, 1)
for (i in 1:n.individuals) {
    for (t in 1:n.years) {
        d2[i, 1:n.detectors, t] <- calculateDistance(sxy[i,
            1:2, t], detector.xy[1:n.detectors, 1:2], z[i,
            t] == 2 * toggle[t])
        y[i, 1:n.detectors, t] ~ dbin_LESS(p0, sigma, trials[1:n.detectors],
            d2[i, 1:n.detectors, t], maxDist, z[i, t] ==
              2 * toggle[t])
    }  #t
}  #i
```

```
    ##----------------------------------------##
    ##---------- DERIVED PARAMETERS ----------##
    ##----------------------------------------##
    for (t in 1:n.years) {
        N[t] <- sum(z[1:n.individuals, t] == 2)
        n.available[t] <- sum(z[1:n.individuals, t] == 1)
    }  #t
})
```

The Nimble model contains two custom NIMBLE functions (NIMBLE Development Team 2019; Valpine et al. 2017) to improve the computing efficiency of the observation model.

- *calculateDistance* calculates the square distance between the location of an individual's activity center (ACi) and all detectors. The third argument of the function takes a numeric value (0/1). A value of 0 turns off the calculation of the distances when individual $i$ is not considered *alive*.
- *dbin_LESS* is a custom Binomial distribution function that calculates the sum of the likelihood of individual detections at all detectors, given an AC location, and p0 and $\sigma$ values. Individual detections at each detector follow a Binomial distribution with a $K$ number of trials (see full details in Milleret et al. (2018)). The fifth argument of the function specifies whether a local evaluation of the individual state space (LESS) should be performed or not (Milleret et al. (2019)). If maxDist>0, then the calculation of the likelihood is limited to the detectors that are located at a distance $<= maxDist$ from the the ACi location. The last argument of the function takes a numeric value (0/1). A value of 0 turns off the likelihood calculation when the indivudal i is not considered *alive (z[i]!=2)*. In addition, the calculations are also turned off when the year contains an interruption ($toggle[t]==0$) to speed up calculations.

```
### ==== 2.NIMBLE DATA ==== ==== 2.1 CONSTANTS ====
nimConstants <- list(n.individuals = dim(y)[1], n.detectors = dim(y)[2],
    n.years = dim(y)[3], nyears1 = dim(y)[3] - 1, n.cells = nrow(lowerCoords),
    maxDist = 12  ## THIS IS THE LOCAL EVALUATION
)
### ==== 2.2 DATA ====
nimData <- list(z = z, y = y, trials = rep(1, dim(y)[2]), toggle = toggle.interuption,
    detector.xy = as.matrix(detectors.xy), lowerHabCoords = as.matrix(lowerCoords),
    upperHabCoords = as.matrix(upperCoords), mu = habitatQuality)

### ==== 2.3 INITS ====
sxy <- MakeInitsXY(y = y, detector.xy = detectors.xy, habitat.r = r,
    dist.move = 3)

nimInits <- list(z = z.init, rho = rho, phi = phi, sigma = sigma,
    psi = 0.1, p0 = p0, sxy = sxy, dispSigma = dispSigma)

### ==== 2.5 PARAMETERS TO MONITOR ====
params <- c("N", "dispSigma", "p0", "phi", "sigma", "rho")
```

```
### ==== 3.NIMBLE RUN ====
model <- nimbleModel(code = modelCode, constants = nimConstants,
    data = nimData, inits = nimInits, check = FALSE, calculate = FALSE)
```

```
## defining model...
## building model...
## setting data and initial values...
```

```
## checking model sizes and dimensions...

## Warning in model$checkBasics(): Possible size/dimension mismatch
## amongst vectors and matrices in BUGS expression: sxy[i, 1:2, 1] ~
## dbinomPPSingle(lowerCoords = lowerHabCoords[1:4, 1:2], upperCoords =
## upperHabCoords[1:4, 1:2], intensityWeights = mu[1:4], areAreas = 1,
## numWindows = 4, lower_ = -Inf, upper_ = Inf). Ignore this warning if the
## user-provided distribution has multivariate parameters with distinct sizes
## or if size of variable differs from sizes of parameters.

## Warning in model$checkBasics(): Possible size/dimension mismatch
## amongst vectors and matrices in BUGS expression: sxy[i, 1:2, t] ~
## dbinomMNormSourcePPSingle(lowerCoords = lowerHabCoords[1:4, 1:2],
## upperCoords = upperHabCoords[1:4, 1:2], sourceCoords = sxy[i, 1:2, t - 1],
## normSD = dispSigma, intensityWeights = mu[1:4], areAreas = 1, numWindows =
## 4, localEvalParam = -1, lower_ = -Inf, upper_ = Inf). Ignore this warning
## if the user-provided distribution has multivariate parameters with distinct
## sizes or if size of variable differs from sizes of parameters.

##  This model is not fully initialized. This is not an error. To see which variables are not initializ
## model building finished.
```

```
cmodel <- compileNimble(model)
```

```
## compiling... this may take a minute. Use 'showCompilerOutput = TRUE' to see C++ compilation details.
## compilation finished.
```

```
cmodel$calculate()  #okay
```

```
## [1] -3998.526
```

```
MCMCconf <- configureMCMC(model = model, monitors = c(params),
    control = list(reflective = TRUE, adaptScaleOnly = TRUE),
    useConjugacy = FALSE)

MCMC <- buildMCMC(MCMCconf)
cMCMC <- compileNimble(MCMC, project = model, resetFunctions = TRUE)
```

```
## compiling... this may take a minute. Use 'showCompilerOutput = TRUE' to see C++ compilation details.
## compilation finished.
```

```
Runtime <- system.time(myNimbleOutput <- runMCMC(mcmc = cMCMC,
    nburnin = nburnin, niter = niter, nchains = nchains, samplesAsCodaMCMC = TRUE))
```

```
## running chain 1...

## |-------------|-------------|-------------|-------------|
## |-------------------------------------------------------|

## running chain 2...

## |-------------|-------------|-------------|-------------|
## |-------------------------------------------------------|

## running chain 3...

## |-------------|-------------|-------------|-------------|
## |-------------------------------------------------------|
```

#### V.PLOT OUTPUT

```
# N
for (t in 1:n.occasions) {
    PlotJagsParams(myNimbleOutput, params = paste("N[", t, "]",
        sep = ""), sim.values = Pop.Compo[[2]][t])
}
```

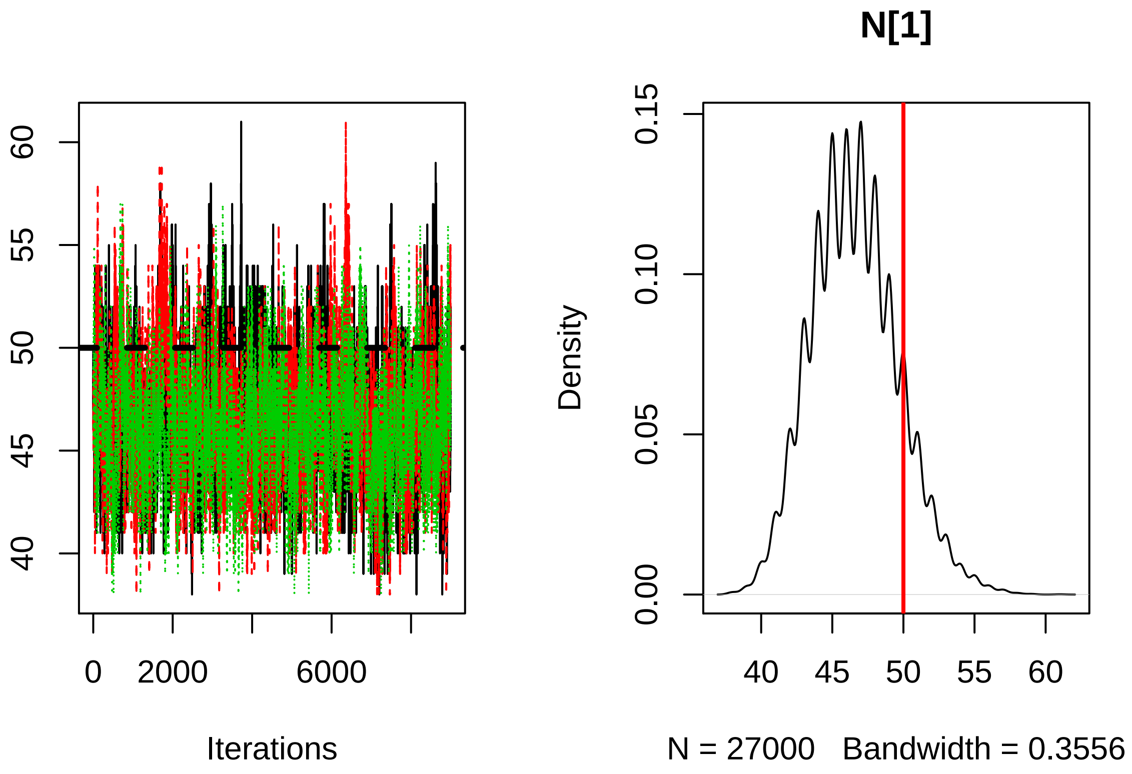

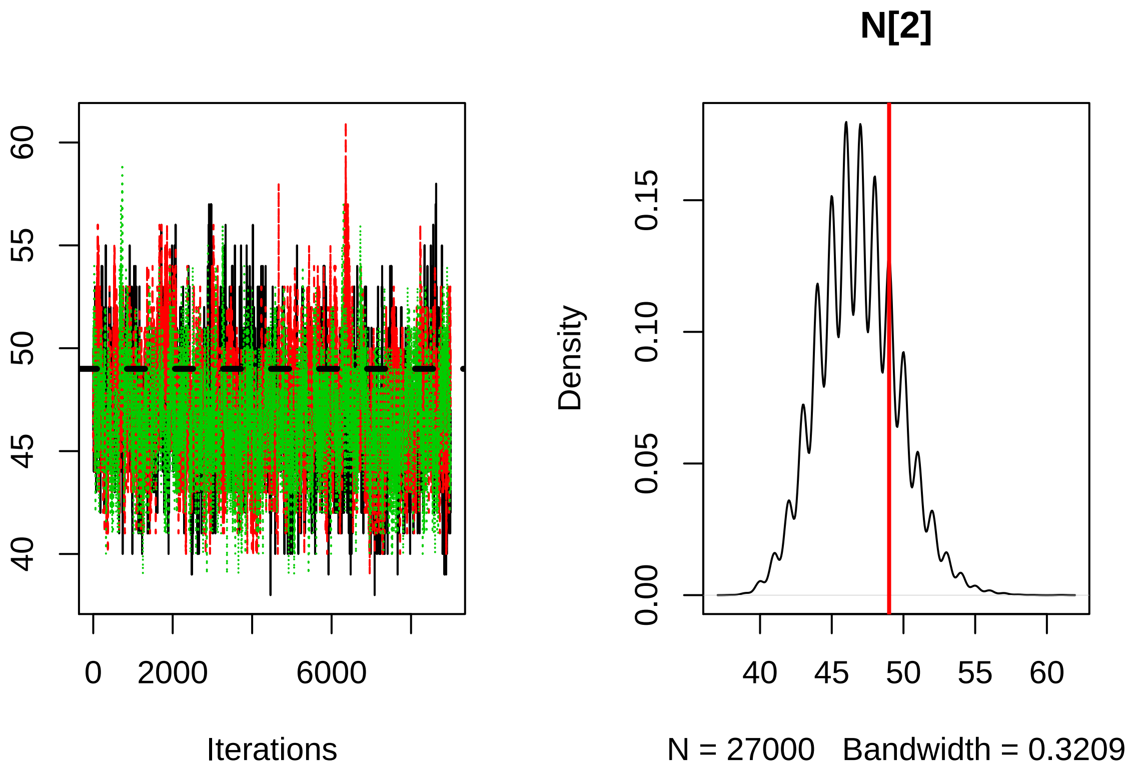
60
55
50
45
40
0
2000
6000
Iterations
N[2]
Density
0.15
0.10
0.05
0.00
40
45
50
55
60
N = 27000 Bandwidth = 0.3209

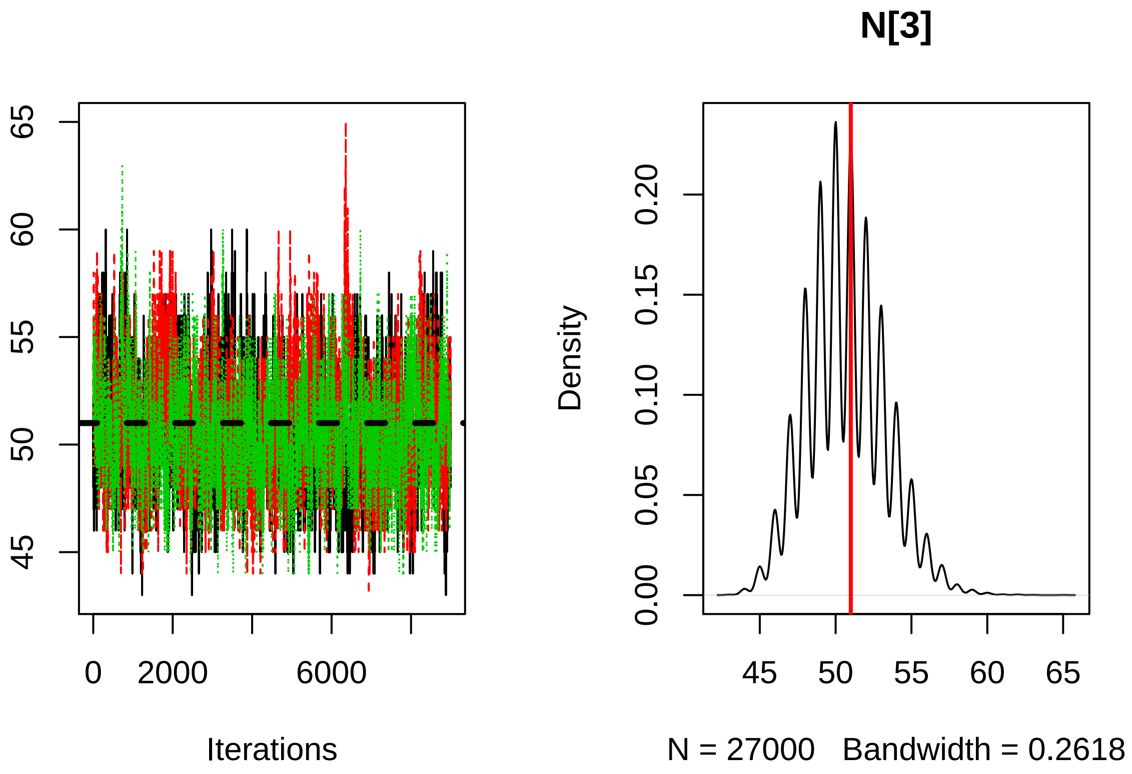
N[3]
65
60
55
50
45
0
2000
6000
Iterations
Density
0.20
0.15
0.10
0.05
0.00
45
50
55
60
65
N = 27000 Bandwidth = 0.2618

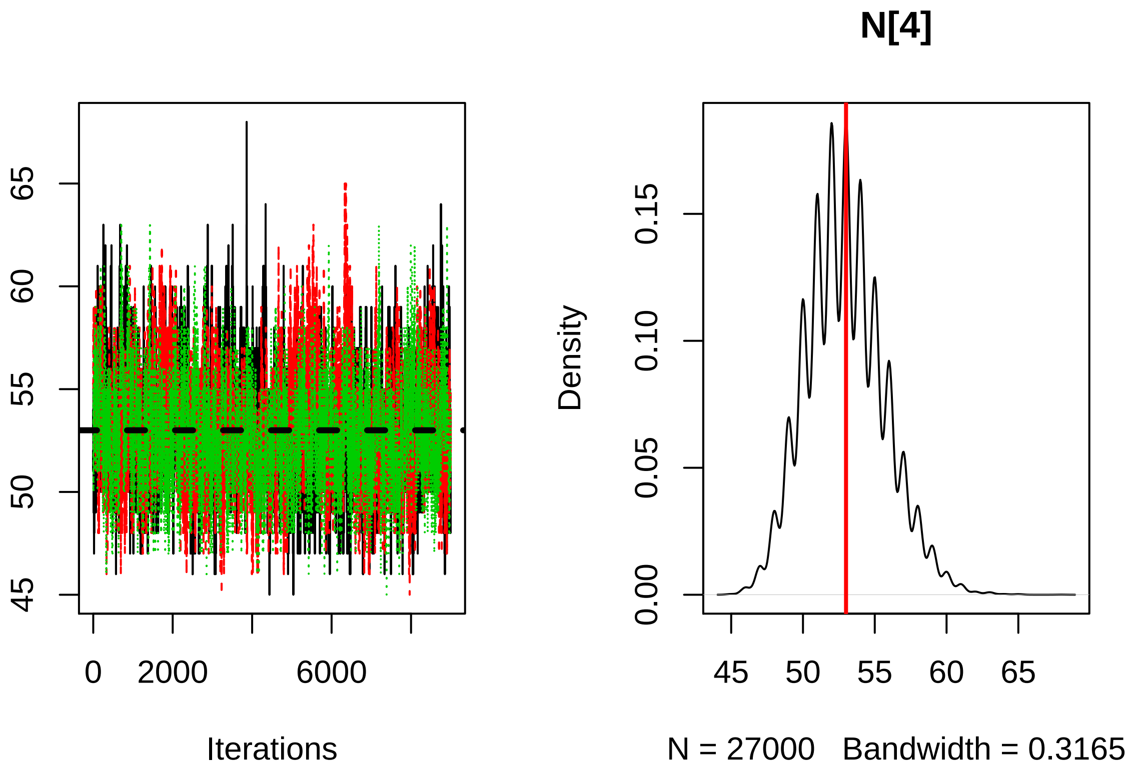
N[4]
45
50
55
60
65
0
2000
6000
Iterations
Density
0.00
0.05
0.10
0.15
45
50
55
60
65
N = 27000 Bandwidth = 0.3165

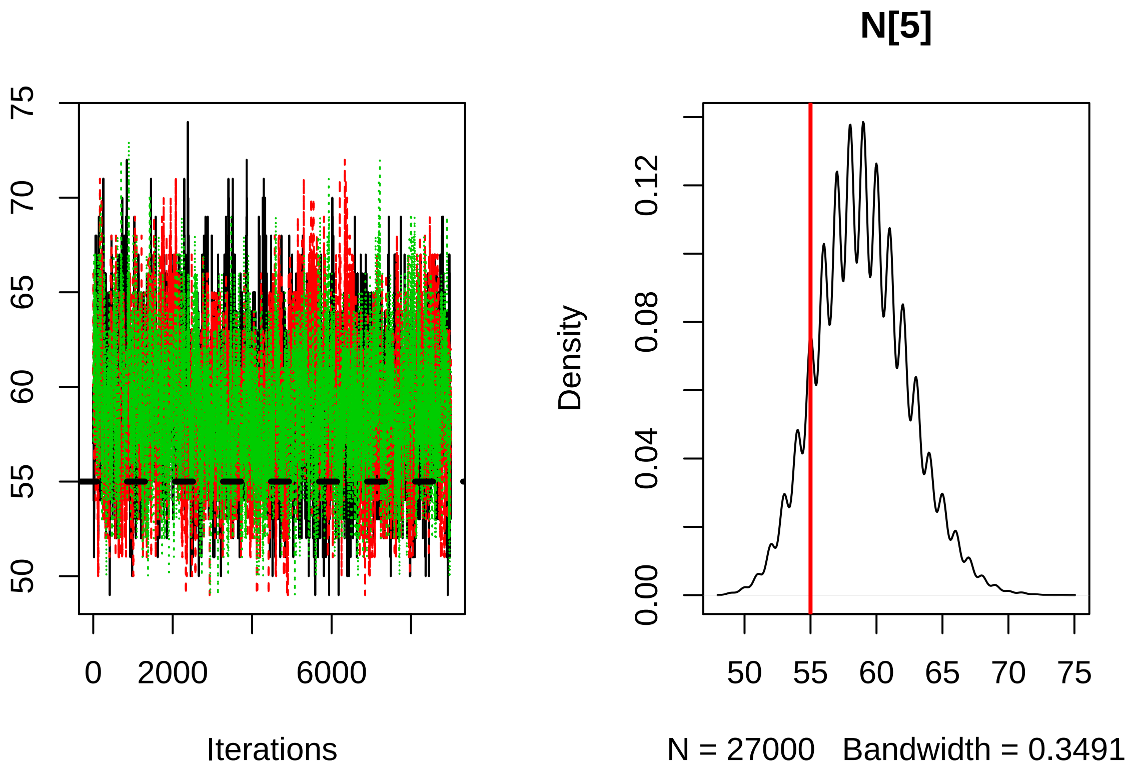


```r
# SIGMA
PlotJagsParams(myNimbleOutput, params = "sigma", sim.values = sigma)
```

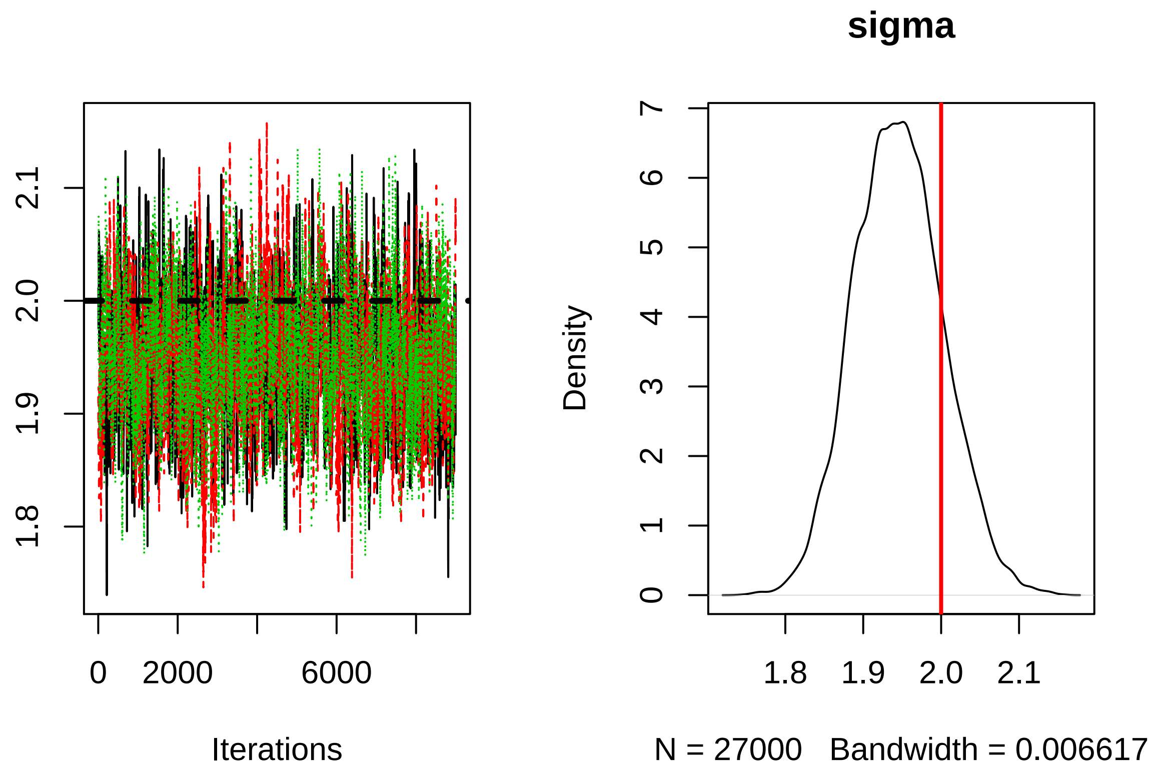


```r
# p0
PlotJagsParams(myNimbleOutput, params = "p0", sim.values = p0)
```

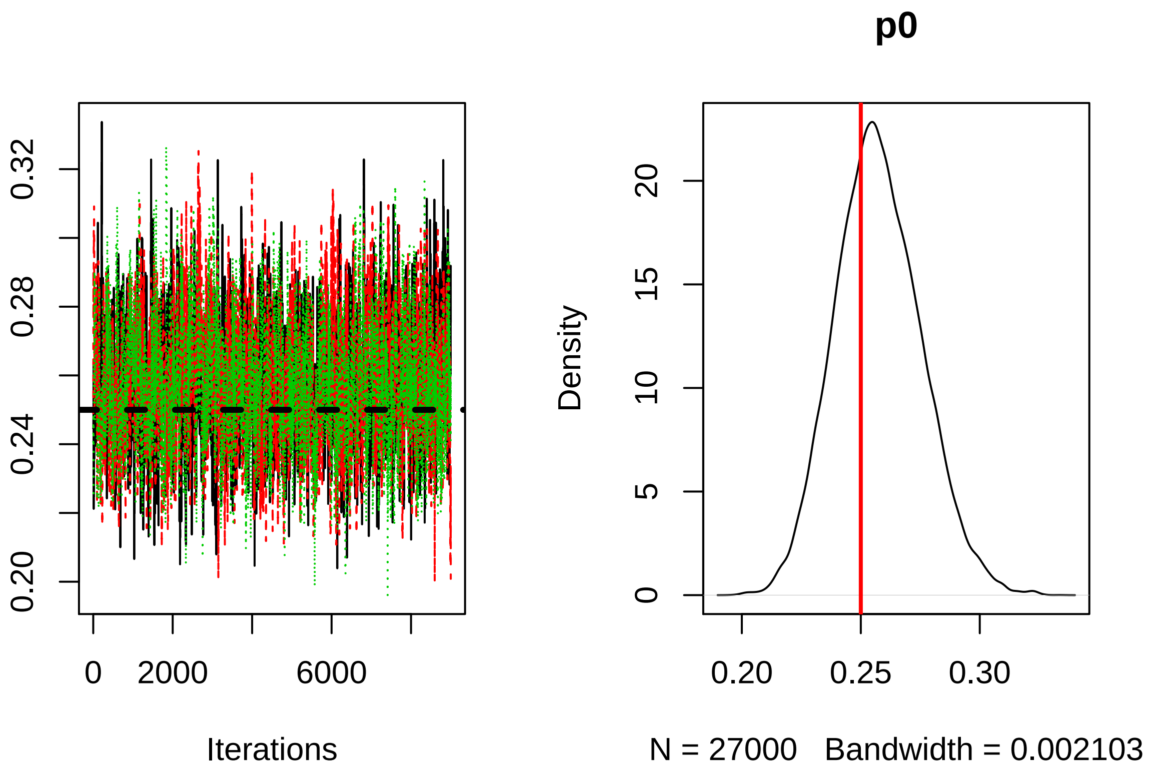


```r
# dispSigma
PlotJagsParams(myNimbleOutput, params = "dispSigma", sim.values = dispSigma)
```

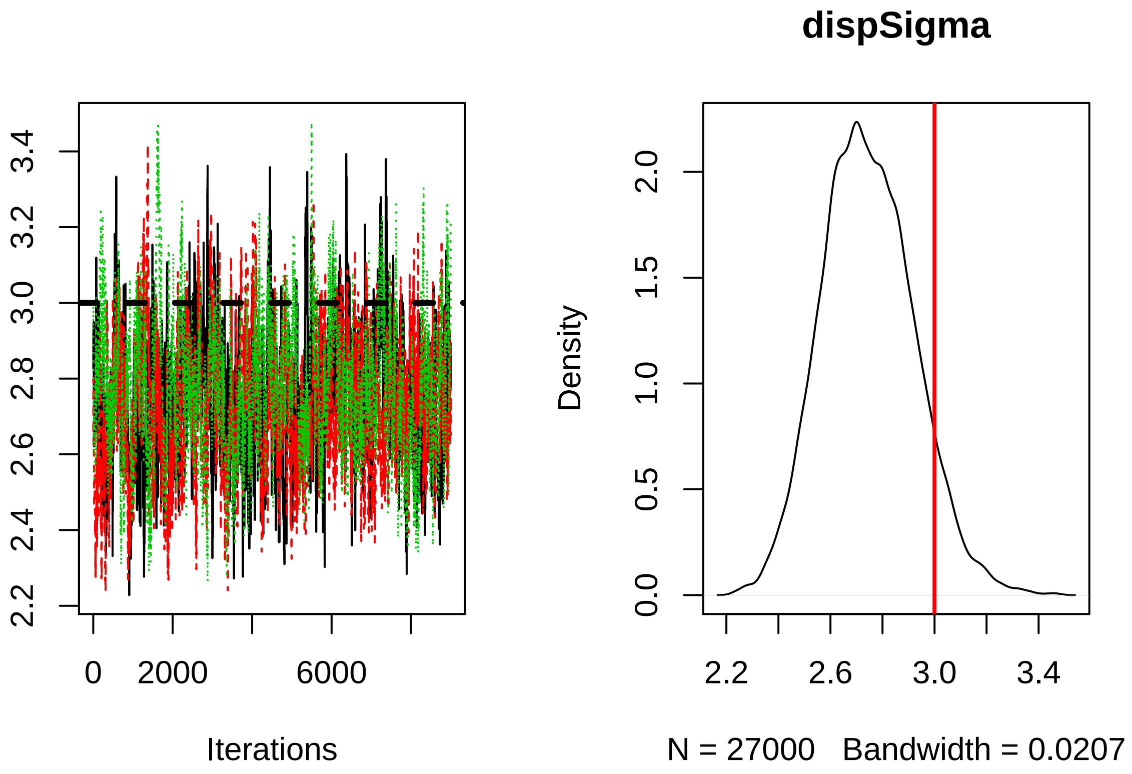


```r
# phi
PlotJagsParams(myNimbleOutput, params = "phi", sim.values = phi)
```

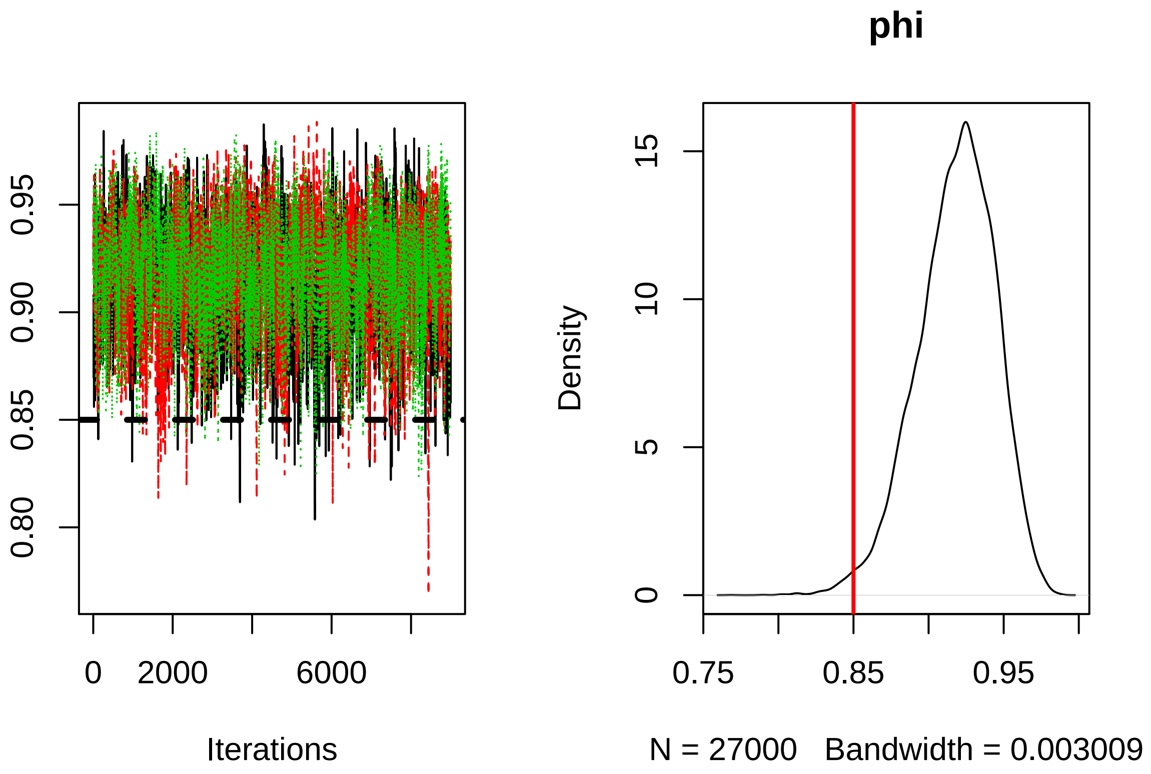


```
# rho
PlotJagsParams(myNimbleOutput, params = "rho", sim.values = rho)
```

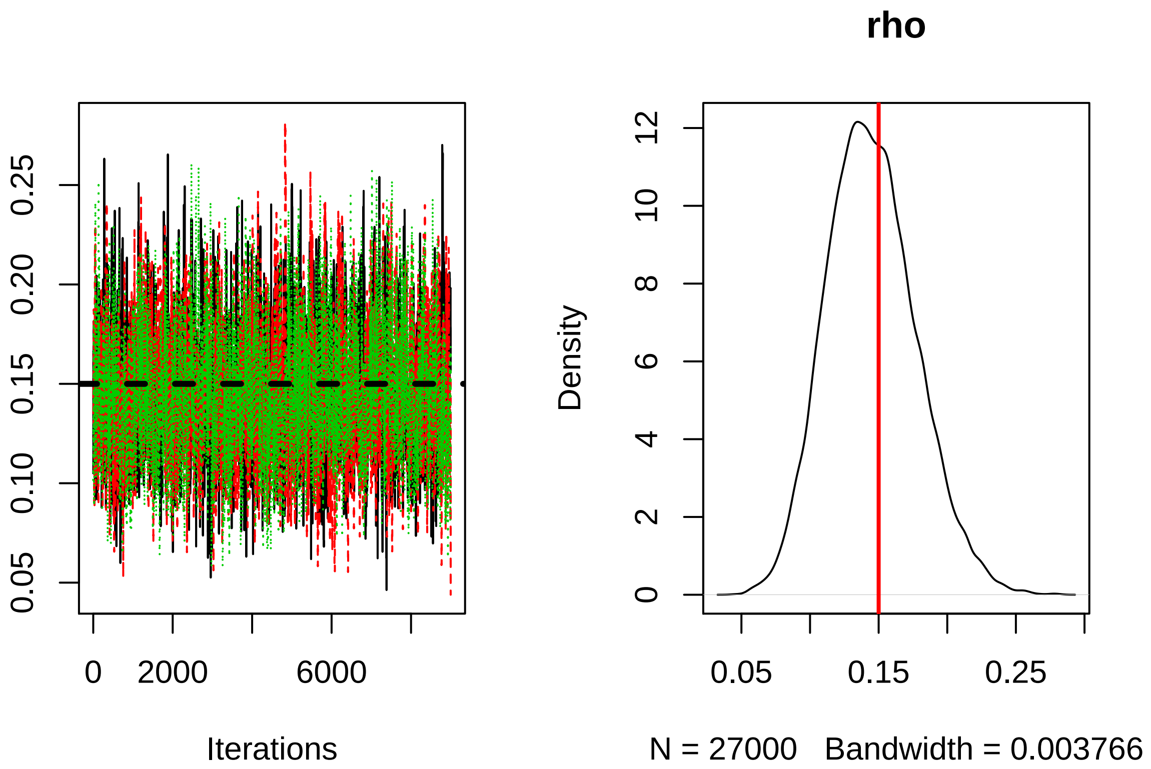


##References

Milleret, C., P. Dupont, C. Bonenfant, H. Broseth, O. Flagstad, C. Sutherland, and R. Bischof. 2019. "A Local Evaluation of the Individual State Space to Scale up Bayesian Spatial Capture Recapture." *Ecology and Evolution* 9 (1): 352–63. https://doi.org/10.1002/ece3.4751.

Milleret, C., P. Dupont, H. Broseth, J. Kindberg, J.A. Royle, and R. Bischof. 2018. "Using Partial Aggregation in Spatial Capture Recapture." *Methods in Ecology and Evolution* 9 (8): 1896–1907. https://doi.org/10.1111/2041-210X.13030.

NIMBLE Development Team. 2019. *NIMBLE: MCMC, Particle Filtering, and Programmable Hierarchical Modeling*. https://cran.r-project.org/package=nimble: https://cran.r-project.org/package=nimble. https://doi.org/http://doi.org/10.5281/zenodo.1211190.

Valpine, P. de, D. Turek, C.J. Paciorek, C. Anderson-Bergman, D.T. Lang, and R. Bodik. 2017. "Programming with Models: Writing Statistical Algorithms for General Model Structures with Nimble." *Journal of Computational and Graphical Statistics* 26 (2): 403–13.
